# Interpreting Omics Data Analysis with Large Language Models for Disease Target and Drug Discovery

**DOI:** 10.64898/2026.04.30.721768

**Authors:** Zixi Xu, Weihang Chen, Wuyu Ren, Tianqi Xu, Somadina Amaechina, Raad Khan, Yixin Chen, Michael Province, Philip Payne, Fuhai Li

## Abstract

In biomedical scientific discovery, synthesizing prior knowledge from the literature is an essential component of interpreting numerical omics data analyses for disease target identification and drug discovery. Large language models (LLMs) alone can rapidly retrieve disease mechanisms from biomedical text, but text-only outputs are general and unreliable for target and drug prioritization without cohort-specific quantitative evidence. Herein, we propose a provenance-aware Text-to-Target framework that couples schema-constrained multi-model LLM retrieval with numeric omics data analysis. The key design is a modality-aware fusion step: candidates are partitioned into overlap-supported anchors, retrieval-only hidden hubs, and network-emergent novelty nodes, then propagated into staged hypothesis and strategy generation under topology constraints. We evaluate the model in Alzheimer’s disease (AD) and pancreatic ductal adenocarcinoma (PDAC). In PDAC, the workflow produced a balanced 75-gene candidate universe and a 23-strategy portfolio, with significant DepMap support at both target level and strategy level. In AD, stricter candidate controls yielded a compact 34-gene universe and 14 strategies; under an expanded CRISPRbrain registry, both target-level axes were significant, with strong strategy-level enrichment. Across both diseases, final strategies preserved full provenance closure to the candidate pool, enabling end-to-end auditability from retrieval artifacts to validation outputs. These results support a transferable discovery architecture in which omics evidence constrains biological activity, LLM retrieval expands mechanistic search space, and network-aware fusion preserves interpretability. The framework provides a reproducible basis for dual-disease target prioritization and motivates continuous literature-mechanism concordance with agentic evidence-refresh loops.

## 1 Introduction

### 1.1 Motivation

Large-scale single-cell omics data can potentially characterize key disease dysfunctional signaling mechanisms for precision target and drug discovery. Although all human cells share a common genome, disease phenotypes emerge from context-dependent regulatory programs and multicellular interactions [1, 2]. In practice, the key bottleneck is not detecting expression shifts, but translating noisy and descriptive omics contrasts into mechanistically prioritized, testable targets and effective drugs.

In biomedical target and drug discovery, literature synthesis remains essential for translating numerical omics contrasts into mechanism-level interpretation [3–9]. Current precision-medicine workflows therefore split into two weakly connected stages. First, computational omics pipelines quantify disease-vs-reference shifts [10–14]. Second, experts manually interpret those outputs using literature synthesis and biological reasoning. This handoff is slow, difficult to standardize, and vulnerable to omission. LLMs improve literature-scale retrieval speed, but text-only reasoning is not sufficient for reliable prioritization. LLM outputs can hallucinate links, miss context-specific dependencies, and drift from cohort-level transcriptional evidence [15]. The core challenge is therefore integration: how to combine broad semantic priors from text with quantitative cohort constraints from omics while preserving interpretability and auditability.

In this study, we propose to address this challenge with a Text-to-Target framework that couples LLM retrieval, Single-cell-resolved differential signals, and PathFinder-weighted network inference in a single provenance-aware pipeline. The framework is designed to produce mechanism-grounded candidates that can be traced from retrieval and network evidence to final validation outcomes.

### 1.2 Single-cell Omics and LLM Priors

Our framework assigns explicit roles to each evidence modality. Single-cell or meta-cell (average of about 200 300 single cells) omics serves as the quantitative backbone: we start from integrated single-cell and single-nucleus atlases, then analyze Single-cell-level profiles to reduce sparsity and dropout while preserving biologically meaningful cell states [16]. This representation improves the stability of differential-expression estimation and downstream network inference.

LLMs serve as structured literature-retrieval engines rather than standalone decision makers. Schema-constrained prompts retrieve cell-type-specific targets, pathway context, and mechanism statements in machine-parseable tables. Prompt templates are disease-matched: PDAC prompts focus on malignant epithelial and tumor-microenvironment signaling, whereas AD prompts emphasize neuroimmune coupling, glial state transitions, and intercellular propagation logic.

This division of labor is the central design principle. Omics constrains what is active in the cohort, LLM retrieval expands mechanistic search space, and network-based fusion resolves their intersection and disagreement. Candidates retained after fusion carry explicit provenance tags, enabling transparent traceability from final strategy outputs back to both quantitative and text-derived evidence.

### 1.3 AD and PDAC Testbeds

We evaluate the framework in two complementary diseases that stress different aspects of target discovery. PDAC is a high-mortality solid tumor with limited long-term benefit from current multi-agent regimens and strong unmet need for mechanism-guided target prioritization [17–19]. AD is a complex neurodegenerative disease affecting millions of people in the United States, and no curative therapy is currently available because key molecular mechanisms remain incompletely resolved [20–24].

PDAC provides a high-contrast benchmark for lineage-matched analysis. Disease progression from normal acinar or ductal states to malignant epithelial states is shaped by recurrent oncogenic programs, including frequent *KRAS* activation, but therapeutic responses remain constrained by resistance and microenvironment complexity [25]. To support robust expression contrasts, we built a high-confidence acinar-lineage reference from HCA, GEO, and PDAC single-nucleus resources [11, 26–28]. Acinar-lineage cells in PDAC cohorts were annotated through a twostage process: lineage classification followed by disease-state stratification into normal-like and malignant-like subsets. This lineage-matched design reduces confounding from cell-identity imbalance and enables cleaner interpretation of malignant transformation-associated signaling. After consistent preprocessing and filtering, retained matrix dimensions differ modestly across groups by design, reflecting group-aware denoising rather than forced symmetry. AD provides a complementary low-contrast benchmark. Disease programs in AD are distributed across interacting neuronal and glial states, including astrocytes, microglia, oligodendroglial lineages, and vascular-associated compartments. Compared with PDAC, AD therefore requires tighter candidate-space control to preserve interpretability under weaker and more heterogeneous signal geometry. This dual-disease setup allows us to test both transferability and boundary conditions of the same pipeline. PDAC probes performance in a stronger malignant-vs-reference regime, whereas AD probes robustness under broader cell-state heterogeneity and diffuse effects.

### 1.4 Text-to-Target Overview

The Text-to-Target workflow is a staged integration cycle that alternates expansion and constraint. We use an ensemble of frontier LLMs (GPT-5, Gemini 2.5-pro, and DeepSeek-r1) to retrieve disease- and cell-context priors in schema-constrained form [29–32]. Single-cell differential-expression analysis then injects cohort-specific quantitative evidence, and PathFinder transforms these signals into weighted network structure.

Candidate fusion is performed through provenance-aware quadrants: *Q*1 (jointly supported anchors), *Q*2 (retrieval-only hidden hubs), and *Q*3 (network-emergent novelty). Hypothesis upgrading and multi-target strategy generation are then constrained by explicit topology metadata, and final outputs are evaluated with disease-matched perturbation resources (DepMap for PDAC, CRISPRbrain for AD). This design moves beyond direct list overlap and yields high-confidence targets supported by convergent text and omics evidence with explicit provenance closure. Our goal is not only stronger prioritization, but also an auditable discovery process in which each final strategy can be traced to concrete retrieval, network, and validation artifacts.

## 2 Methods

### 2.1 Workflow Overview

The Text-to-Target framework unifies semantic priors from biomedical text with quantitative evidence from disease-state omics while preserving the distinct epistemic role of each modality. Rather than treating the literature branch and expression branch as two lists to be overlapped, the workflow first converts transcriptomic contrast into a weighted signaling graph and then projects language-derived targets into this graph. This design separates jointly supported candidates from text-only and network-emergent candidates and retains provenance for each candidate class during downstream hypothesis generation and validation.

PathFinder is used as the primary omics integration engine that translates DEG evidence into topological context. Candidate construction is therefore not defined by direct LLM-DEG overlap alone. DEG-informed PathFinder networks provide weighted nodes and edges, and these are fused with curated LLM targets to form a quadrant candidate pool that supports both conservative and novelty-oriented discovery.

The full pipeline can therefore be read as a gradual information-constraining process. Literature retrieval starts broad and mechanistic, expression analysis constrains this space with disease-state measurements, network propagation organizes measured signals into pathway structure, quadrant construction labels candidates by evidence provenance, and final hypothesis/therapy generation is conditioned on this structured candidate pool rather than on an untyped merged list. The same provenance-aware design also supports disease-matched downstream validation at both single-target and multi-target strategy resolution, implemented in DepMap for PDAC and CRISPRbrain for AD.

### 2.2 Data Curation and Harmonization

Expression inputs were retrieved from the OmniCellTOSG ecosystem through CellTOSG Loader workflows [6]. For PDAC, we focused on a lineage-matched contrast by selecting malignant epithelial meta-cells as the disease state and acinar-lineage meta-cells as the reference state. Loader-level query constraints preserved key meta-data fields needed for traceability and were configured to reduce avoidable cohort imbalance across major descriptors when feasible.

For AD, we used the same loading and harmonization principles but instantiated disease-control contrasts across brain cell-state contexts relevant to neurodegeneration. In downstream hypothesis generation, a glia-centric emphasis (astrocyte and microglial disease programs with linked neuronal context) was prioritized to reduce diffuseness of whole-brain candidate space and improve mechanistic specificity of cross-cell signaling hypotheses.

Raw matrices were harmonized before inferential analysis. Gene symbols were standardized to a common symbol space, non-numeric artifacts were coerced and audited, duplicate symbols were collapsed at gene level, and genes with zero aggregate expression within a group were removed. This harmonization avoids inflation of low-information features and improves comparability across the disease and reference matrices. Because filtering is group aware, final matrices are comparable but not forced to have identical gene counts; this preserves biologically meaningful asymmetry after denoising.

The resulting PDAC matrices are summarized in Table 1. Disease-specific AD matrices were processed under the same normalization and filtering logic and then passed to AD PathFinder and candidate-pooling modules. These harmonized matrices provide the numerical substrate for differential-expression statistics and graph-based signaling inference in both disease branches.

**Table 1:** Summary of processed PDAC and AD expression matrices used for down-stream analysis.

| Dataset | Cell population | Genes | Samples | Matrix shape |
| --- | --- | --- | --- | --- |
| PDAC | Acinar (normal-like) | 17,250 | 201 | $17,250 \times 201$ |
| PDAC | Malignant (cancer-like) | 18,790 | 214 | $18,790 \times 214$ |
| AD | Astrocyte (Disease) | 41,149 | 176 | $41,149 \times 176$ |
| AD | Astrocyte (Normal) | 41,149 | 176 | $41,149 \times 176$ |
| AD | Microglia (Disease) | 41,149 | 139 | $41,149 \times 139$ |
| AD | Microglia (Normal) | 41,149 | 139 | $41,149 \times 139$ |
| AD | Glutamatergic Neuron (Disease) | 41,149 | 128 | $41,149 \times 128$ |
| AD | Glutamatergic Neuron (Normal) | 41,149 | 128 | $41,149 \times 128$ |

**Table 2:** Stage-wise cardinalities.

| Pipeline layer | Count |
| --- | --- |
| Curated upstream prior ($Q1 \cup Q2$) | 49 |
| Network novelty reservoir ($Q3$) | 26 |
| Integrated candidate universe ($Q1 \cup Q2 \cup Q3$) | 75 |
| Upgraded mechanism coverage | 19 |
| Final strategy portfolio size | 23 |
| Final strategy target diversity | 23 |

### 2.3 Differential Expression Priors

For each retained gene *g*, group means over malignant and acinar meta-cells were calculated and summarized by

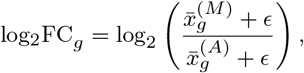

where *ϵ* is a small stabilization constant. Statistical significance was tested using two-sided Welch *t*-tests, followed by Benjamini-Hochberg correction across genes. Genes satisfying |log_2_ *FC*_*g*_ | *>* 1 and FDR_*g*_ *<* 0.05 were treated as differential-expression signals.

In this framework, DEG signals are not consumed directly as a final target list. Instead, they serve as quantitative priors for network learning in PathFinder. This distinction is important: the objective is not only to identify strongly shifted genes, but also to infer which parts of the signaling graph organize these shifts into coherent disease-associated routes. The same inferential principle was applied to AD with disease-vs-control contrasts defined in a neurodegeneration setting, while allowing disease-specific downstream filtering rules in candidate pooling.

### 2.4 LLM Retrieval and Curation

The text branch uses multiple LLM families as independent retrieval engines to reduce model-specific idiosyncrasy. Retrieval is implemented as a chained prompt workflow with strict schema constraints. The process begins by eliciting a disease-specific cell-type and subpopulation map at atlas-compatible granularity, with subpopulations grouped into biologically coherent compartments and accompanied by concise functional descriptions. This taxonomy output is then partitioned into chunks so that subsequent retrieval can be executed with bounded context per branch while preserving full coverage of major compartments in each disease setting.

For each cell-type chunk, the model is asked to retrieve three linked target classes in disease context: therapeutic targets, surface markers, and key driver genes. The output format is fixed as a structured table containing cell type, standardized target symbol, target class, short disease-function text, and evidence citation fields. Prompt constraints explicitly require literature support and disallow unsupported entries. Citation reporting is normalized to either PMID or author-year style so that each row can be audited independently. In a subsequent linked pass, the retrieved target table is used as input for pathway mapping, where targets are assigned to standardized pathway resources (for example KEGG, Reactome, or Hallmark-style nomenclature) with associated cell type, key member targets, biological effect, and citation fields.

Prompting is therefore schema-oriented rather than free-form, and each phase produces machine-parseable markdown tables instead of narrative prose. Branch-level outputs are merged, deduplicated, and normalized to HGNC-style symbols; unresolved or malformed entities are removed from the discovery layer. The final normalized text-derived set is denoted *T*_LLM_. Each retained target keeps provenance attributes including model family, prompt lineage, cell-type annotation, target class, and linked citation metadata, so that downstream integration and hypothesis generation remain fully traceable to retrieval evidence. In AD instantiation, additional emphasis was placed on astrocyte-microglia-neuron interaction logic and state-transition language so that retrieval outputs better support intercellular mechanism construction rather than only intracellular cascade descriptions.

### 2.5 PathFinder Network Inference

PathFinder [11] is applied as the core network-inference engine in both disease branches. For PDAC, malignant epithelial meta-cells were used as case state and acinar cells as reference state. For AD, disease-control contrasts were instantiated over neurodegeneration-relevant cellular states and then integrated into a unified weighted intra-network view. In both settings, PathFinder operates on a prior biological interaction graph and learns path importance through graph-transformer supervision.

Let *G* = (*V, E*) denote the prior graph and 𝒫 = {*p*_*k*_} the candidate signaling paths. In our implementation, path length is constrained to biologically plausible ranges (up to 10 hops) to avoid diffuse routes that are hard to interpret mechanistically. Node features are derived from expression profiles over meta-cells and encoded through graph-aware transformer layers; the original PathFinder configuration is preserved with six transformer layers, eight attention heads, and hidden dimension 16. Optimization follows the established training setup (learning rate 1×10^−5^, L2 weight decay 3 × 10^−7^, gradient clipping at norm 5.0, batch size 4, 25 epochs, fixed seed).

Path-level importance parameters *α*_*k*_ are learned jointly with phenotype classification. To incorporate expression contrast as a soft biological prior, each path is assigned

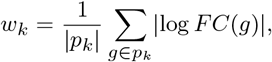

and training minimizes

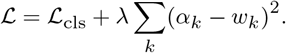

The regularization term encourages consistency between learned path salience and DEG-derived signal magnitude while preserving predictive discrimination through ℒ_cls_. Importantly, the exported node weight is an integrated PathFinder importance score learned under DEG-informed constraints, rather than a direct one-to-one remapping of a single differential-expression statistic. As a result, the network weights retain quantitative linkage to disease-vs-reference transcriptomic contrast while also encoding pathway-level topological context learned during model fitting.

After fitting, high-importance paths are aggregated to produce graph artifacts in JSON format (full network and reduced-radius subnetworks). Exported nodes carry importance weights and exported edges carry relation weights. These weighted artifacts are not auxiliary visualizations; they are the primary omics representation consumed during candidate construction.

For AD specifically, we retained three cell-branch PathFinder exports and built an integrated weighted graph using observed cell-distribution proportions (glutamatergic : microglial : astrocyte = 128 : 139 : 176). Because branch topologies were identical while weights differed, weighted integration preserved shared structure and produced a single graph for downstream quadrant construction and bridge analysis.

### 2.6 Quadrant Candidate Construction

Candidate construction proceeds by integrating *T*_LLM_ with the set of genes represented in the PathFinder export, denoted *V*_PF_. The integration is written as

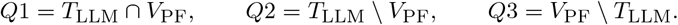

The meaning of each subset is biologically and methodologically distinct. Genes in *Q*1 are anchors jointly supported by semantic priors and network activation. Genes in *Q*2 retain text-prior support but are absent from the current network realization, so they are preserved as potentially context-conditional or underrepresented candidates rather than removed. Genes in *Q*3 are network-emergent novelty nodes that are not directly retrieved by the LLM branch.

For *Q*1, each entry carries both network and text metadata, including PathFinder node weight, LLM cell-type annotation, and functional class. For *Q*2, curation emphasizes entity validity and interpretable class assignment so these candidates can be incorporated explicitly during mechanism expansion. For *Q*3, ranking does not rely solely on node weight; it also quantifies bridge behavior relative to *Q*1 anchors. Given a novelty node *u* ∈ *Q*3, we compute two bridge descriptors from network connectivity patterns: the number of anchor pairs linked through *u* (reported as `q1_pair_count`) and the number of unique anchors participating in those links (reported as `q1_unique_count`). We further preserve relation-level direction class (directed versus undirected-only) and compact path representations in detail tables. This produces a novelty ranking that favors nodes that are both weight-supported and topologically relevant to known anchor structure.

In AD, direct transfer of PDAC-style broad *Q*2 construction produced an overly large retrieval-only set relative to downstream interpretability. We therefore constrained *Q*2 through a stricter refinement trajectory. A connectivity-aware core was first extracted by requiring non-overlap retrieval targets to remain linked to *Q*1 anchors through pathway-supported reachability, prioritizing directed support where available and using undirected support as a controlled fallback. Because this strict core under-covered several biologically central AD programs, we then applied a capped layered supplement that prioritized targets recurring in astrocyte-, microglial-, and glutamatergic-relevant pathway contexts and recurring across merged retrieval outputs. This preserved interpretability while avoiding broad re-expansion of retrieval-only noise.

Likewise, AD *Q*3 construction was based on explicit threshold profiling rather than a single hard cutoff. Raw novelty candidates were assembled from *Q*1 first/second-neighborhood nodes and intermediate nodes on *Q*1-*Q*1 bridge paths. Each candidate was scored by PathFinder node weight, direct anchor-neighbor support, and bridge frequency. We compared strict, balanced, and loose regimes over these metrics and retained the balanced regime (high-weight with non-zero bridge support) as the final AD setting, because it preserved bridge explainability and yielded a tractable novelty set under weaker effect concentration than in PDAC.

The final candidate universe used for hypothesis generation is

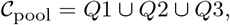

with explicit provenance tags retained for all elements. This provenance tagging enables every downstream hypothesis or therapy proposal to be traced back to its evidentiary origin.

### 2.7 Hypothesis and Strategy Generation

Mechanistic generation is conditioned on provenance-labeled candidates rather than on a flat merged list, and is executed through a progressive prompt chain aligned with the quadrant design. The initial mechanism pass receives *Q*1 anchors together with *Q*2 hidden-hub candidates and asks the model to assemble candidate signaling networks without introducing therapy content at this stage. Outputs are constrained to a structured mechanism table that records hypothesis identifiers, directed network sequences, and role decomposition into upstream nodes, core nodes, and downstream nodes, plus short biological rationale fields.

After this baseline mechanism layer is established, the model is supplied with *Q*3 novelty-node details and bridge-path constraints derived from the network artifacts. At this point, prompt instructions explicitly require consistency with provided topology metadata, including bridge direction class and path length. The model is asked to insert *Q*3 nodes at biologically plausible positions within existing signal flow and to treat longer bridge paths as potentially indirect links requiring mechanistic interpretation of intermediate processes. Updated hypotheses are again returned in structured table format so that pre- and post-novelty mechanism versions can be compared directly.

Only after upgraded mechanism tables are finalized does the workflow transition to strategy design. Therapy generation is constrained to select target combinations directly from the upgraded network sequences, typically using compact 2-4 node sets, and to provide a rationale for combinational logic and expected synergistic effect. Prompt constraints favor existing or repurposable drug classes when available, while preserving explicit linkage between each strategy and its parent mechanism hypothesis. Because provenance is retained throughout the chain, final strategy rows can be audited as intersection-supported, text-prior-supported, or network-emergent-supported, which is essential for transparent prioritization and downstream statistical validation.

For AD, the same staged prompting template was retained but with disease-tailored emphasis. Prompts de-emphasized purely tumor-style linear intracellular chains and instead encouraged mechanistic narratives centered on glial state transitions, neuroinflammatory feed-forward loops, and astrocyte-microglia-neuron coupling. This adjustment was important because AD candidates are distributed across multiple interacting cell states; forcing a rigid linear style reduced biological plausibility and strategy traceability in pilot runs.

### 2.8 Disease-Matched Validation

Because available perturbation resources differ between oncology and neurodegeneration settings, validation was implemented as two disease-specific modules under a shared permutation-testing logic.

For PDAC, validation is conducted in DepMap using two complementary statistical views that address different levels of evidence. At target resolution, PDAC model identifiers are derived from `Model.csv` using pancreas/PDAC matching over Oncotree metadata fields. For each gene *g*, we compute

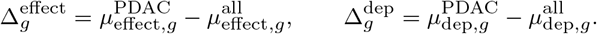

For a target set *S*, summary statistics are set means over Δ^effect^ and Δ^dep^. Significance is estimated by same-size permutation against a shared gene universe (default 5000 draws). Effect is evaluated with a left-tail empirical test (more negative indicates stronger PDAC-selective vulnerability), and dependency is evaluated with a right-tail empirical test (more positive indicates stronger PDAC-selective dependency), both using (+1)*/*(*N* + 1) correction.

At PDAC strategy resolution, each therapy row is treated as one multi-gene strategy unit *s* with gene set *G*_*s*_. We compute

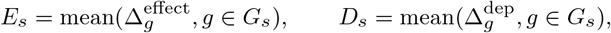

then map to directional standardized scores

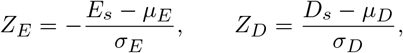

and combine them as

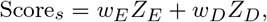

with default *w*_*E*_ = *w*_*D*_ = 0.5. Local significance is estimated via same-size permutation for each strategy, while global significance is estimated by permutation over the full strategy collection with observed strategy-size profile preserved.

For AD, validation is conducted in CRISPRbrain using AD-labeled versus nonAD-labeled screen registries. Screen-level gene scores are standardized within screen (default `abs_z` transform) and aggregated across included screens. For each gene *g*, we compute

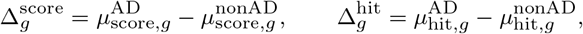

where hit is derived from screen hit-class annotations (with threshold fallback when needed). Target-level significance is then assessed by same-size right-tail permutation for set means of Δ^score^ and Δ^hit^.

At AD strategy resolution, each strategy *s* with gene set *G*_*s*_ is summarized by

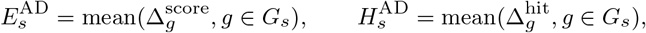

followed by global-universe standardization and composite scoring:

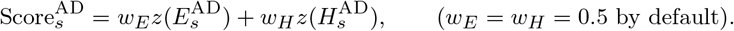

As in PDAC, local same-size permutation and global profile-preserving permutation are both reported.

Together, these validation modules separate broad target-pool enrichment from combination-level actionability while preserving disease-appropriate perturbation semantics.

### 2.9 Drug-enabled Network Construction and Interactive Evidence Visualization

To make downstream outputs directly usable for therapeutic triage, we introduced a drug-enabled concordance layer that augments the mechanism backbone with evidence-linked compounds. The design preserves the same provenance-aware principle used throughout the framework: signaling routes remain the primary mechanistic object, while drug associations are attached as explicit evidence records rather than replacing route-level interpretation.

For each disease branch, construction integrates three synchronized inputs: (i) a curated Mermaid signaling topology (core target-target routes), (ii) a deduplicated disease-specific target-drug evidence ledger, and (iii) PathFinder-derived target weights from the corresponding weighted network export. Target nodes retain their signaling-role classes (for example overlap-supported, literature-emergent, LLM-supported, or contradicted) and inherit quantitative size scaling from PathFinder support. Because these PathFinder weights are learned under DEG-informed priors, the rendered node sizes remain anchored to disease-state transcriptomic contrast while preserving network-level context. When gene-level *p*-value tables are available, the same rendering interface can optionally provide a sensitivity-view using − log_10_(*p*) scaling without changing edge semantics.

Drug nodes are generated by matching ledger targets to topology aliases, followed by deduplication at the target-drug-citation level. Each target-to-drug edge stores explicit evidence attributes, including evidence direction (support/mixed/contradict), development status (approved/investigational/preclinical/unknown), citation text, and hypothesis linkage fields. This schema keeps therapeutic interpretation auditable at edge resolution, avoiding loss of evidence polarity in flat target summaries.

The resulting graph bundle is exported in machine- and human-readable formats: structured JSON for downstream analysis, interactive HTML for node/edge inspection, and publication-oriented static renders. The interactive layer exposes target-level network evidence and drug-level therapeutic metadata in hover fields, allowing direct tracing from signaling context to literature-backed drug support.

In this study, we intentionally evaluated two evidence-input regimes. The core outputs (PDAC core and AD core) were seeded by manual literature-review tests before graph construction. PDAC core used one manually curated resistance-focused benchmark study [33]. AD core used four manually curated mechanistic exemplars spanning cGAS-STING neuroinflammation, ABCA7/NLRP3 glial signaling, CD74/MIF immune-metabolic modulation, and hippocampal glucose rescue [34–37].

We then stress-tested an automated expansion mode in PDAC. The extended PDAC output used LLM-assisted retrieval to aggregate a larger, continuously updateable literature pool into standardized node-evidence-edge-node records, and then merged that evidence graph with prior LLM mechanism hypotheses plus the deduplicated drug evidence ledger. This manual-seed versus automated-expansion contrast is retained as an explicit provenance axis in downstream result interpretation.

### 2.10 Reproducibility and Audit Trail

The full workflow is script-driven, seed-controlled, and artifact-oriented. Intermediate outputs are persisted in machine-readable formats at each integration boundary, including normalized LLM retrieval tables, PathFinder weighted JSON networks, quadrant tables (Q1/Q2/Q3), bridge-detail summaries, structured hypothesis and therapy tables, and disease-specific validation reports (DepMap for PDAC; CRISPRbrain registries and score summaries for AD) at both target and strategy levels.

This artifact lineage design supports independent re-analysis under alternative thresholds, model settings, or ranking criteria without rerunning the full pipeline from scratch. It also ensures that any final claim can be traced backward to the exact computational objects that generated it, which is particularly important in multi-module pipelines where evidence from language models, networks, and perturbation screens is fused.

## 3 Results

We report results in the same order as the pipeline: retrieval quality, provenance-aware candidate fusion, topology-constrained hypothesis expansion, and disease-matched perturbation validation. This organization separates stage-wise soundness checks from final actionability evidence and makes cross-disease comparisons explicit.

### 3.1 Retrieval Quality and Upstream Prior

The prompt-chained retrieval workflow generated a structured upstream prior that could be handed to the network-integration stage without additional manual rewriting. After normalization and curation, the retrieval-facing pool (defined as *Q*1 ∪ *Q*2) contained 49 unique targets, each represented in table-form outputs that preserved biologically interpretable fields rather than free-text fragments. This structure is important for reproducibility: each target remains linked to a specific retrieval lineage and can be followed downstream into candidate construction, hypothesis expansion, and final strategy validation.

At this stage, we focused on soundness checks that test whether the retrieval layer is computationally usable before asking whether it is therapeutically useful. The key checkpoint was symbol-level interoperability with the downstream PathFinder node space. The curated retrieval pool could be projected into the network fusion step in a deterministic manner, indicating that this prior was sufficiently clean for provenance-aware integration rather than ad hoc post-processing.

A second checkpoint examined whether the retrieval layer was overly narrow. Instead of collapsing onto only one PDAC signaling axis, the curated set retained both canonical malignant regulators and context-linked components that become relevant when hypotheses are later expanded under bridge constraints. This balance prevented the candidate pipeline from becoming a simple confirmation exercise and created room for network-emergent novelty in later stages.

### 3.2 Candidate Pool Composition

Integrating the curated retrieval pool with PathFinder-derived network nodes yielded a 75-target candidate universe that remained balanced across provenance classes: 27 overlap-supported anchors in *Q*1, 22 retrieval-only hidden hubs in *Q*2, and 26 network-emergent novelty nodes in *Q*3. In relative terms, these correspond to 36.0%, 29.3%, and 34.7% of the final candidate universe, respectively. This near-balanced composition indicates that neither modality dominated candidate construction.

The biological profile of each quadrant also aligned with its intended role. The overlap quadrant (*Q*1) contained highly weighted disease-relevant anchors such as *MYC* (13871.35), *STAT3* (9115.33), and *EGFR* (2280.81), indicating convergence between text-derived priors and network-activated structure. The retrieval-only quadrant (*Q*2) retained plausible but currently non-projected entities (for example *KRAS, IL6, TBK1*, and *AURKA*), preserving potentially context-dependent hypotheses rather than discarding them prematurely. The network-only quadrant (*Q*3) provided a dedicated novelty reservoir that was not reducible to simple retrieval overlap.

From a stage-wise reliability perspective, this candidate construction result is central to the method logic. The fused set is explicitly topology-aware and provenance-labeled, enabling downstream mechanism generation to reason over anchor certainty, hidden priors, and emergent bridges as distinct evidence types.

**Table 3:** Quadrant composition of the integrated candidate universe.

| Quadrant | Count | Share | Representative nodes |
| --- | --- | --- | --- |
| Q1 (LLM ∩ PathFinder) | 27 | 36.0% | MYC, STAT3, EGFR, VIM, TGFBR2 |
| Q2 (LLM-only hubs) | 22 | 29.3% | KRAS, IL6, TBK1, AURKA, ZEB1 |
| Q3 (PathFinder-only) | 26 | 34.7% | MAX, EGR1, CTCF, ELK1, MAZ |

### 3.3 Q3 Bridge Topology

To test whether *Q*3 novelty nodes were mechanistically informative rather than peripheral artifacts, we quantified bridge statistics against *Q*1 anchors. Across all *Q*3 targets, the median `q1_pair_count` was 18.5 (range 3–94), and the median `q1_unique_count` was 15.5 (range 4–42). These values indicate that many novelty nodes participate in multiple anchor-anchor relationship patterns and therefore occupy topologically central, not merely terminal, positions in the inferred network.

The highest-ranked examples reinforced this interpretation. *MAX* combined the largest weight in *Q*3 (10415.66) with broad anchor bridging (63 pairs, 41 unique anchors), while *MAZ* achieved the strongest pair-level bridge signal (94 pairs, 42 unique anchors). *EGR1, CTCF*, and *ELK1* similarly showed mixed weight-topology support. In the context of hypothesis upgrading, these nodes are valuable because they provide concrete insertion points where network-informed novelty can be introduced into previously anchor-centered mechanistic narratives.

A practical implication is that novelty integration can be constrained rather than speculative. Because each *Q*3 candidate is accompanied by bridge statistics, downstream prompts can be instructed to prioritize nodes with high joint support (weight plus bridge connectivity), which reduces the risk of arbitrary expansion.

**Table 4:** Top *Q*3 novelty nodes prioritized by weight and bridge connectivity.

| Q3 node | Node weight | Representative top anchor hubs |
| --- | --- | --- |
| MAX | 10415.66 | APEX1, RARA, KIT |
| EGR1 | 9442.34 | CXCR6, EGFR, SDC1 |
| CTCF | 7118.87 | IL2RA, ADRB2, STAT3 |
| ELK1 | 7101.45 | ADRB2, IL2RA, EGFR |
| MAZ | 6323.25 | KIT, NCR1, VIM |

### 3.4 Hypothesis Upgrading and Traceability

Mechanism generation began with *Q*1 + *Q*2 and was then upgraded by introducing *Q*3 bridge constraints. This sequence changed the role of novelty from optional decoration to structured mechanism expansion. The upgraded corpus referenced 19 unique `Upgraded-Hypo-*` identifiers and yielded 23 final strategy rows, indicating that expanded mechanism space was translated into concrete design outputs rather than remaining purely descriptive.

A critical stage-wise soundness criterion at this boundary is provenance closure: final strategy targets should remain within the integrated candidate universe. This criterion was fully satisfied. All 23 strategy targets were members of *Q*1 ∪ *Q*2 ∪ *Q*3, with a provenance mix of 12 from *Q*1 (52.2%), 2 from *Q*2 (8.7%), and 9 from *Q*3 (39.1%). The resulting pattern is informative. The strategy layer remained anchored to overlap-supported biology, but still incorporated substantial network-emergent novelty, consistent with the intended role of the upgraded hypothesis stage.

This balance suggests that the upgraded prompts did not over-privilege either conservative anchors or novelty nodes. Instead, provenance classes were translated into complementary mechanism roles, allowing strategies to remain interpretable while expanding beyond direct overlap candidates.

**Table 5:** Provenance composition of targets used in finalized strategies.

| Source quadrant | Targets used | Share |
| --- | --- | --- |
| $Q1$ anchors | 12 | 52.2% |
| $Q2$ hidden hubs | 2 | 8.7% |
| $Q3$ novelty bridges | 9 | 39.1% |

### 3.5 Strategy Portfolio Structure

The final therapy corpus produced 23 valid strategies from 23 parsed rows, so no candidate strategy was lost during structural parsing. Strategies were contributed by all three generation blocks (9, 7, and 7 rows), reducing sensitivity to any single prompt branch and improving internal robustness.

Strategy size distribution indicated a deliberately compact design space: 12 two-target strategies (52.2%), 10 three-target strategies (43.5%), and one four-target strategy (4.3%), with an average of 2.52 targets per strategy. This profile is useful for translation because it balances mechanistic coverage with practical combinational tractability.

Target recurrence patterns were coherent but not degenerate. *MYC* appeared in 8 strategies, while *EGFR* and *STAT3* each appeared in 7, and *AXL* appeared in 4. This indicates a consistent mechanistic center around receptor-transcription and stress-adaptation axes, yet without complete collapse into one repeated combination. Additional recurrence among *MAX* and *IL6* (three each) further supports a multi-axis architecture linking proliferative control, stromal signaling, and transcriptional adaptation.

Another stage-wise checkpoint asked whether statistically stronger strategies were concentrated in only one generation block. They were not. Locally significant strategies (right-tail *p <* 0.05) were distributed across all blocks (4, 4, and 5), supporting the conclusion that the upgraded hypothesis layer produced broadly usable mechanism families rather than one dominant template.

**Table 6:** Therapy portfolio structure and recurrence profile.

| Composition statistic | Value |
| --- | --- |
| Total valid strategies | 23 |
| Source blocks (Table-1 / Table-2 / Table-3) | 9 / 7 / 7 |
| Strategy size distribution (2 / 3 / 4 targets) | 12 / 10 / 1 |
| Average targets per strategy | 2.52 |
| Local significant strategies by block ($p < 0.05$) | 4 / 4 / 5 |
| Most frequent target | MYC (8 strategies) |
| Next recurrent targets | EGFR (7), STAT3 (7) |

### 3.6 PDAC DepMap Validation

DepMap validation was performed on the finalized 23-target strategy universe across 69 PDAC models, and evaluated separately at target and strategy levels. This two-resolution design is important because it distinguishes general pool quality from combination-level therapeutic prioritization.

At target resolution, the effect metric showed a significant left shift relative to random same-size sets (left-tail empirical *p* = 0.0039992, *z* = −3.2523), while the dependency metric showed a significant right shift (right-tail empirical *p* = 0.0361928, *z* = 2.0780). The directionality is consistent with the intended interpretation of stronger PDAC-selective vulnerability and dependency in the curated target set.

At strategy resolution, enrichment was stronger and more decisive. The observed mean strategy score was 1.1400, compared with a permutation mean near zero (−0.0021), yielding *z* = 8.9883 and right-tail *p* = 0.00019996. Local testing identified 13 significant strategies out of 23 (56.5%), demonstrating that signal concentration was not limited to only a few outliers.

Top-ranked strategies were enriched for receptor-transcription couplings and dual-axis interventions, led by `T2_R1` (EGFR+MYC), `T1_R1` (ERBB2+EGFR+MYC), and `T3_R4` (AXL+MYC). At target directionality level, the strongest negative effect deltas were seen for *MYC, PPARG, CTCF*, and *MAX*, while strongest positive dependency deltas included *PPARG, ERBB2, EGFR*, and *MAX*. These directional leaders are concordant with the recurrent mechanism cores observed in the strategy portfolio.

### 3.7 PDAC Synthesis

Taken together, the PDAC pipeline produced a coherent evidence trajectory from structured retrieval to network fusion, from provenance-constrained hypothesis expansion to strategy-level functional validation. Each stage passed a local soundness checkpoint before feeding the next stage: retrieval outputs were computationally interoperable, candidate fusion remained modality-balanced, novelty nodes showed measurable bridge centrality, upgraded hypotheses preserved provenance closure, and final strategies showed both structural diversity and concentrated validation signal.

**Table 7:** Condensed DepMap validation summary.

| Validation level | Statistic | Result |
| --- | --- | --- |
| Target-level (effect) | Left-tail empirical $p$ | 0.0039992 |
| Target-level (effect) | Directional $z$-score | -3.2523 |
| Target-level (dependency) | Right-tail empirical $p$ | 0.0361928 |
| Target-level (dependency) | Directional $z$-score | 2.0780 |
| Strategy-level (global) | Right-tail empirical $p$ | 0.00019996 |
| Strategy-level (global) | Global $z$-score | 8.9883 |
| Strategy-level (local) | Significant strategies ($p < 0.05$) | 13/23 (56.5%) |

**Table 8:** Top strategies by strategy-level score.

| Strategy ID | Target set | Score | Local $p$<br>(right-tail) |
| --- | --- | --- | --- |
| T2_R1 | EGFR, MYC | 3.523 | 0.00340 |
| T1_R1 | ERBB2, EGFR, MYC | 3.142 | 0.00240 |
| T3_R4 | AXL, MYC | 2.741 | 0.00980 |
| T1_R8 | PPARG, MAP2K1, MYC | 2.540 | 0.00520 |
| T3_R1 | EGFR, EGR1, MYC | 2.320 | 0.00600 |

**Table 9:** Leading target-level directional signals from DepMap summaries.

| Signal type | Top targets |
| --- | --- |
| Most negative effect deltas | MYC, PPARG, CTCF, MAX, ERBB2, EGFR, AXL, HSP90AA1 |
| Most positive dependency deltas | PPARG, ERBB2, EGFR, MAX, MYC, HSP90AA1, AXL, CD4 |

These outcomes support the central methodological claim for the PDAC branch: PathFinder is most effective when treated as an analysis engine that reshapes candidate construction, rather than as a downstream verification add-on. Under this design, LLM retrieval contributes broad mechanistic priors, network inference contributes topology-aware evidence organization, and validation serves as a quantitative stress test of the jointly constructed strategy space.

### 3.8 AD Candidate Construction

Applying the same backbone workflow to AD produced a distinct but internally coherent signal profile. The merged AD retrieval layer contained 151 unique targets after deduplication and audit. PathFinder network artifacts for glutamatergic, microglial, and astrocyte branches were topology-consistent and were integrated into a weighted intra-network representation. Unlike PDAC, where direct fusion already produced a well-sized candidate universe, AD required additional control of candidate-space expansion to preserve interpretability under a weaker and more distributed disease signal.

The stage-wise contraction pattern makes this explicit. A broad retrieval-only AD pool remained large after initial pathway-aware projection, but strict connectivity filtering reduced it to a compact core that was then selectively expanded by a capped layered supplement. In parallel, novelty extraction started from a large bridge-candidate reservoir and was reduced by balanced topology constraints before final selection. This is reflected in final quadrant cardinalities: the AD branch produced a compact candidate universe of 34 genes, composed of 10 overlap-supported anchors, 14 retrieval-prior hidden hubs, and 10 network-emergent novelty nodes. No cross-quadrant overlap was retained in the finalized lists, preserving strict provenance separation. In AD context, this compactness is a feature rather than a deficit because it limits combinational explosion in downstream hypothesis and strategy generation.

**Table 10:** AD stage-wise cardinalities.

| Pipeline layer | Count |
| --- | --- |
| Merged AD retrieval targets (deduplicated) | 151 |
| Integrated AD PathFinder graph size | 469 nodes / 467 edges |
| Broad AD $Q_2$ (pre-refinement) | 106 |
| Strict AD $Q_2$ core | 4 |
| $Q_1$ anchors | 10 |
| $Q_2$ hidden hubs (strict+layered) | 14 |
| $Q_3$ novelty nodes (balanced) | 10 |
| Final AD candidate universe ($Q_1 \cup Q_2 \cup Q_3$) | 34 |
| Final AD strategy rows | 14 |
| Unique AD strategy targets | 13 |

### 3.9 AD Novelty Bridge Structure

The AD novelty pool behaved differently from PDAC in both scale and topology. Bridge-candidate extraction initially produced 134 potential *Q*3 nodes. Threshold profiling over strict, balanced, and loose regimes showed that strict filtering over-pruned mechanistically informative candidates, while loose filtering diluted interpretability with low-support tails. The balanced regime selected a final set of 10 nodes and preserved a high-signal novelty layer that remained manageable during hypothesis upgrading.

Relative to PDAC, AD novelty nodes showed lower bridge density overall, consistent with broader but weaker coupling structure in neurodegenerative state transitions. Within the selected AD *Q*3 set, bridge-count median was 5 (range 1–13) and direct anchor-neighbor median was 0 (range 0–1), indicating that many novelty nodes contribute through intermediate bridges rather than immediate anchor adjacency. Even under this sparser topology, the selected nodes remained mechanistically useful because they formed reproducible bridge motifs linking inflammatory and stress-response anchors.

### 3.10 AD Hypothesis-to-Strategy Translation

The AD hypothesis layer produced a compact but diverse strategy portfolio. Fourteen strategy rows were retained after parsing, covering 13 unique targets and referencing 11 unique `Upgraded-Hypo-*` identifiers. This indicates that AD hypotheses did not collapse into a single repetitive template despite tighter candidate-space control.

**Table 11:** AD selected novelty nodes and bridge statistics (balanced Q3).

| Q3 node | Node weight | bridge_count | q1_neighbor_count |
| --- | --- | --- | --- |
| MYC | 10169.20 | 1 | 0 |
| MAX | 8133.49 | 13 | 1 |
| EGR1 | 7623.48 | 5 | 1 |
| ELK1 | 4839.19 | 1 | 0 |
| TP53 | 2796.30 | 5 | 1 |

Provenance closure was fully maintained: every strategy target belonged to the finalized AD candidate universe, with target-source composition of 6 from *Q*1, 1 from *Q*2, and 6 from *Q*3. Thus, AD strategies remained anchored in overlap-supported biology while still incorporating network-emergent novelty. Recurrence patterns were biologically coherent, with *STAT3* as the dominant recurrent node (7 strategies), followed by *C3* and *PPARG* (5 each), and *EGR1* /*TP53* (4 each), consistent with an inflammatory-transcriptional core in the generated AD mechanism set.

**Table 12:** AD strategy architecture summary.

| Composition statistic | Value |
| --- | --- |
| Valid AD strategies | 14 |
| Unique AD strategy targets | 13 |
| Referenced upgraded hypothesis IDs | 11 |
| Target provenance mix (Q1 / Q2 / Q3) | 6 / 1 / 6 |
| Strategy size distribution (2 / 3 / 4 targets) | 5 / 6 / 3 |
| Most recurrent targets | STAT3 (7), C3 (5) |

### 3.11 AD CRISPRbrain Validation

AD validation was performed using CRISPRbrain with two registry settings: a default astrocyte ITC-focused profile and an expanded profile that additionally included neuron PSAP-KO contrasts and microglial immune-activation context. The expanded setting is treated as the primary AD validation view because it recovers full target coverage and better reflects AD-relevant cellular heterogeneity.

Under the expanded registry, target-level tests were significant on both axes (`delta_score` right-tail *p* = 0.0346, *z* = 1.9152; `delta_hit` right-tail *p* = 0.0402, *z* = 1.7247), while strategy-level global enrichment remained strongly significant (*p* = 0.00019996, *z* = 6.4864), with 7 of 14 strategies locally significant. Compared with default, expanded coverage increased from 10/13 to 13/13 targets. As expected, effect-size concentration on `delta_score` decreased when heterogeneity increased, while hit-consistency became stronger and remained significant.

At strategy resolution, the leading combinations were centered on an inflammatory-transcriptional axis and remained coherent with the upgraded hypothesis lineage. Top-ranked entries included EPHA4+STAT3 (`AD_THERAPY_014`), PPARG+STAT3 (`AD_THERAPY_012`), and STAT3+MYC (`AD_THERAPY_010`), each with local right-tail significance below 0.01. The concentration of high-scoring strategies around STAT3-linked combinations is consistent with recurrence patterns observed in the AD strategy table and supports a non-random, mechanism-consistent prioritization signal despite attenuated target-level effect concentration relative to PDAC.

**Table 13:** AD CRISPRbrain validation summary: default vs expanded registry.

| Metric | Default | Expanded |
| --- | --- | --- |
| AD / nonAD screens | 3 / 3 | 9 / 9 |
| Usable targets / total | 10 / 13 | 13 / 13 |
| Target `delta_score` right-tail $p$ | 0.0062 | 0.0346 |
| Target `delta_hit` right-tail $p$ | 0.2879 | 0.0402 |
| Strategy global right-tail $p$ | 0.0002 | 0.0002 |
| Strategy global $z$ | 5.6778 | 6.4864 |
| Local significant strategies | 7 / 13 | 7 / 14 |

### 3.12 Drug-enabled Network Outputs and Evidence Traceability

To evaluate whether the framework can move beyond target prioritization toward therapeutic triage, we generated three drug-enabled signaling outputs under two evidence regimes: PDAC core (manual-review seed), PDAC extended (automated retrieval expansion), and AD core (manual-review seed). All regenerated outputs remained topologically consistent with their mechanism backbones while adding explicit target-to-drug evidence edges. PDAC core contained 11 target nodes, 11 drug nodes, and 11 target-to-drug edges; PDAC extended expanded to 29 targets, 15 drugs, and 16 target-to-drug edges; and AD core contained 18 targets, 17 drugs, and 17 target-to-drug edges.

Evidence polarity was preserved at edge resolution rather than collapsed into binary support labels. In extended PDAC, target-to-drug edges were distributed as 12 support, 2 mixed, and 2 contradict links, while AD contained 8 support, 7 mixed, and 2 contradict links. This direction-aware representation is important for translational triage because it separates convergence-supported candidates from context-dependent or conflicting evidence patterns.

Layout diagnostics indicated that interactive outputs remained visually tractable. Estimated target-to-drug edge crossings were 0 in PDAC core, 2 in PDAC extended, and 0 in AD core despite increased extended-edge density. Representative renders are shown in Figures 2, 3, and 4. Together, these results establish an operational text-to-target-to-drug bridge in which signaling structure, quantitative weighting, and literature evidence are jointly inspectable in a provenance-preserving artifact, while also exposing a practical migration path from manual evidence seeding to automated literature expansion.

**Fig. 1:**
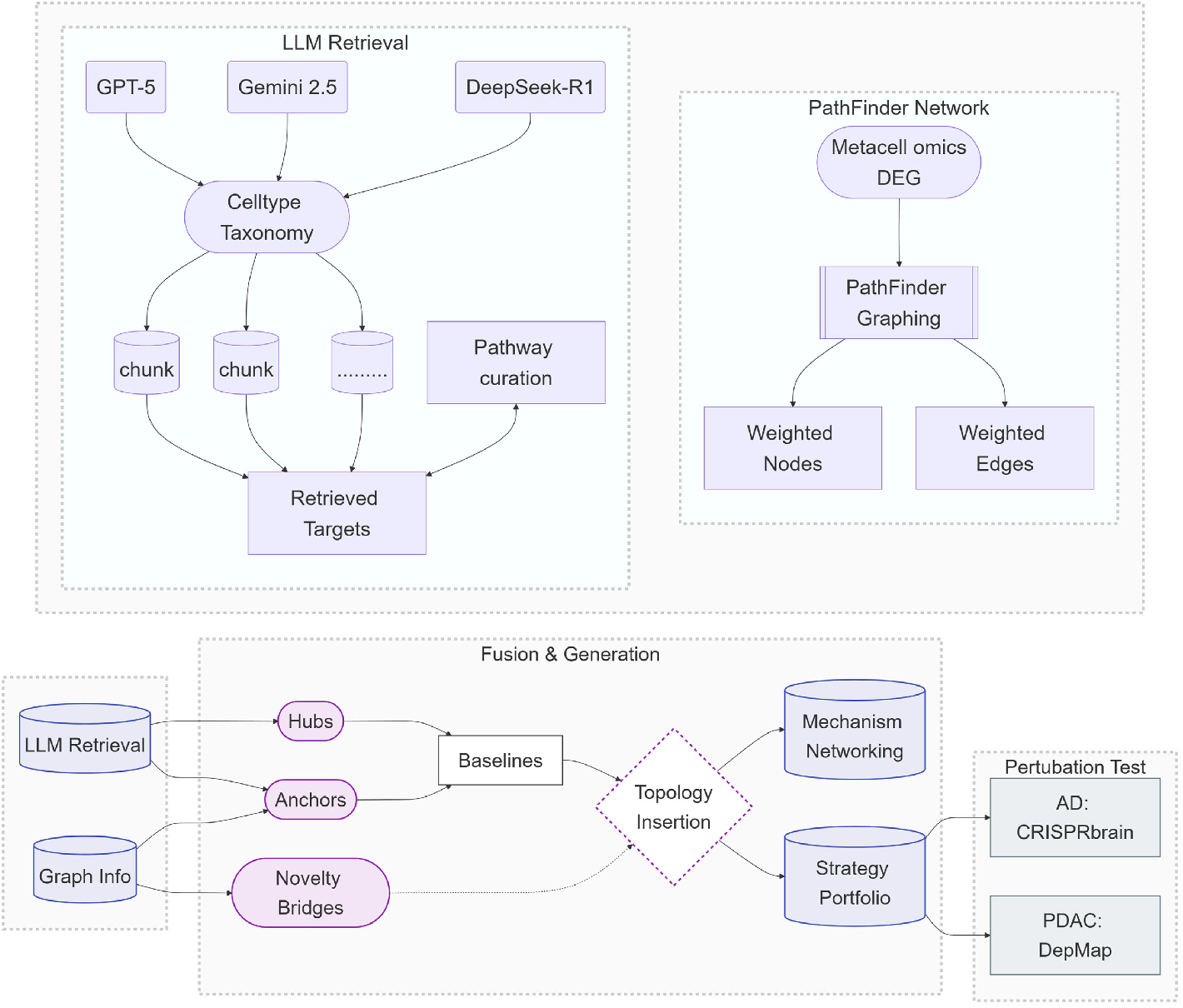
Overview of the LLM-Omics provenance-aware target discovery framework. The workflow integrates semantic priors with cohort-level omics evidence across three decoupled phases. **(A) Dual-Modality Extraction:** The language branch (LLM Engine) constructs a structural prior (*T*_LLM_) via schema-constrained chunked retrieval and pathway curation. In parallel, the omics branch (Omics Engine) transforms differential expression signals into a weighted biological network (*V*_PF_) using PathFinder as an inferential analysis engine. **(B) Fusion and Generation:** Bypassing simple list intersection, the framework partitions candidates into distinct provenance quadrants: dual-supported anchors (Q1), retrieval-only hidden hubs (Q2), and network-emergent novelty bridges (Q3). During hypothesis generation, the LLM first assembles baseline mechanistic sequences using Q1 and Q2, and subsequently upgrades them by inserting Q3 nodes under explicit network topology constraints, yielding a structured strategy portfolio. **(C) Disease-Matched Validation:** The generated multi-target strategies are evaluated through permutation testing using DepMap for the PDAC cohort and CRISPRbrain registries for the AD cohort, ensuring independent functional support for the prioritized combinations.

**Fig. 2:**
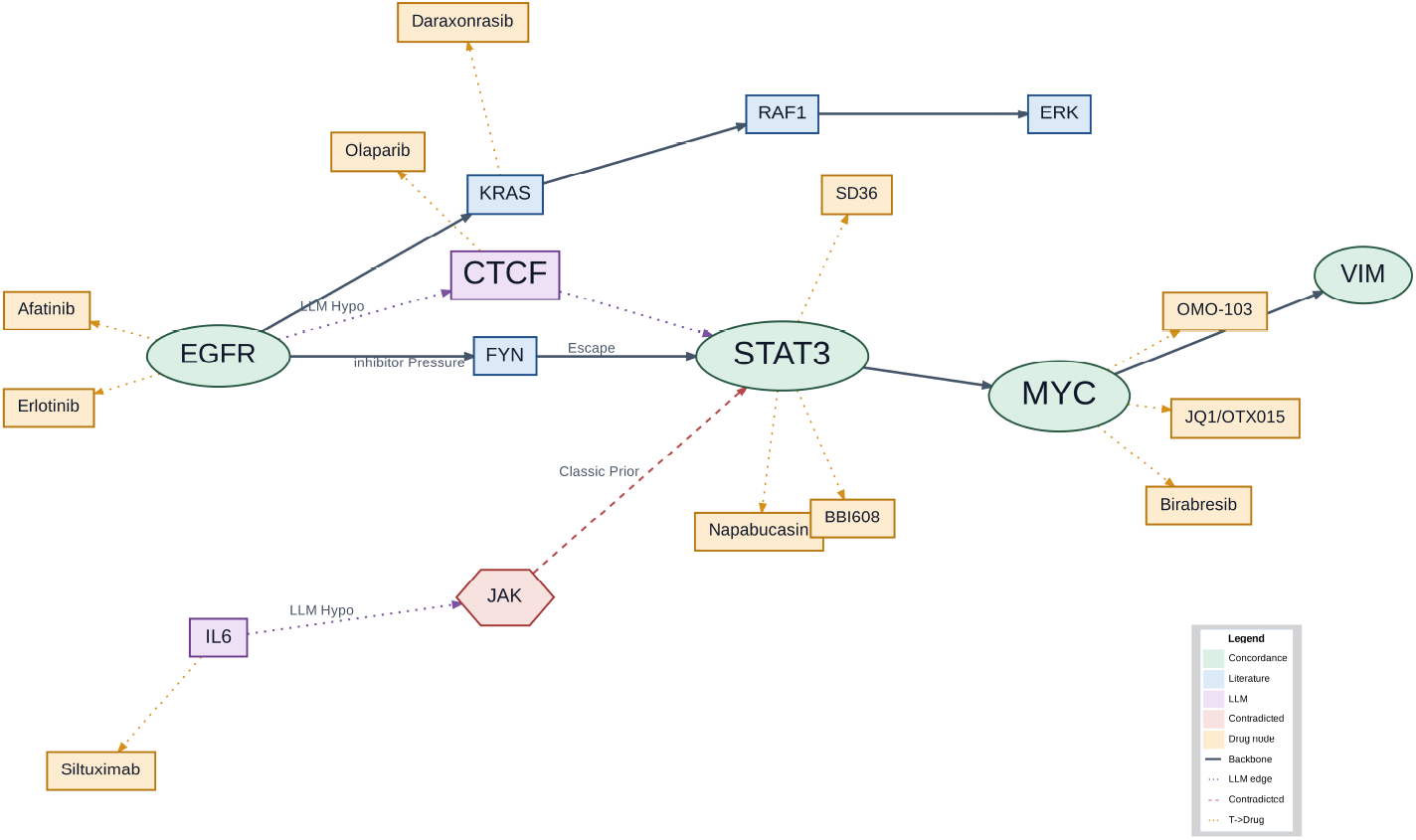
PDAC core drug-enabled concordance network (manual-review-seeded baseline). This baseline output was constructed from manually curated literature-seed evidence. Target node size follows quantitative network support, and target-to-drug links preserve direction-aware literature evidence and development-stage metadata.

**Fig. 3:**
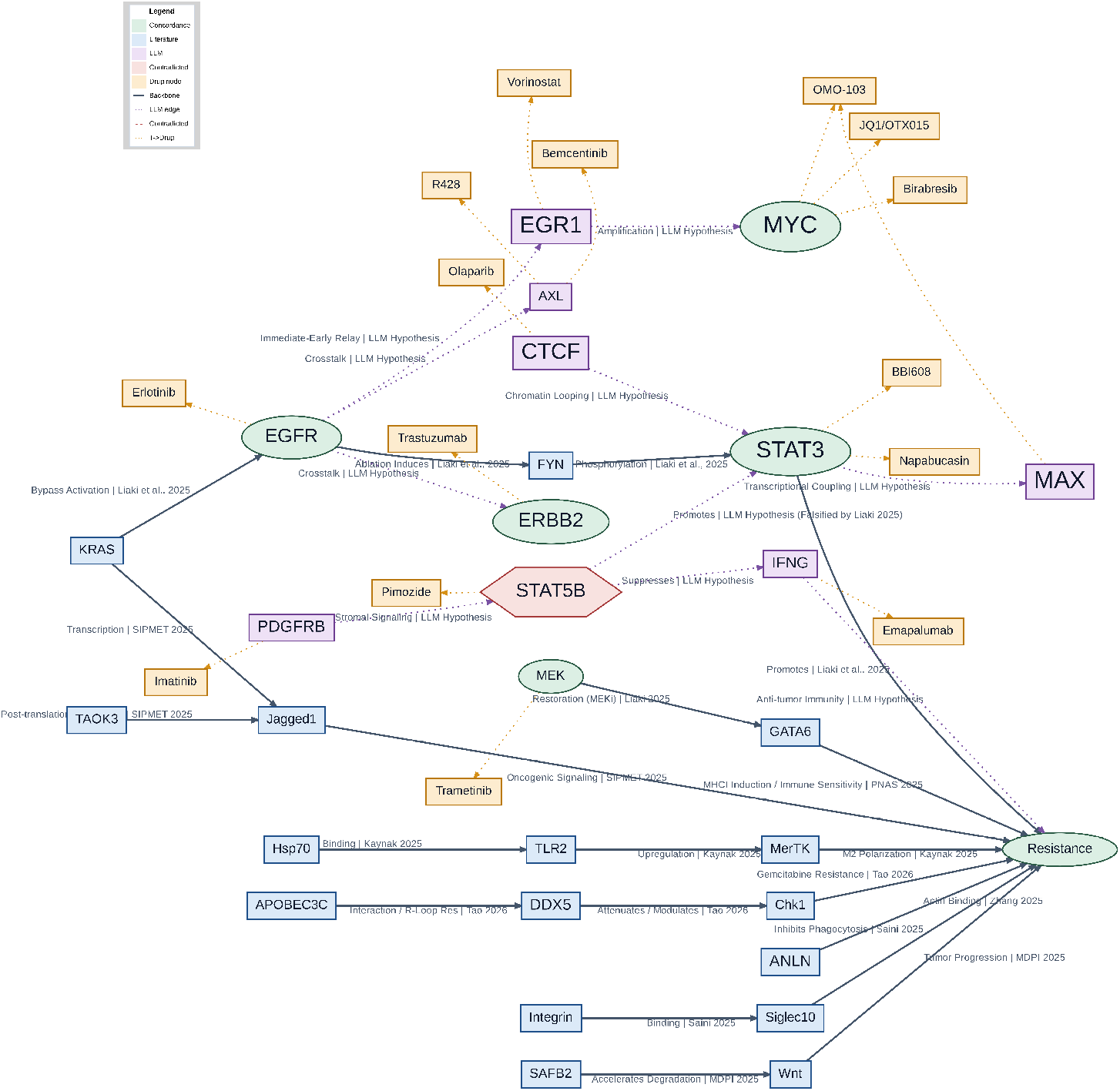
PDAC extended drug-enabled concordance network (LLM-assisted retrieval expansion). The extended output integrates a larger automatically retrieved literature pool normalized into node-evidence-edge-node records and merged with prior mechanism hypotheses plus the deduplicated drug ledger. Target node size follows quantitative network support, and target-to-drug links preserve direction-aware literature evidence and development-stage metadata.

**Fig. 4:**
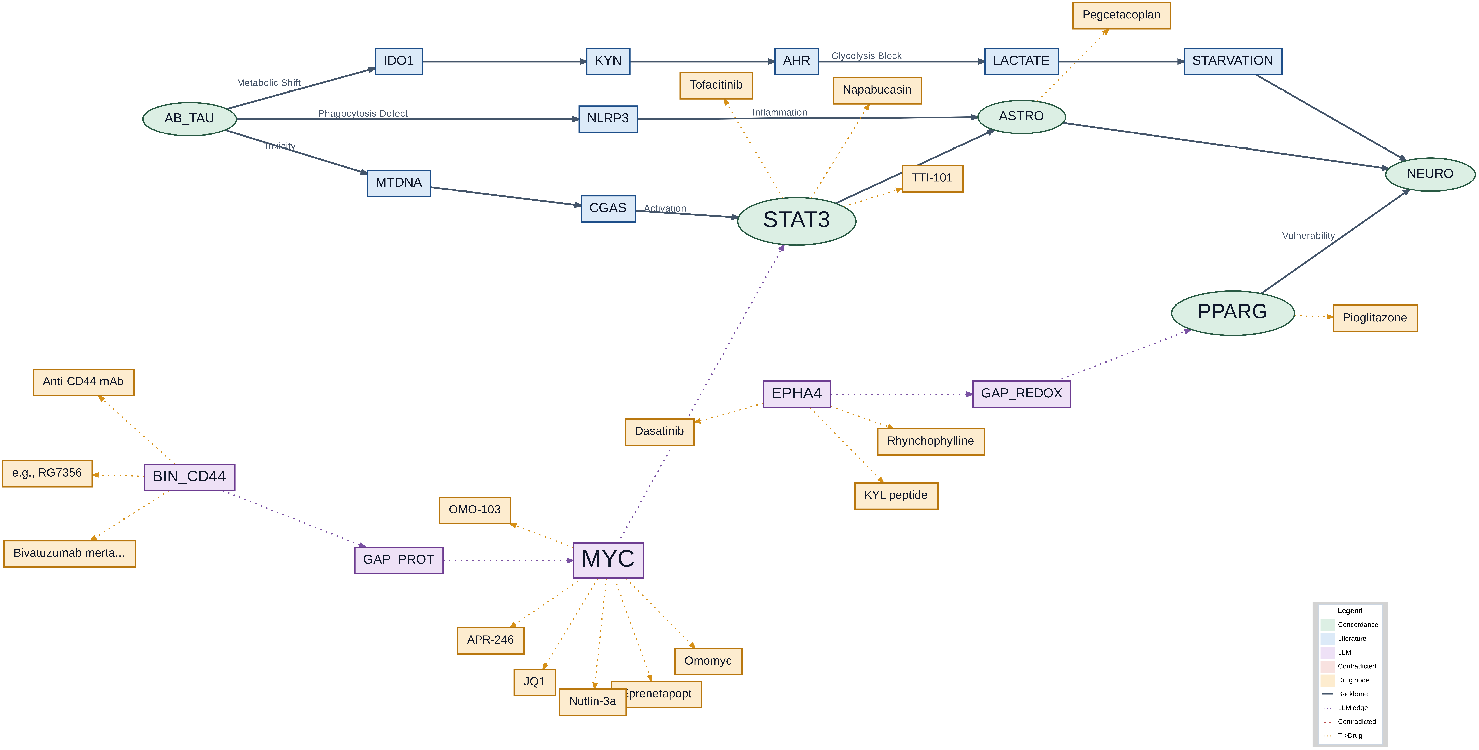
AD core drug-enabled concordance network (manual-review-seeded output). This output was generated from manually curated AD mechanistic exemplar literature. Target node size follows quantitative network support, and target-to-drug links preserve direction-aware literature evidence and development-stage metadata.

### 3.13 Cross-Disease Synthesis

Viewed jointly, PDAC and AD results delineate two operating regimes of the same workflow. In PDAC, the malignant-versus-reference contrast is sharper and leads to a larger integrated candidate universe (75 genes), whereas AD requires stronger compression to maintain interpretability (34 genes). Despite this difference in scale, quadrant structure remains comparably balanced in both diseases (PDAC: *Q*1*/Q*2*/Q*3 = 27*/*22*/*26; AD: 10*/*14*/*10), indicating that text priors, overlap-supported anchors, and network-emergent novelty all continue to contribute meaningful evidence rather than collapsing into one dominant source.

The downstream mechanism layer shows a similar pattern of shared structure with disease-specific intensity. PDAC generated 23 final strategies, and AD generated 14, both within a tractable combinational range. In both branches, provenance closure was preserved from candidate pool to strategy output, supporting the claim that hypothesis upgrading did not introduce untracked targets. The practical implication is that the framework can preserve mechanistic traceability even when disease-specific filtering pressure differs substantially.

Validation further clarifies where the two diseases diverge and where they converge. At target resolution, PDAC shows stronger directional separation, consistent with concentrated oncogenic dependence, while AD shows moderate but significant effects under the expanded CRISPRbrain setting, consistent with distributed neurodegenerative signal geometry. At strategy resolution, however, both branches exhibit strong non-random enrichment with identical global right-tail significance (*p* = 0.00019996). The corresponding global *z*-scores differ in magnitude (PDAC: 8.9883; AD: 6.4864), but both remain in a high-confidence range. Local significance rates are also comparable in practical terms (PDAC: 13/23; AD: 7/14), indicating that each branch yields an actionable strategy subset rather than isolated outliers. Taken together, these results support a cross-disease interpretation centered on robustness rather than direct score competition. PDAC represents a high-contrast setting in which effect concentration is stronger, while AD represents a heterogeneous setting in which stricter candidate control and context-matched validation are essential. The key finding is that the same provenance-aware retrieval-network-hypothesis-validation pipeline remains coherent, statistically productive, and interpretable across both conditions.

**Table 14:**
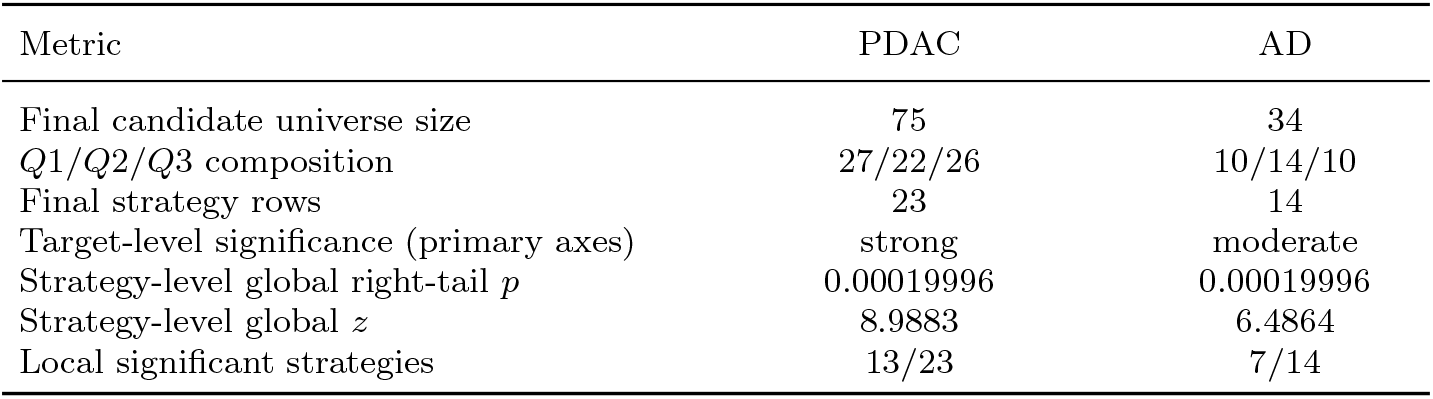
Cross-disease summary of key pipeline outputs.

## 4 Discussion

### 4.1 Retrospective Concordance: Validating Hypotheses Against Emergent Literature

Since Text-to-Target integrates retrieval priors and PathFinder-derived topology before generating any output, a meaningful test of the framework is not whether it recovers known targets but whether the *mechanism sequences* it produces remain coherent as experimental literature evolves. We therefore mapped generated hypotheses against recent independent benchmarks in PDAC and AD, examining both what the framework got right and where it fell short.

#### Predictive Robustness in PDAC

In PDAC, the framework recovered a high-value therapeutic axis that is concordant with recent resistance-focused studies. Specifically, it prioritized EGFR–STAT3 coupling, which matches subsequent reports supporting a KRAS–EGFR–STAT3 triple-axis strategy for suppressing refractory tumor growth and resistance [19, 25, 33]. This alignment indicates that the framework can identify clinically relevant control backbones before full experimental consolidation.

Concordance also appears at the mechanism level, not only at the target level. Generated sequences (for example, Upgraded-Hypo-15 and Table 3, Hypo-7) captured an EGFR-to-STAT3 relay and positioned STAT3 as a hub linked to mesenchymal plasticity and EMT-associated states through factors such as VIM and CTCF [2, 27]. These links are consistent with desmoplastic and invasive phenotypes in resistant PDAC, supporting the biological coherence of the inferred signaling backbone.

At the same time, retrospective comparison identifies an important resolution gap. Our earlier hypotheses emphasized the canonical IL6-JAK-STAT3 bridge, which is common in steady-state priors, whereas newer perturbation-focused studies in KRAS/EGFR-inhibited settings indicate an alternative SRC-family route, including FYN-mediated Tyr705 activation of STAT3 [19, 33, 38]. This mismatch illustrates “canonical inertia”: LLM priors can preserve broadly valid motifs while missing context-specific rewiring under treatment pressure.

The same retrospective lens becomes more informative when projected onto a drug-enabled layer. Instead of asking only whether a target reappears in later literature, we can evaluate whether target-linked compounds carry convergent, mixed, or contradictory support in the same mechanistic context. This extension improves translational interpretability: backbone-consistent routes with support-dominant drug links become higher-priority intervention candidates, whereas contradiction-heavy links are explicitly flagged for perturbation-specific re-evaluation before strategy escalation.

#### Topological Robustness in AD

In AD, retrospective mapping shows strong macro-level concordance between inferred topology and later mechanistic evidence. The workflow prioritized *STAT3* as a central transcriptional regulator linked to reactive astrogliosis and *C3* induction [23, 39, 40], consistent with studies describing IL-6/STAT3 as an amplifier of neuroinflammatory feed-forward loops [23, 24, 41]. It also highlighted a *PPARG*-centered metabolic-inflammatory coupling, concordant with reports connecting energy-state reprogramming to both astroglial toxicity and tau-related pathology [39, 42, 43].

However, as in PDAC, concordance at the backbone level does not guarantee molecular-level completeness. The model identified “Proteostasis” and “Redox” junctures as critical gaps but represented them as abstract placeholders. Recent studies provide finer-grained fillers, including mtDNA-associated *cGAS-STING*/*NLRP3* signaling for proteostatic stress and an *IDO1-KYN-AhR* axis for metabolic immune hijacking [23, 41, 44]. These comparisons suggest that Text-to-Target is effective for topology-aware triage, while perturbation-resolved updating remains necessary for high-resolution mechanism specification.

### 4.2 Prospective Design: A Continuous Literature-Mechanism Concordance Framework

The retrospective analysis identifies a consistent failure mode across diseases: stable backbone recovery with incomplete perturbation-specific detail. To address this gap, we propose a continuous Literature-Mechanism Concordance layer that replaces onetime retrospective checking with ongoing, perturbation-aware updating [45].

The key design choice is to use ordered mechanism edges, rather than flat target sets, as the operational unit. Each hypothesis is serialized into directed relations (for example, *A* → *B, B* → *C*) with explicit context tags (disease, cell type, perturbation mode, and directionality). A retrieval-augmented evidence pipeline continuously collects claim-level support for each edge [46, 47], and stores each evidence object in a normalized schema that records support polarity (support/contradict/mixed), experimental modality (in vitro, in vivo, cohort, perturbation screen), and temporal metadata.

Edge-level evidence is then aggregated into sequence-level concordance profiles using interpretable metrics such as support coverage, contradiction burden, and recency-weighted support strength. This scoring separates broadly supported sequences from conflict-prone or weakly evidenced alternatives, enabling explicit triage instead of binary keep/drop decisions. Mechanism sequences are also versioned; when new studies change edge support, updated releases include transparent change logs and citation deltas.

Under this design, hypothesis validation becomes a living process rather than a static endpoint. For fast-moving domains such as neuroinflammation and combination oncology, continuous 7-day/24-hour operation is not an engineering preference but a methodological requirement for maintaining translational relevance and mechanistic precision over time.

In this prospective setting, edge-level concordance can be propagated directly into target-to-drug evidence maintenance. When an updated study strengthens, weakens, or reverses support for a mechanism edge, linked therapeutic annotations can be refreshed in the same version cycle by updating evidence direction and citation fields on corresponding target-to-drug edges. This creates a unified update surface in which mechanistic and therapeutic revisions are synchronized rather than managed as separate post hoc layers.

Our current PDAC/AD comparison offers an initial design signal for this prospective loop: manual-review-seeded core outputs provide a high-precision bootstrap, while automated retrieval expansion increases coverage and route diversity when normalized into the same node-evidence-edge-node schema. This staged pattern directly motivates an agentic operating model in which human-curated seeds initialize the system and autonomous retrieval-refresh agents maintain breadth over time.

### 4.3 Agentic Outlook

A practical implementation is an always-on 7-day/24-hour agentic loop with explicit role separation across evidence acquisition, interpretation, and release control. In this architecture, literature watchers monitor new outputs; retrieval and extraction services convert documents into structured edge evidence; adjudication services update support status; hypothesis revision services regenerate mechanism sequences when contradictions accumulate; validation orchestration services trigger scheduled re-analysis; and release governance services publish only versioned snapshots that pass predefined quality gates. OpenClaw-style computer-use/tool-use modules can be added at the acquisition boundary to automate source monitoring, parsing, and citation normalization.

To reduce drift, this loop should run at multiple cadences. Fast cycles (6–24 hours) update evidence ledgers and contradiction flags, medium cycles (weekly) revise or prune hypothesis branches, and slow cycles (monthly) freeze stable releases for benchmarking. Progress should be tracked longitudinally with concordance gain per cycle, hypothesis stability under refresh, novelty-to-support conversion, and incremental changes in downstream validation statistics.

This outlook aligns with the broader shift from one-shot prompting to tool-grounded iterative scientific agents [48–50]. In our setting, continuous operation is valuable only insofar as it improves mechanism fidelity and translational utility as the evidence landscape changes.

### 4.4 Limitations and Future Work

The current framework has clear scope limitations. Transcriptomic signals support association-driven prioritization but do not establish causal sufficiency; post-transcriptional regulation, protein activity states, and spatial microenvironment effects remain incompletely modeled. In addition, AD validation depends on registry composition and assay heterogeneity, whereas PDAC validation may underrepresent microenvironment-mediated vulnerabilities when epithelial contrasts dominate.

Continuous 7-day/24-hour operation also creates new methodological risks, including retrieval-noise accumulation, citation leakage, policy drift across model updates, and self-reinforcing loops around highly cited but context-mismatched mechanisms. For this reason, governance is not optional infrastructure: it is part of the method definition.

The drug-enabled layer has additional practical boundaries in its current form. Node-size rendering is currently based on PathFinder importance weights, which are learned under DEG-informed constraints and therefore remain quantitatively linked to transcriptomic differential signals, but are not a direct visualization of a single gene-level statistic such as − log_10_(*p*). Interactive displays also expose citation granularity more strongly at edge level than at node summary level. Although full evidence objects are retained in JSON, manuscript-facing evidence digest tables still require explicit export. These limitations are tractable but should be addressed to strengthen reviewer-facing translational reporting.

Future work will focus on four directions: perturbation-coupled prospective feedback, spatial/proteomic augmentation, refreshable edge-level literature concordance, and governance with deterministic checkpoints and release policies. These upgrades would move Text-to-Target from high-quality hypothesis triage toward continuously improving, mechanism-auditable therapeutic discovery.

## Acknowledgment

Funding: This research was partially supported by NIA 4R33AG078799-02, NLM 1R01LM013902-01A1, NIA R56AG065352, NIA 1R21AG078799-01A1.

## 5 Supplementary Tables

**Table S1:**
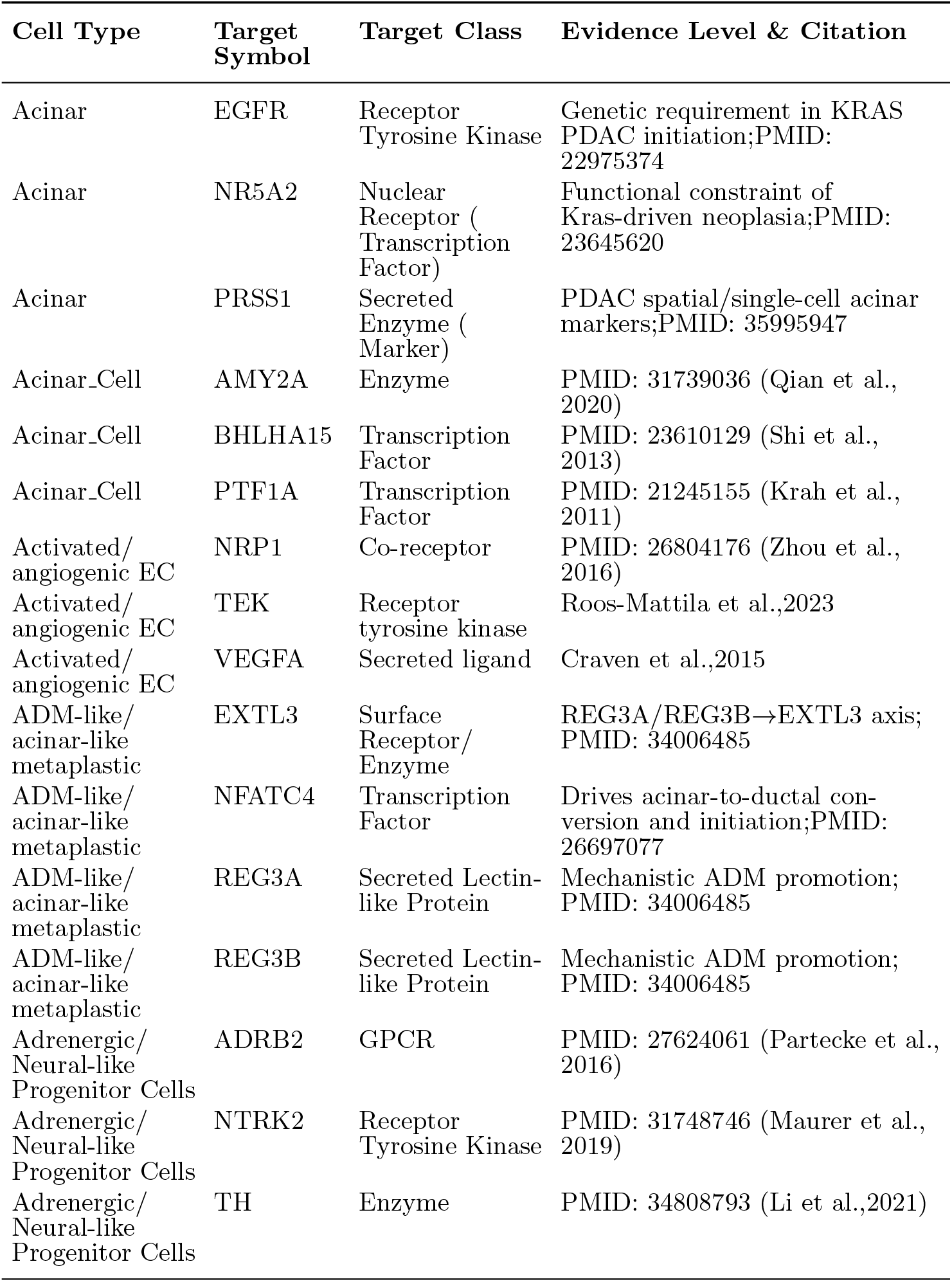
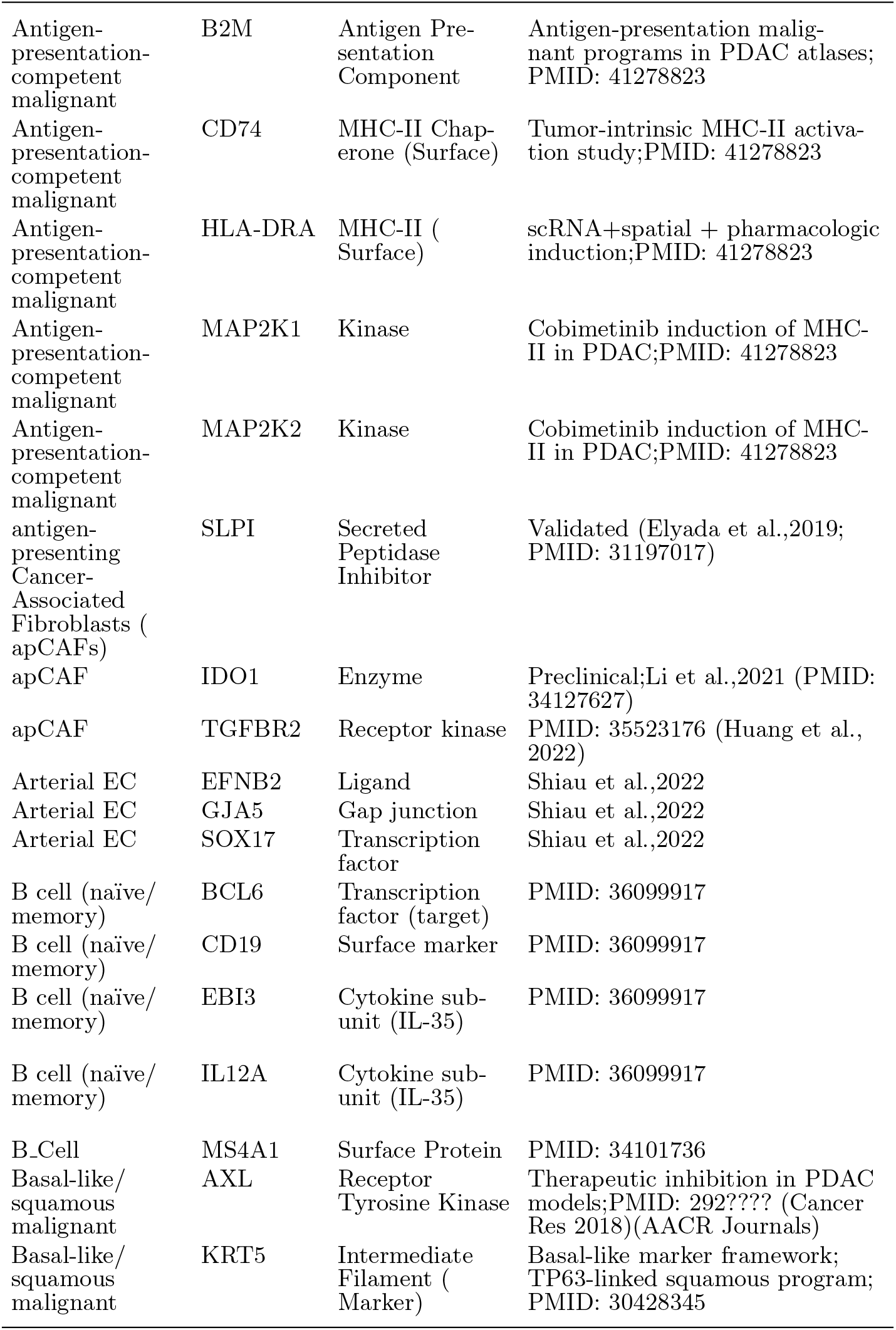
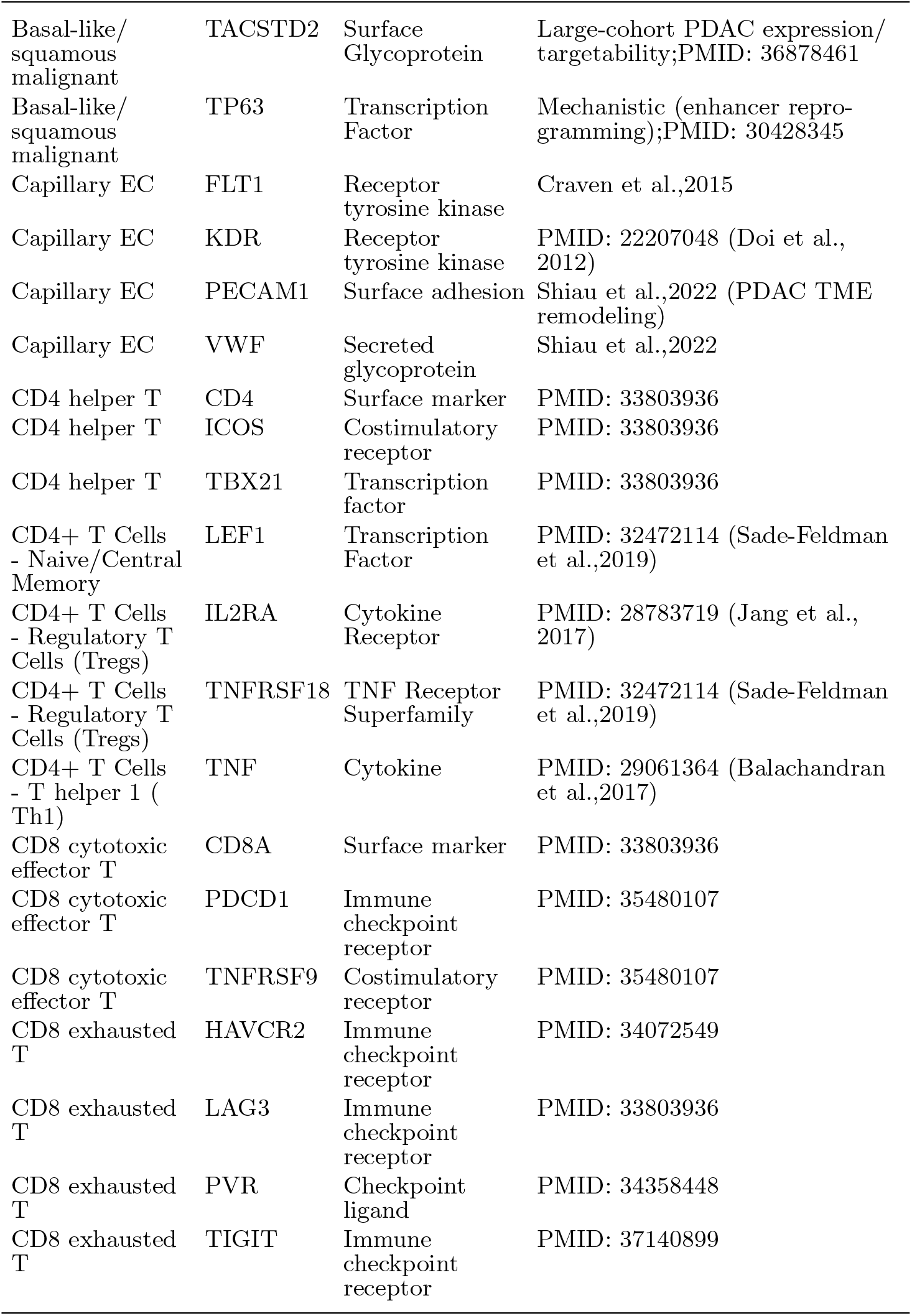
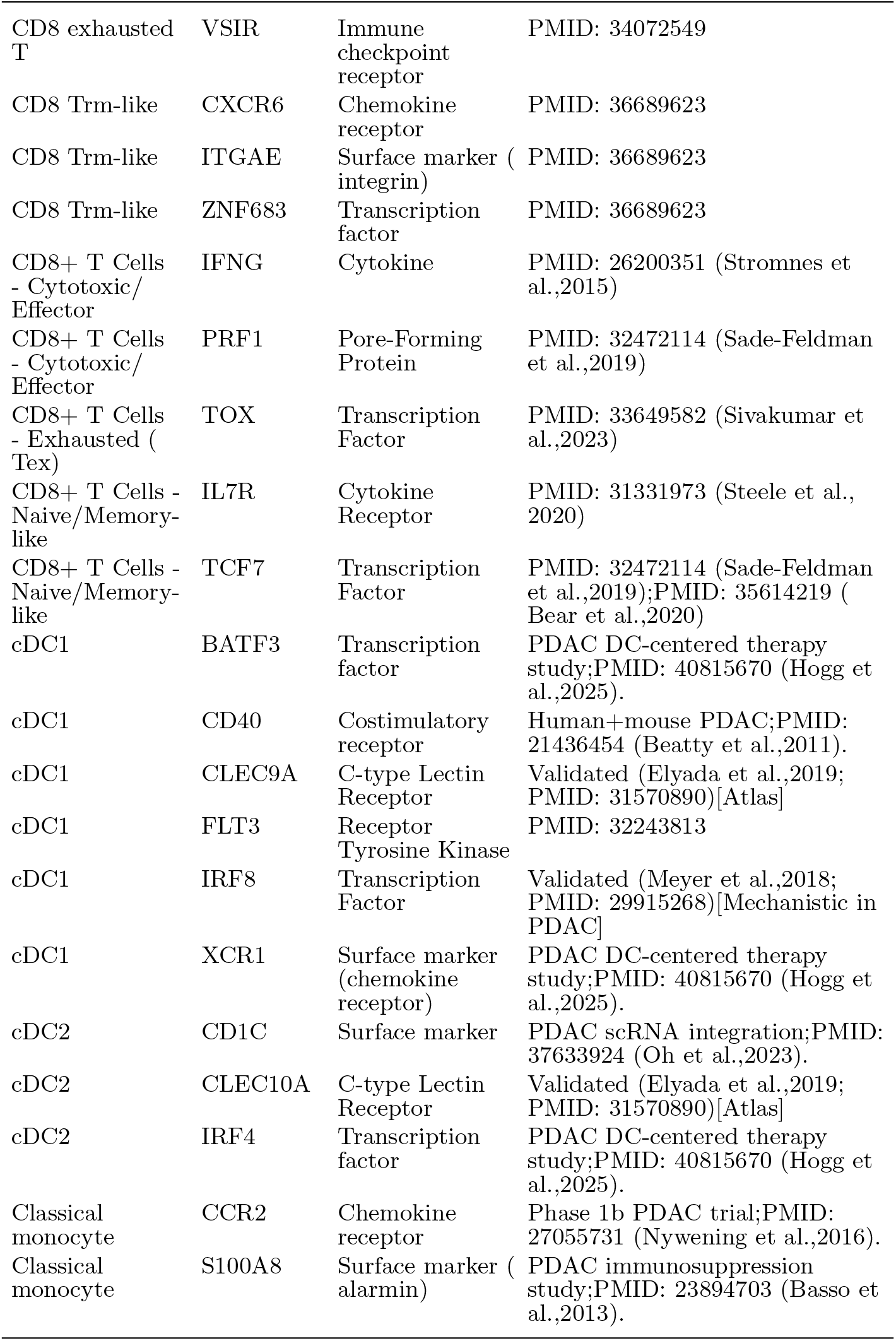
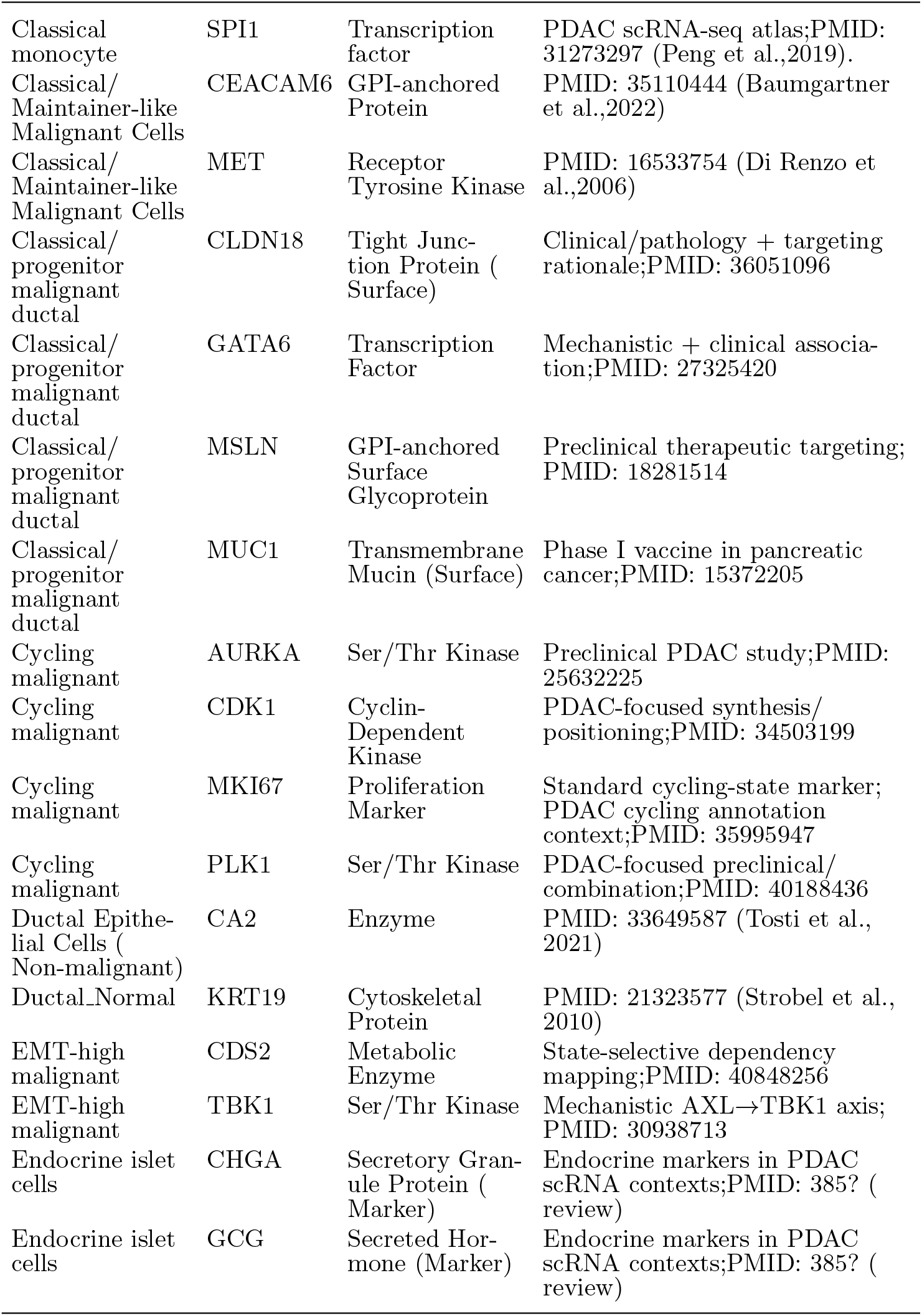
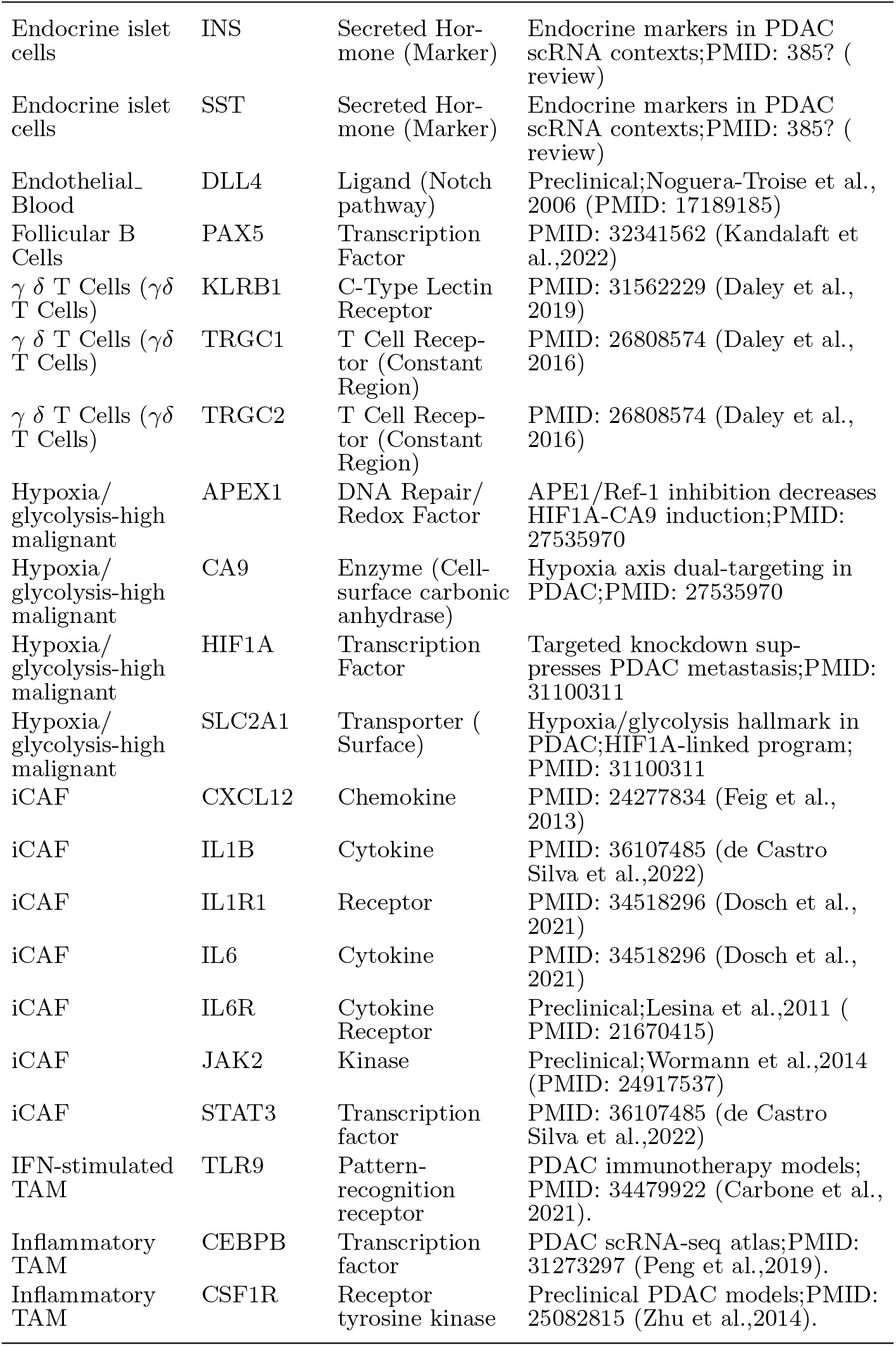
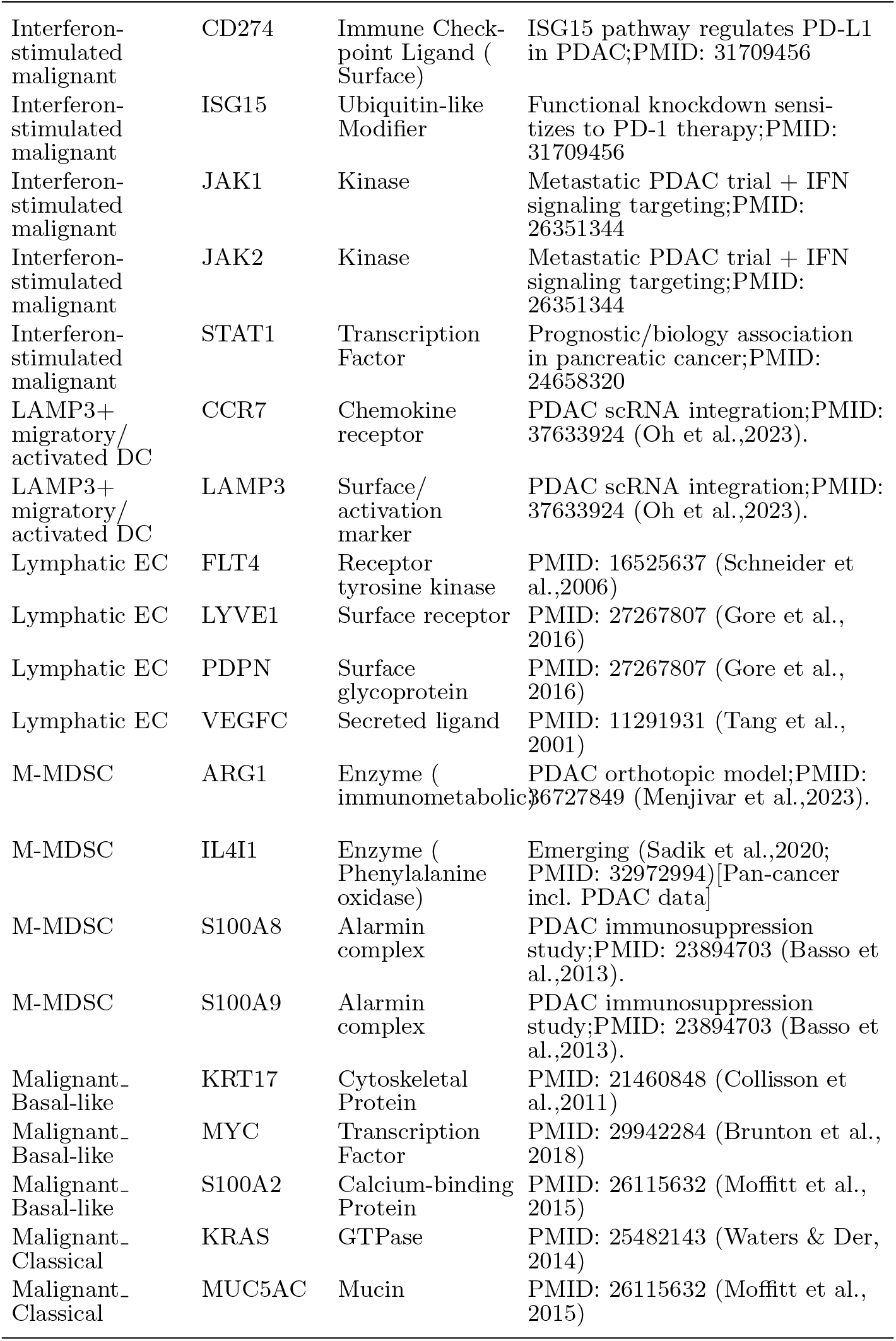
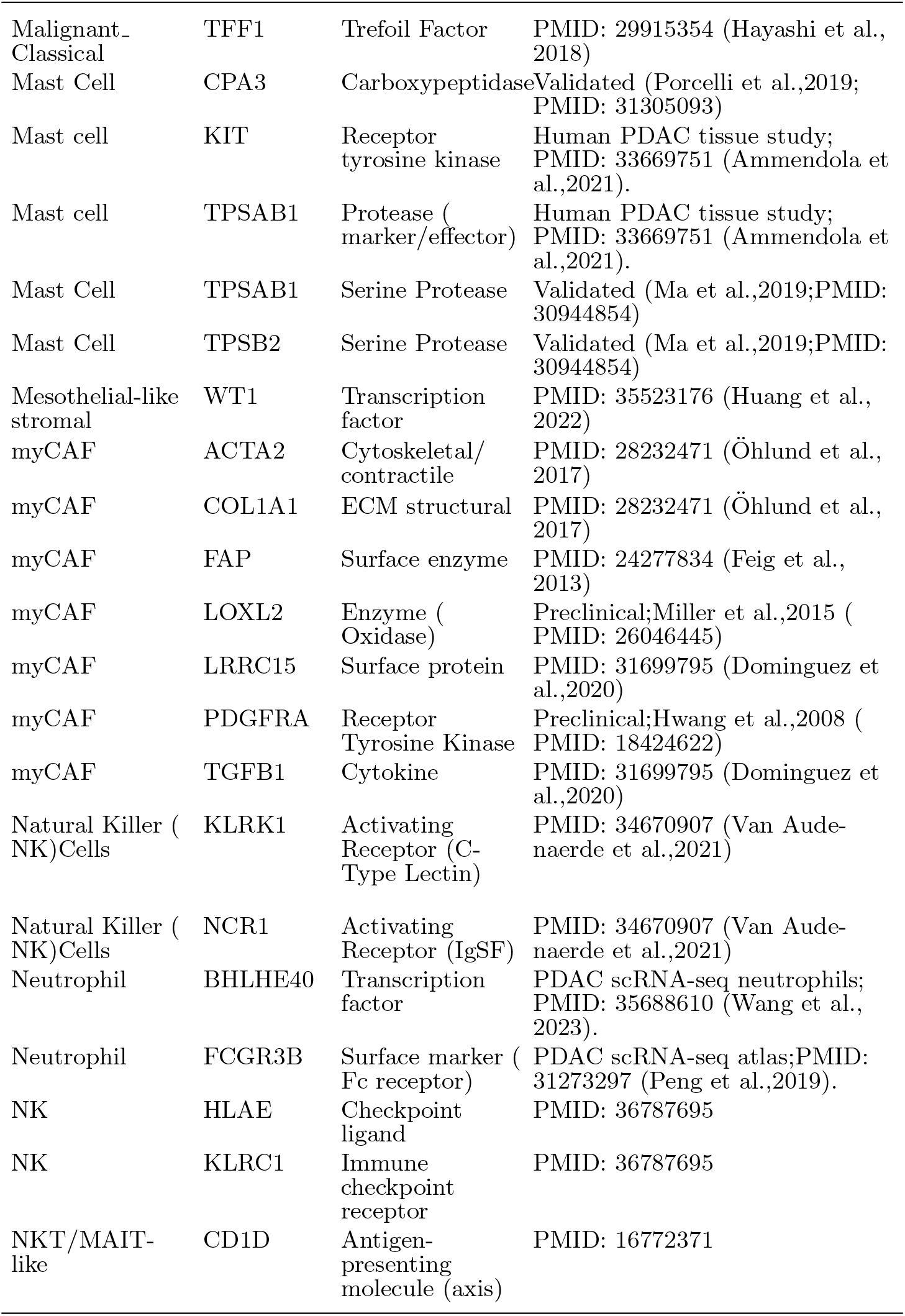
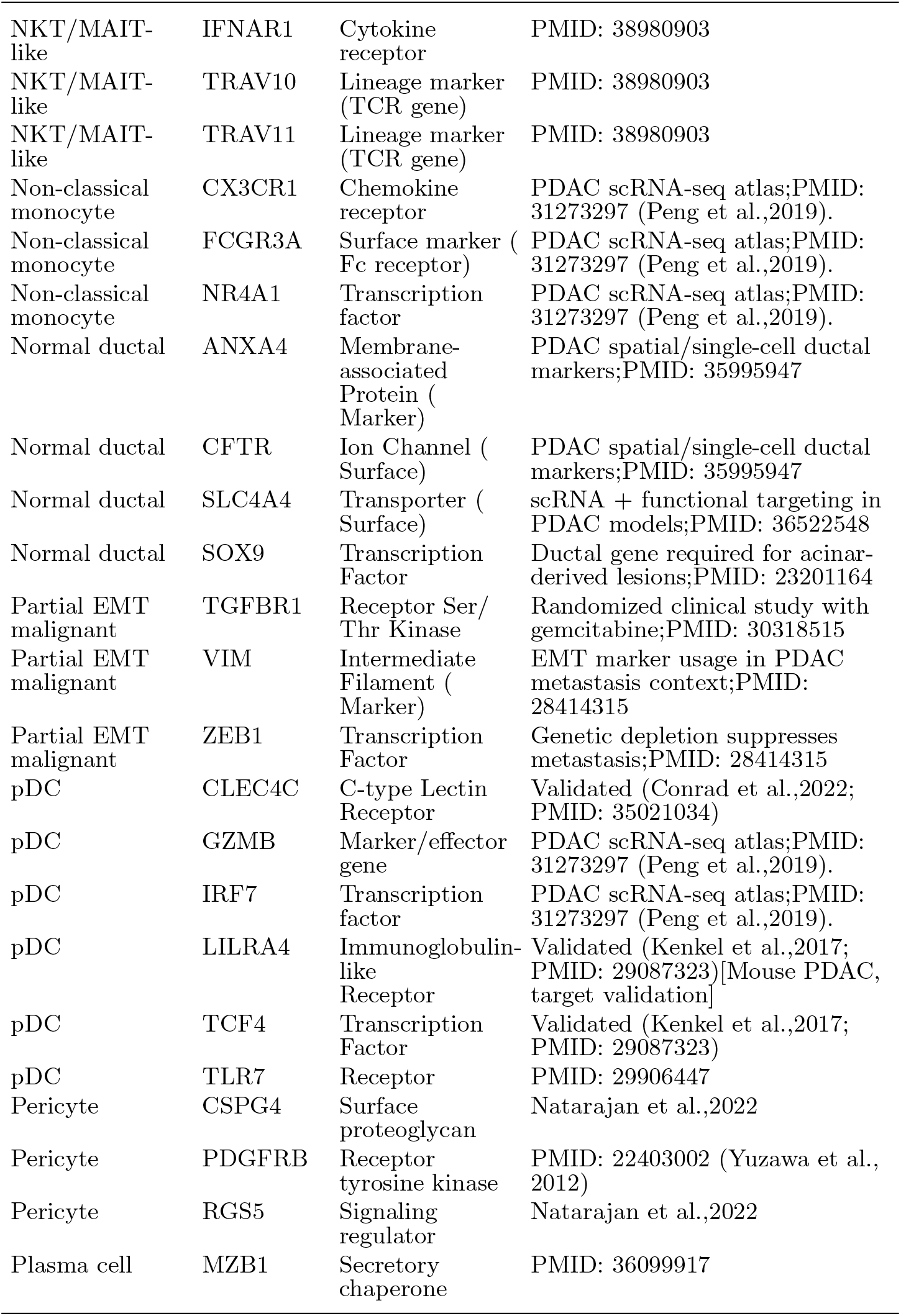
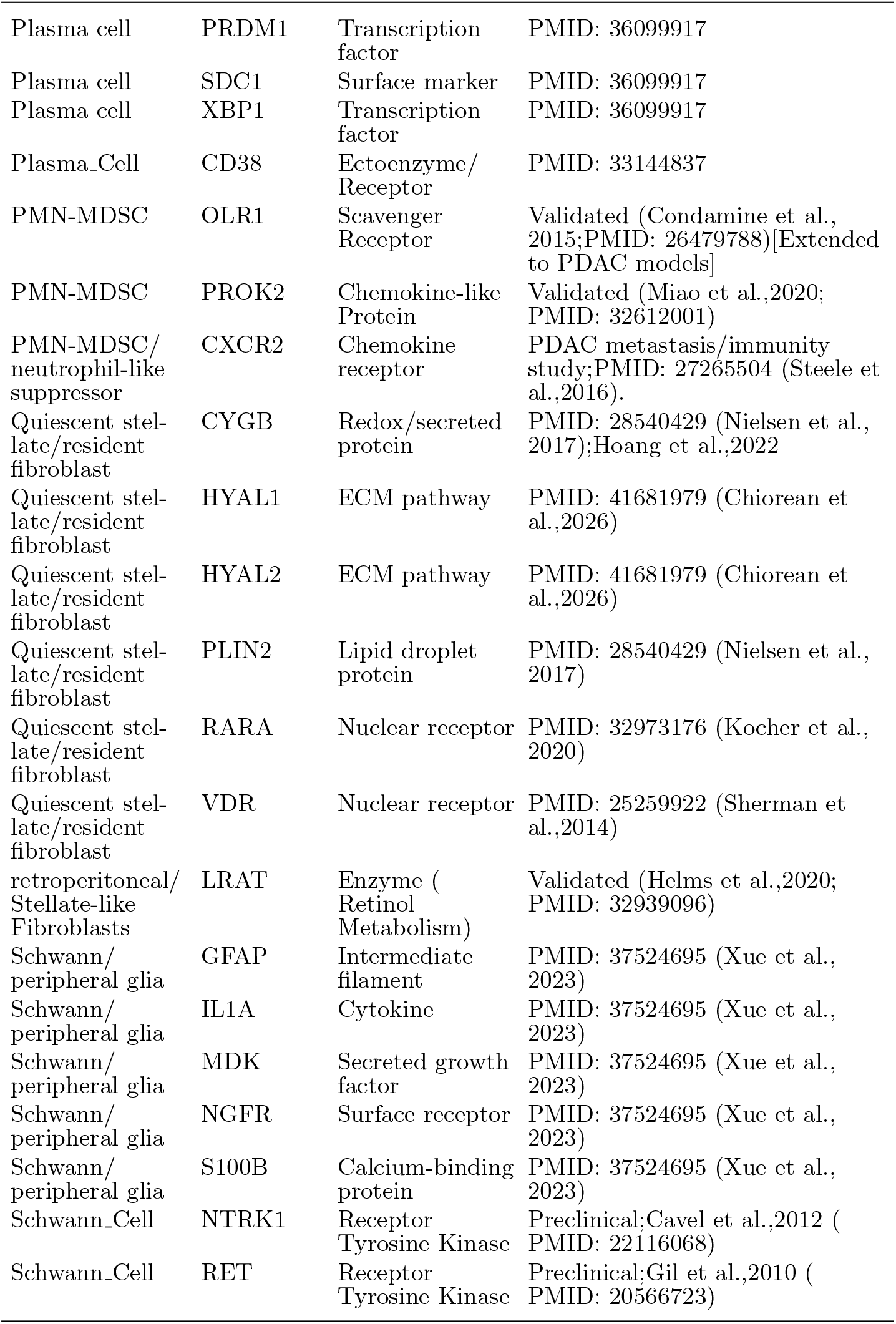
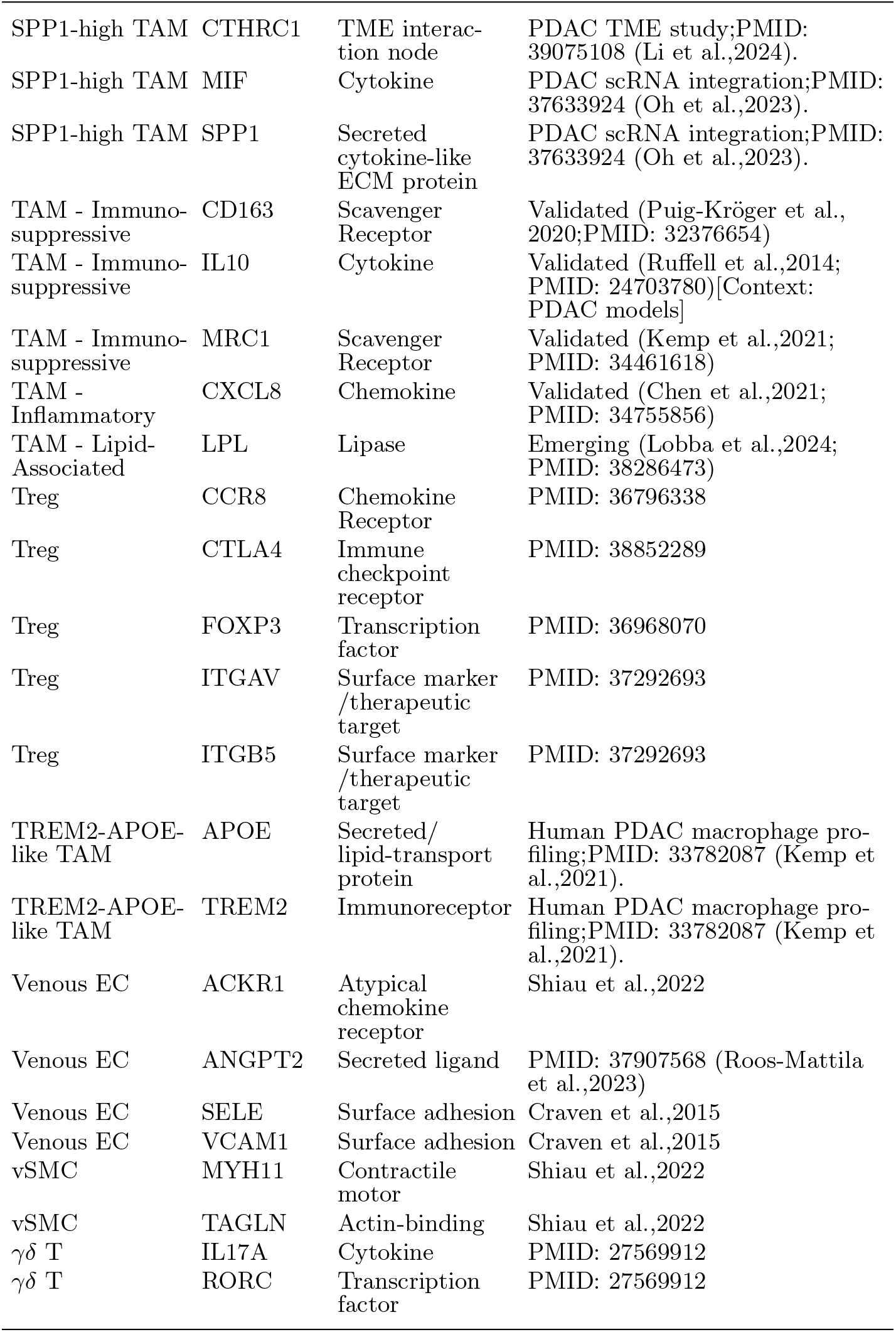
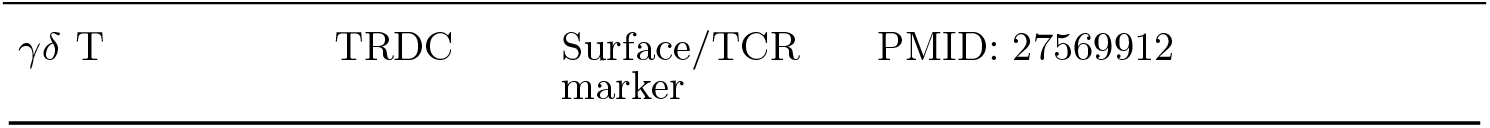
PDAC cell-specific targets.

**Table S2:**
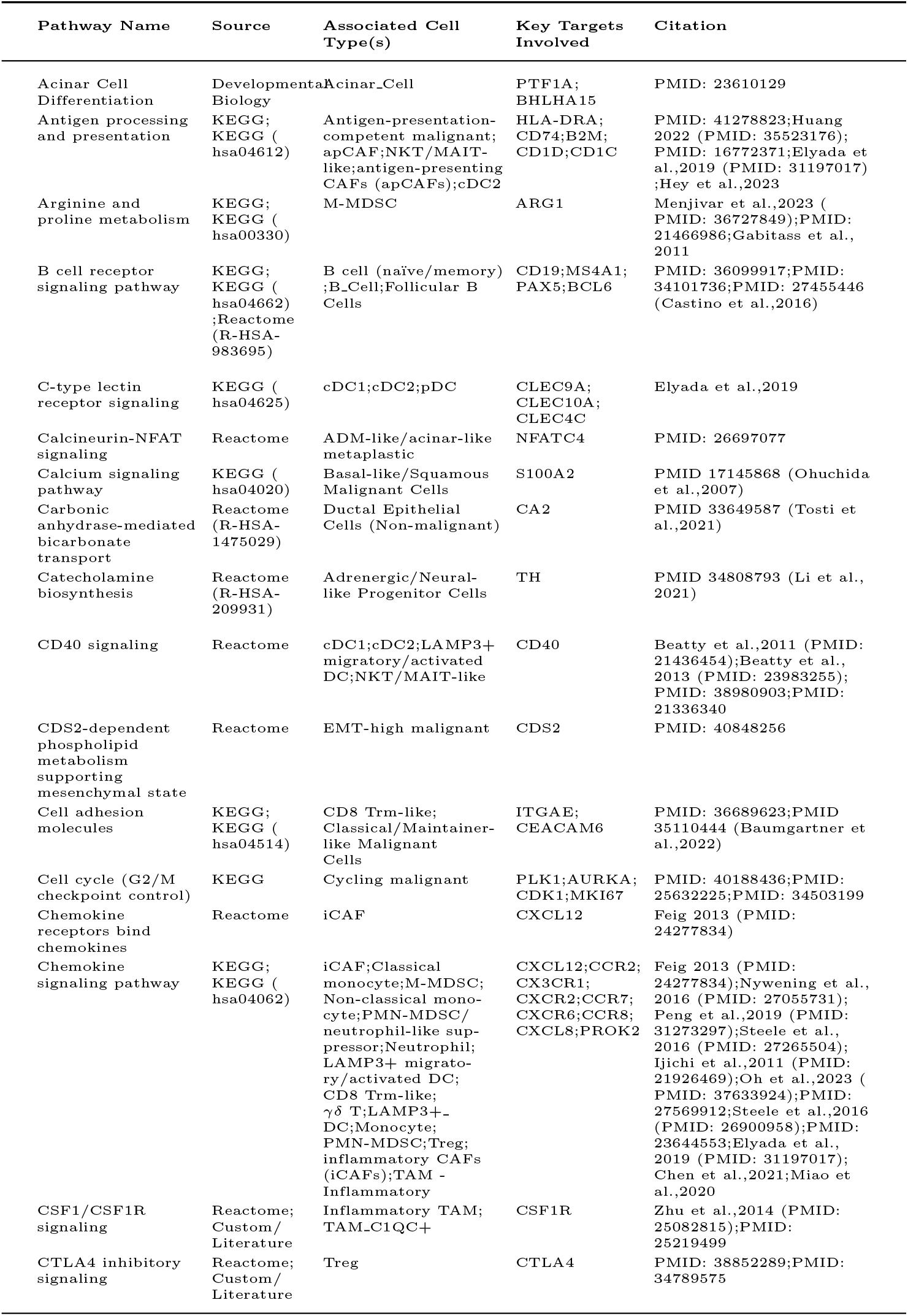
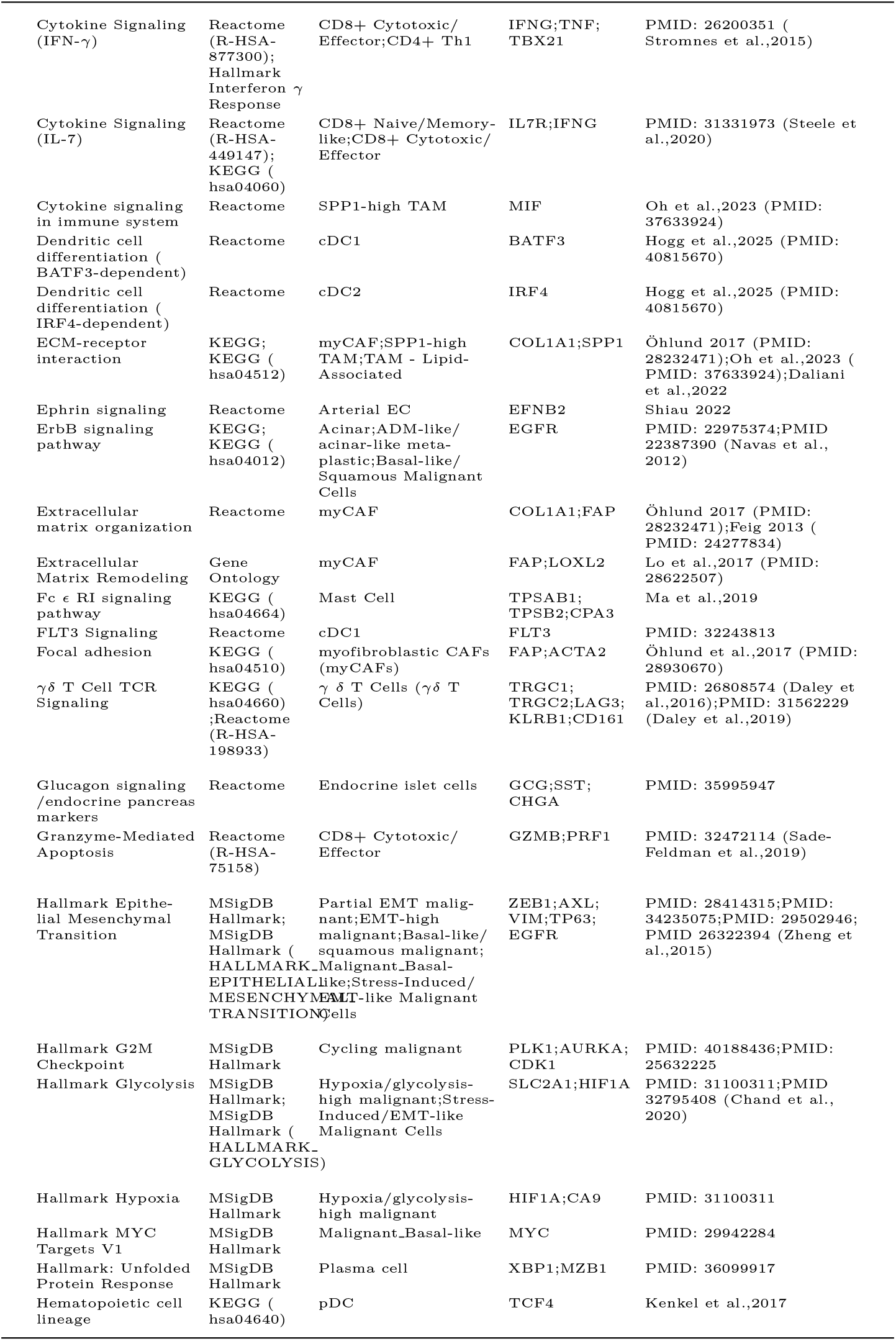
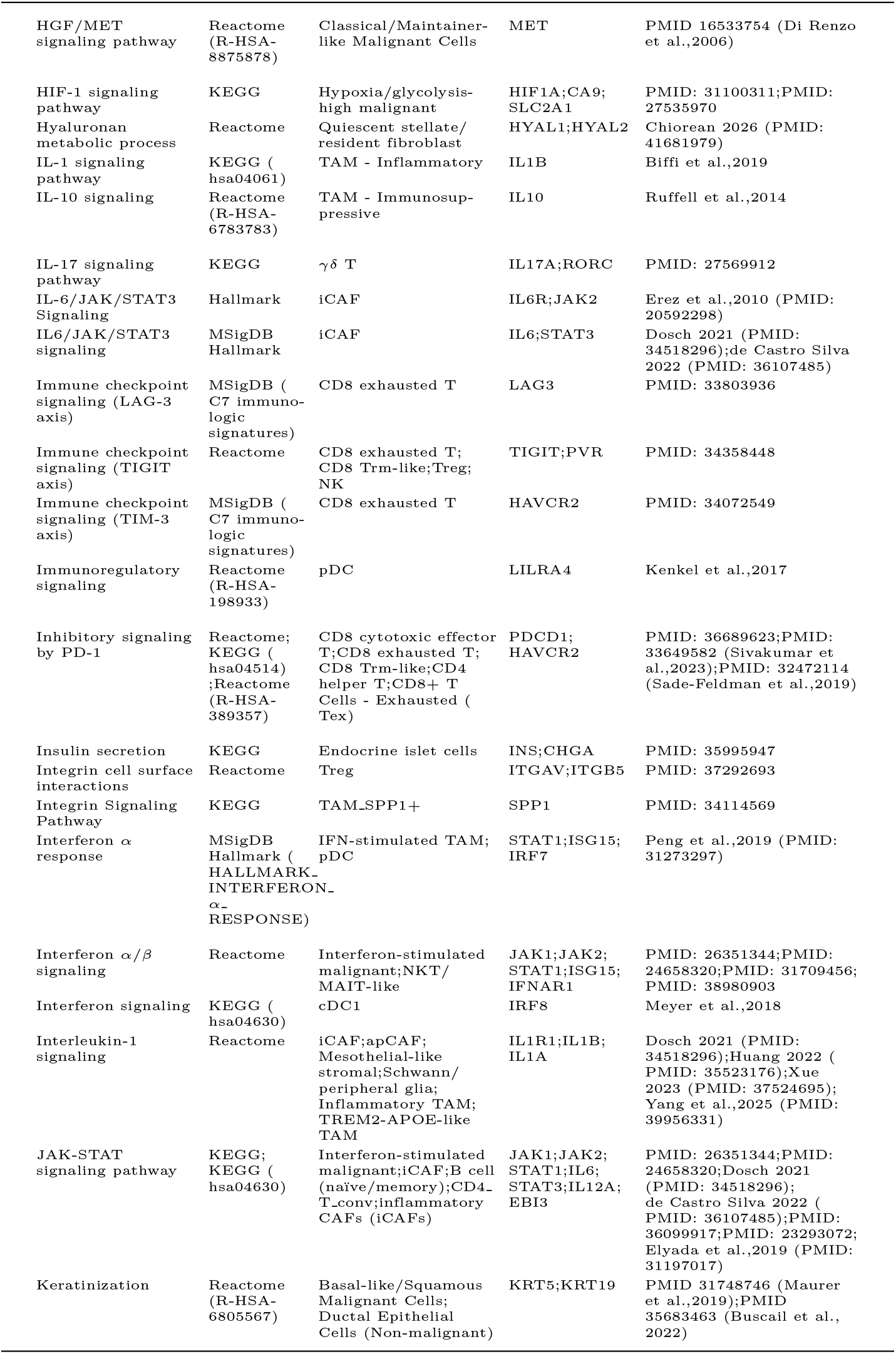
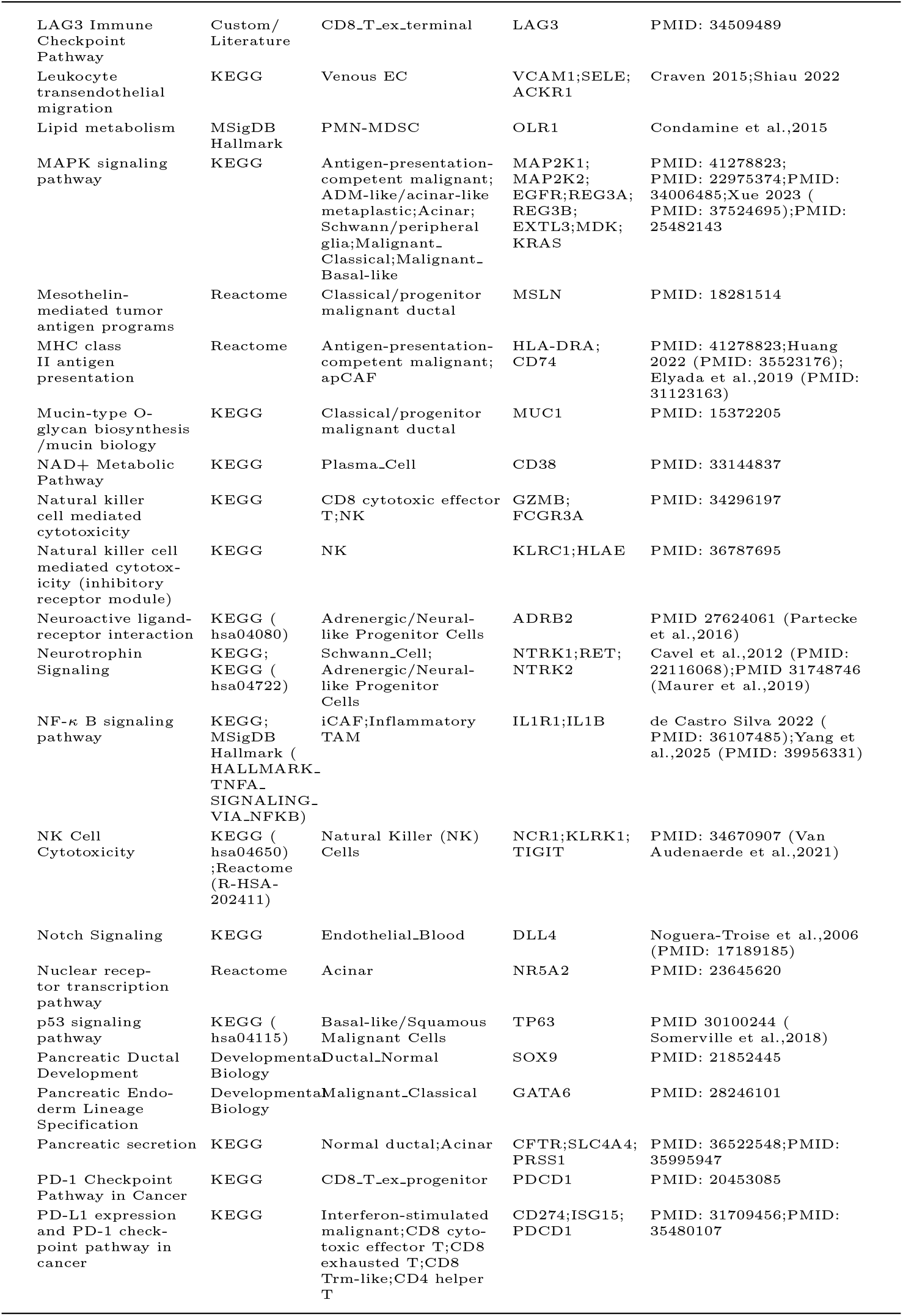
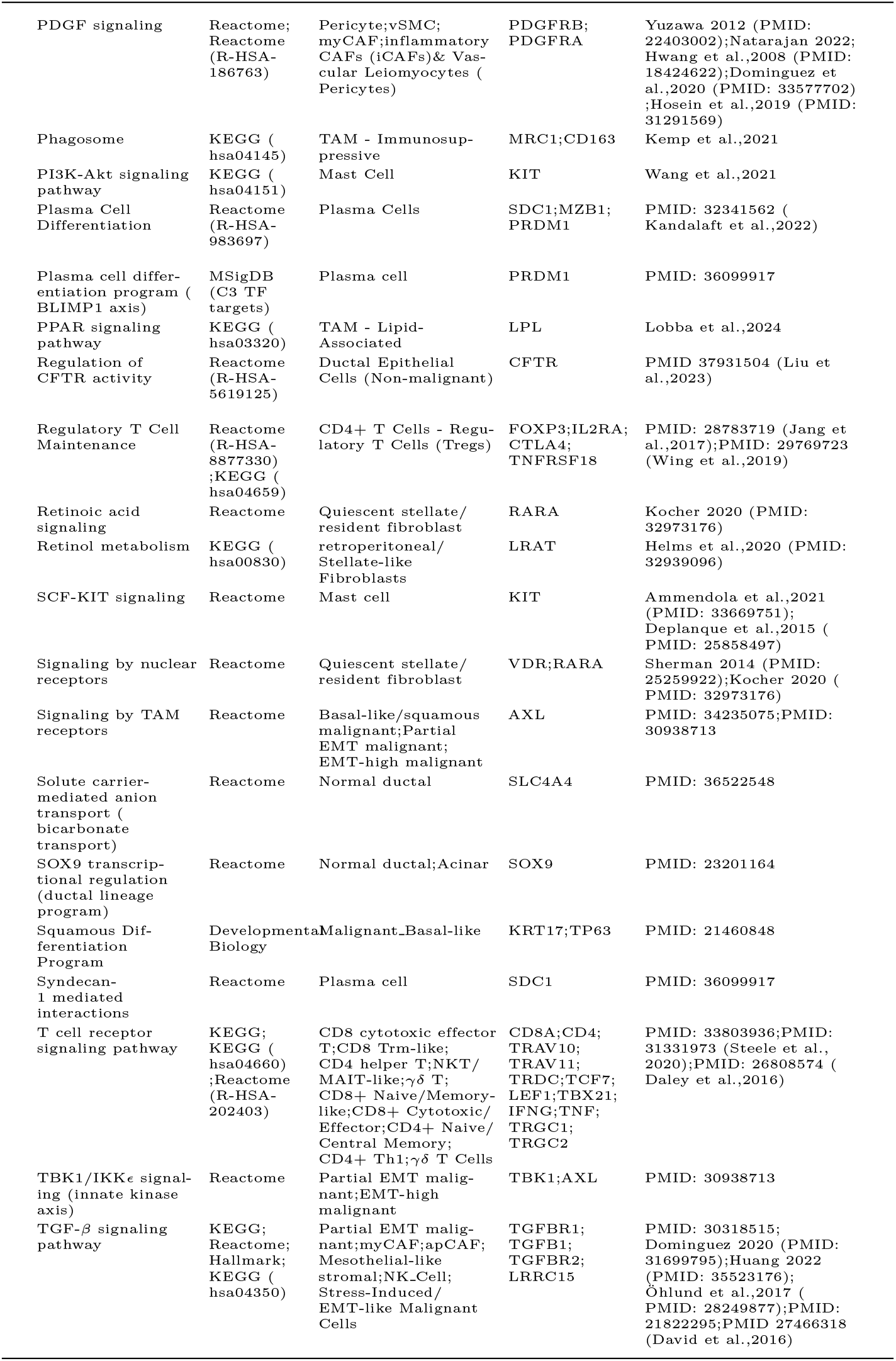
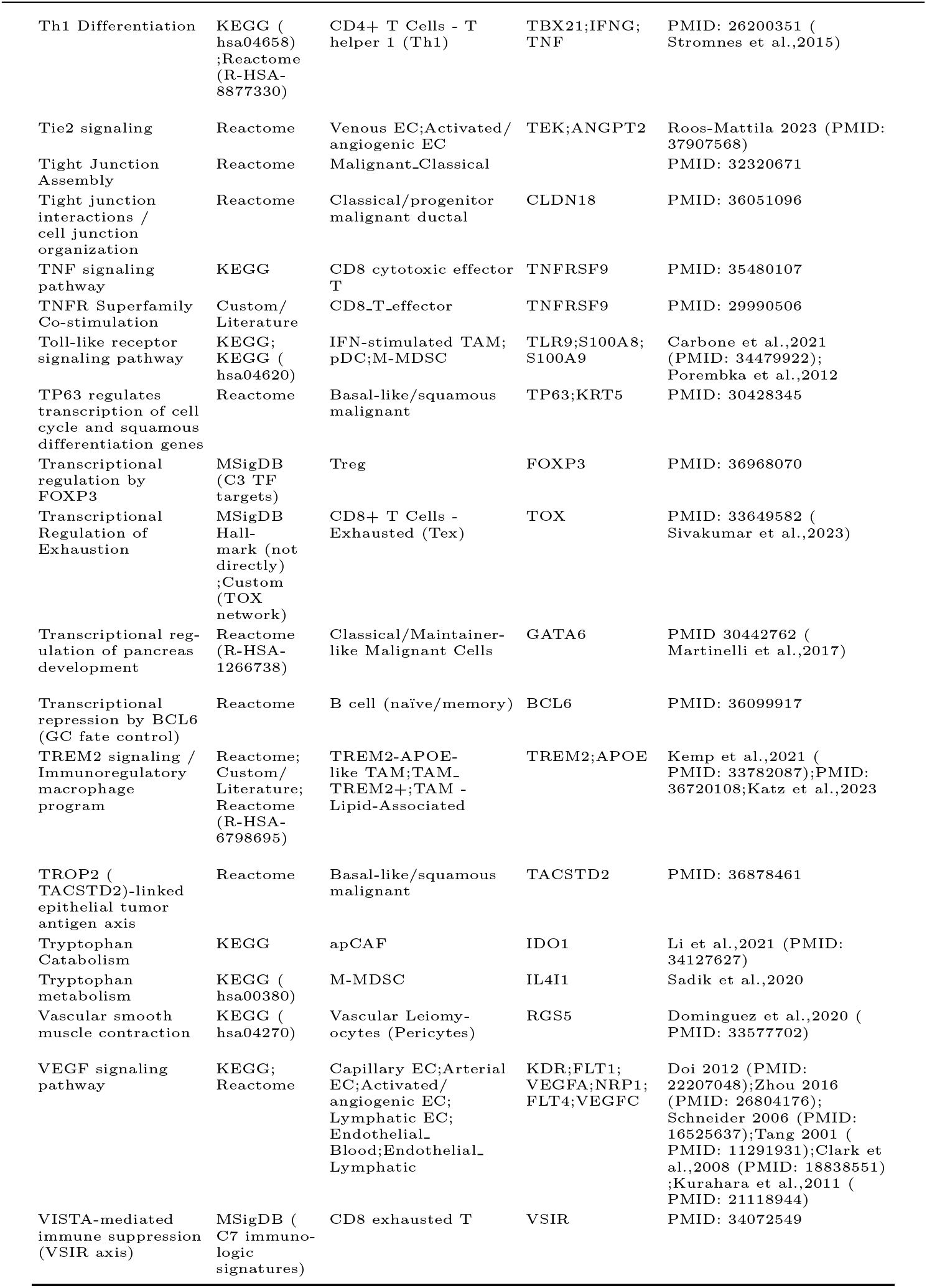
PDAC cell-specific pathways.

**Table S3:**
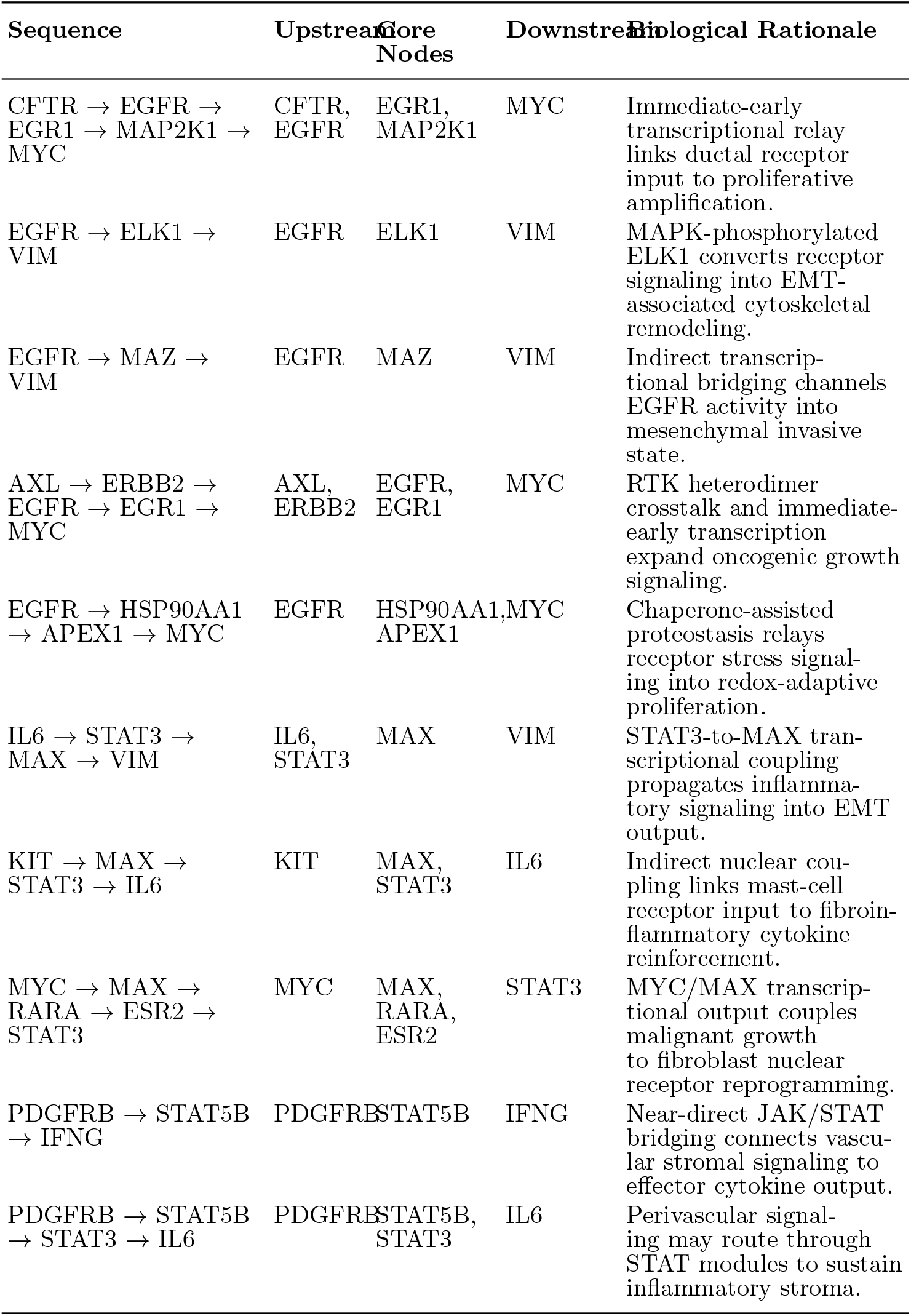
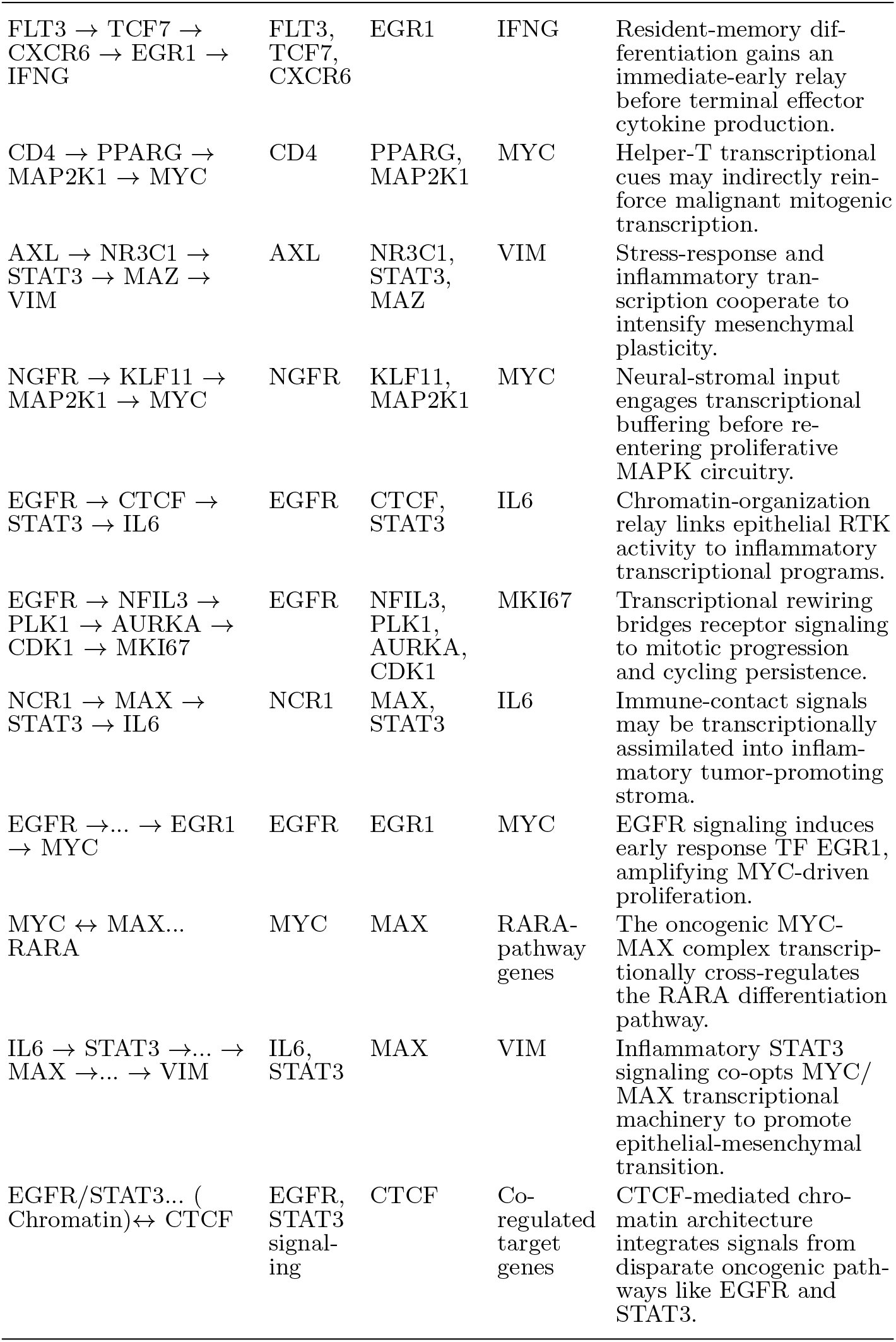
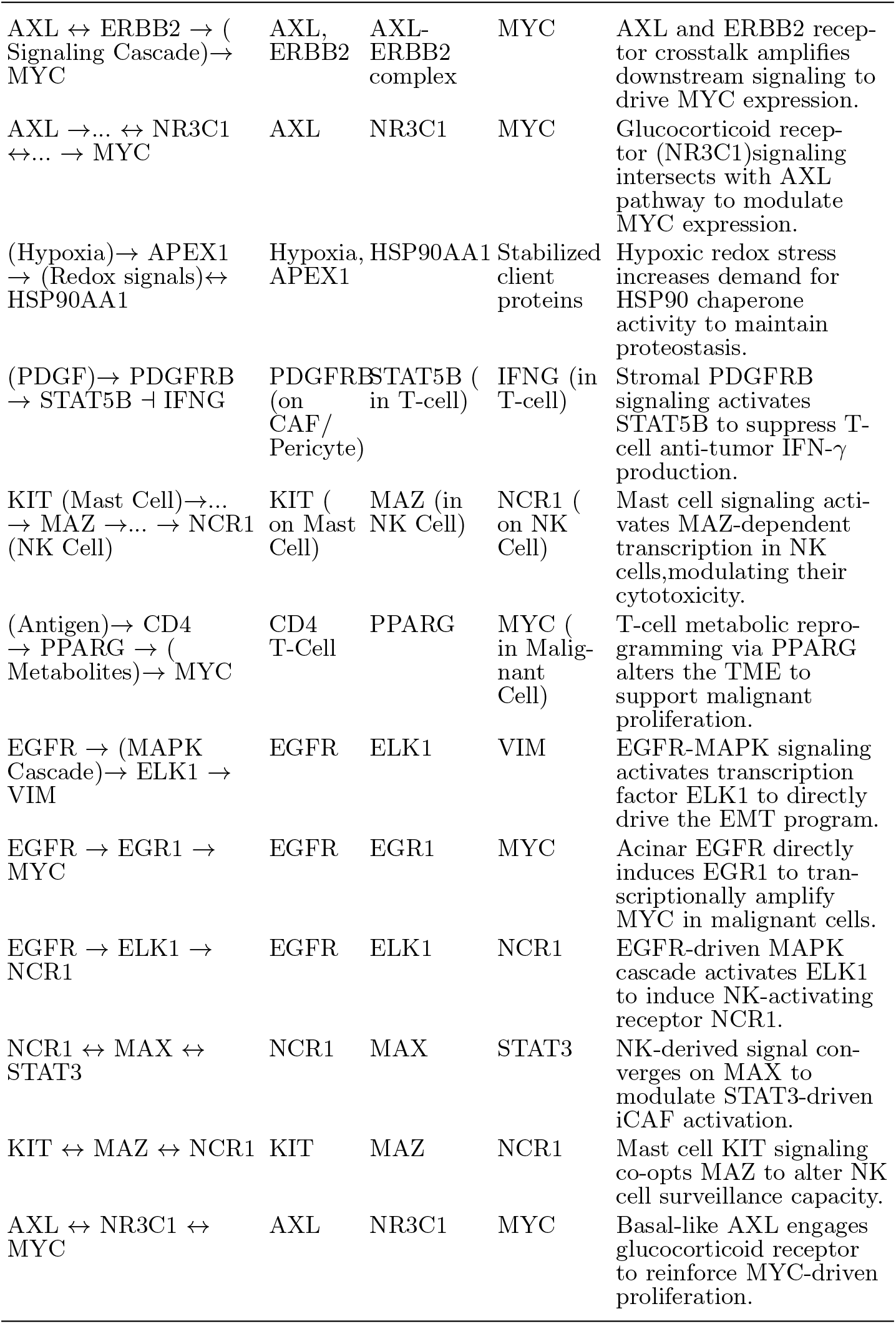
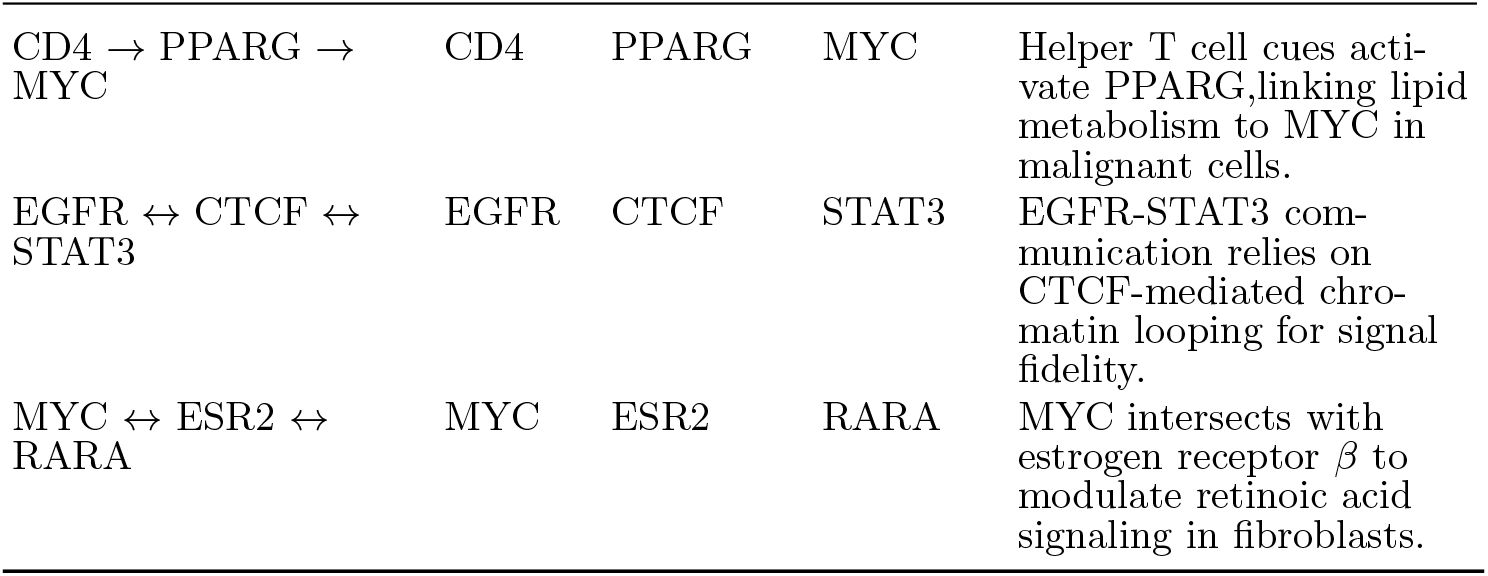
PDAC LLM-driven mechanism hypotheses.

**Table S4:**
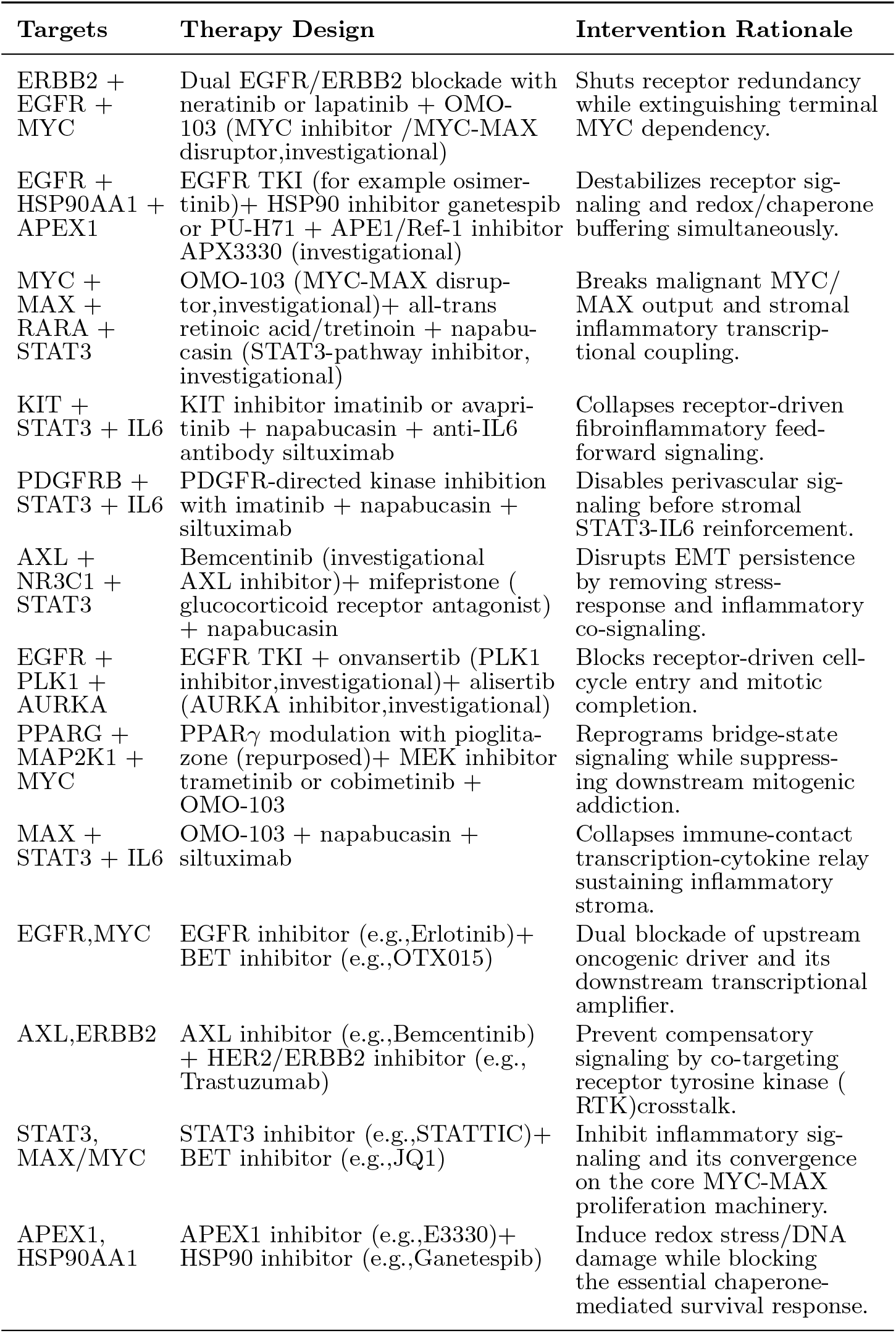
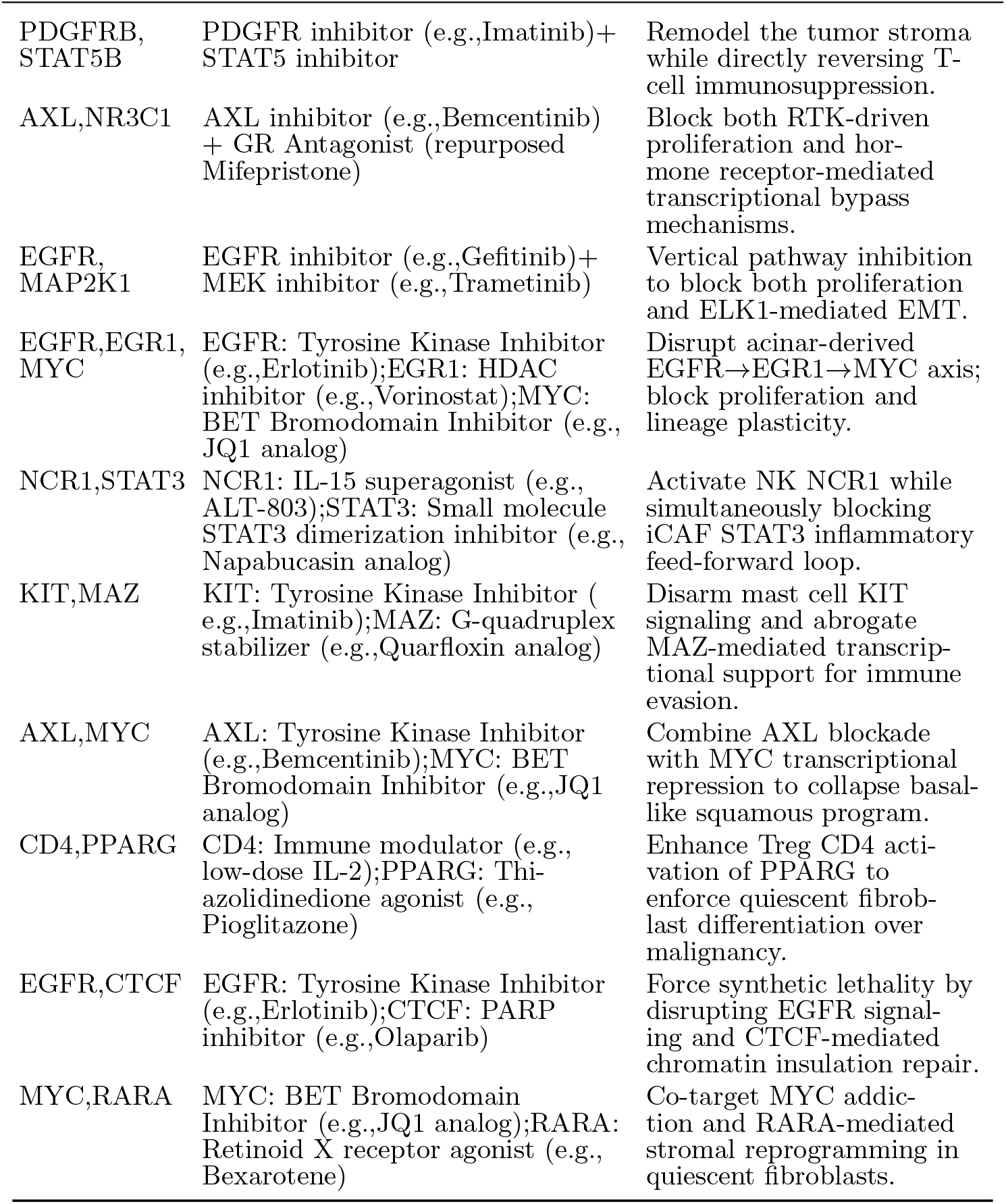
PDAC upgraded therapy combinations.

**Table S5:**
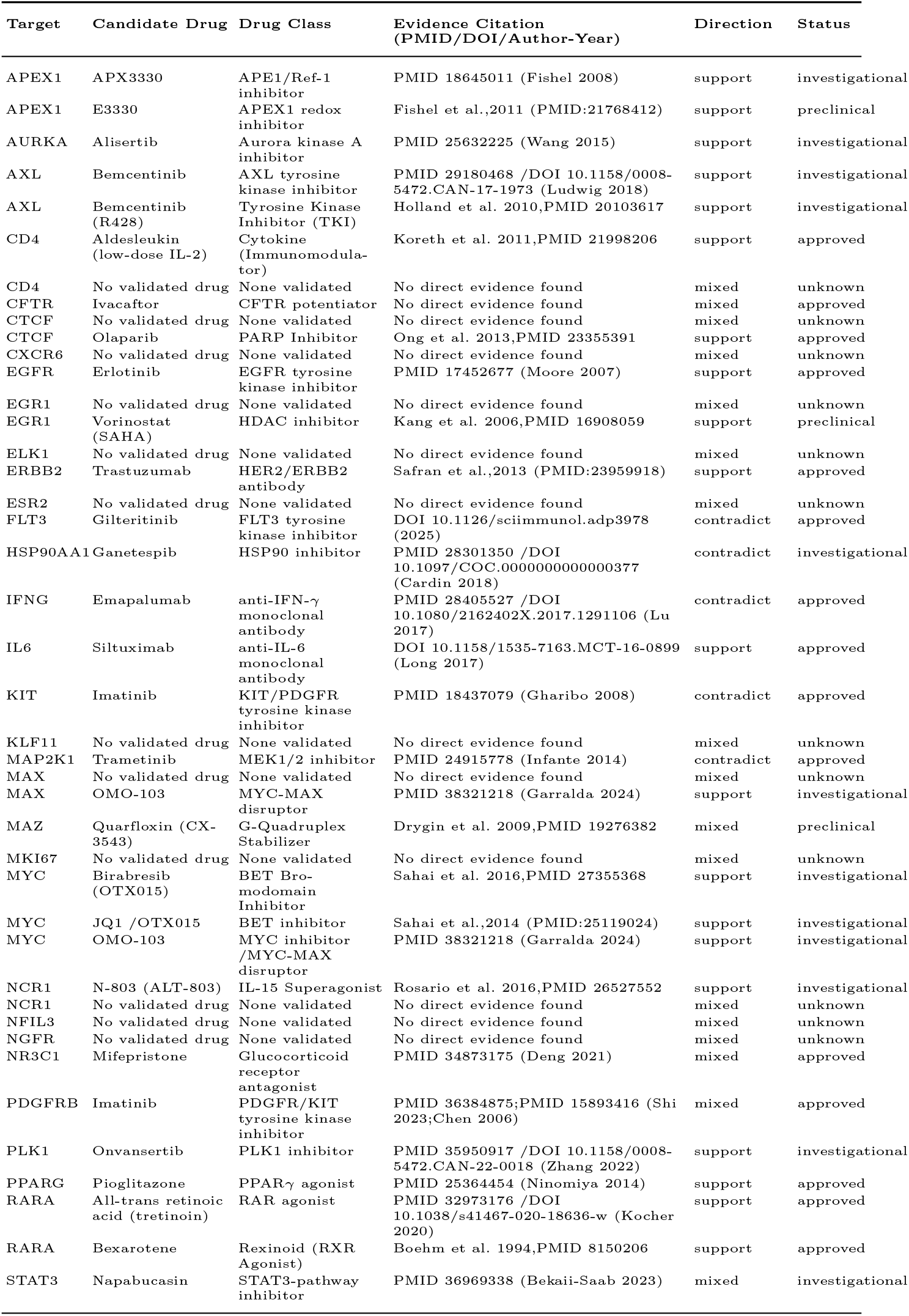
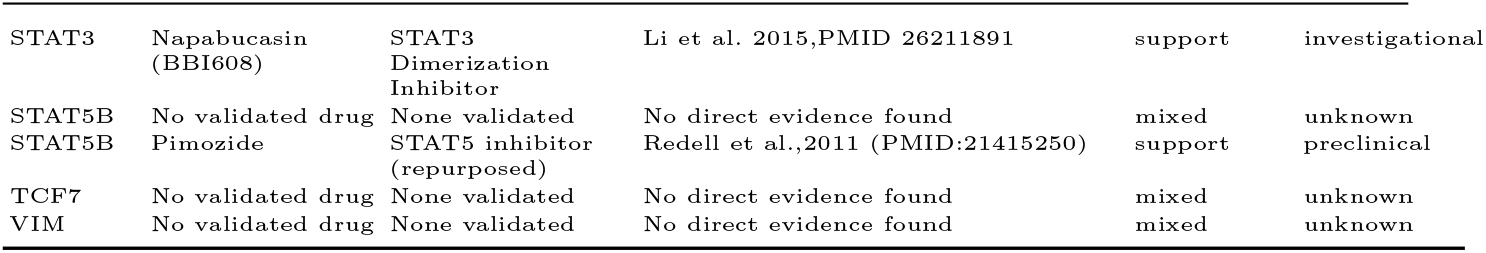
PDAC target-drug evidence ledger (deduplicated)

**Table S6:**
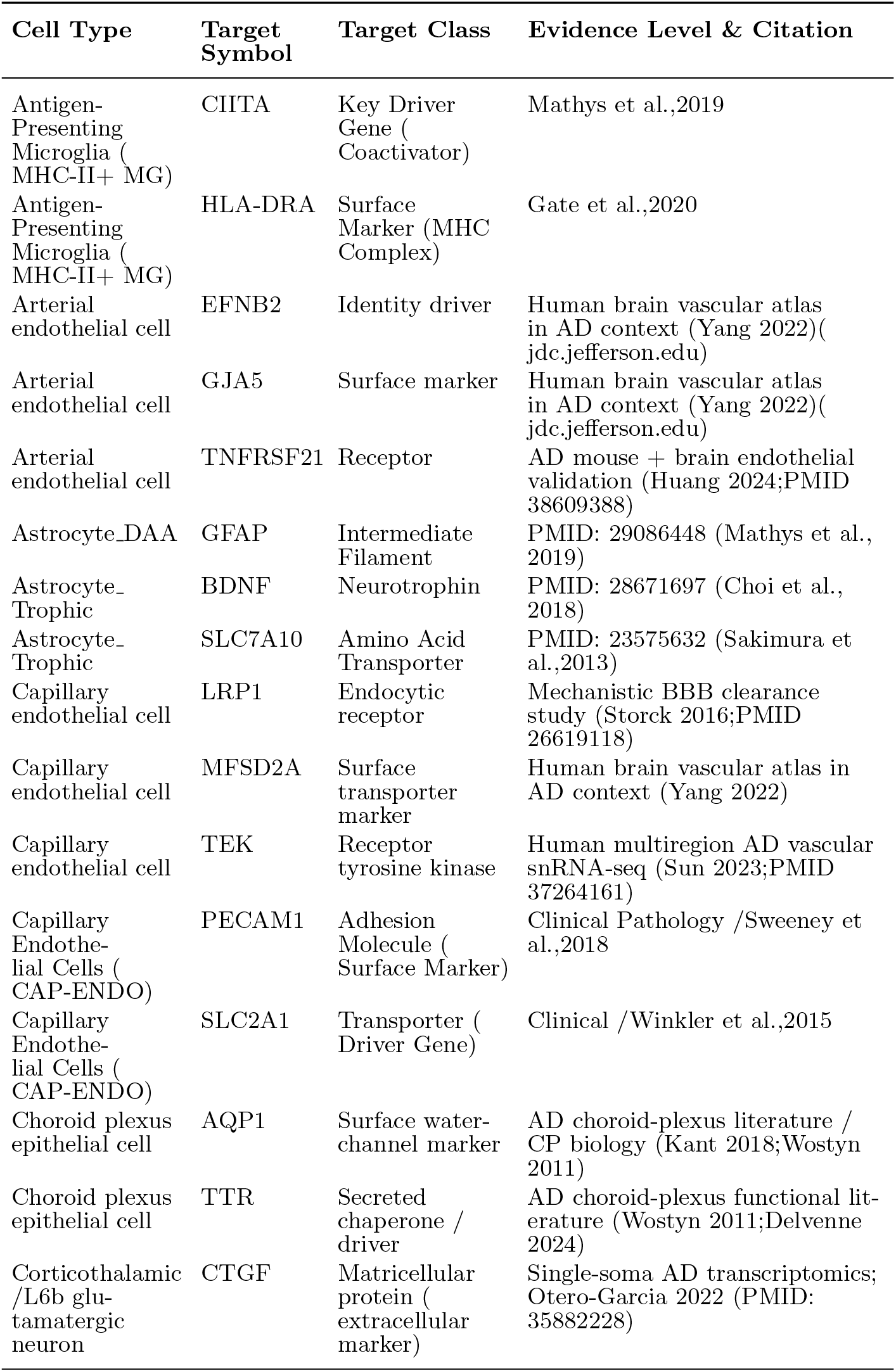
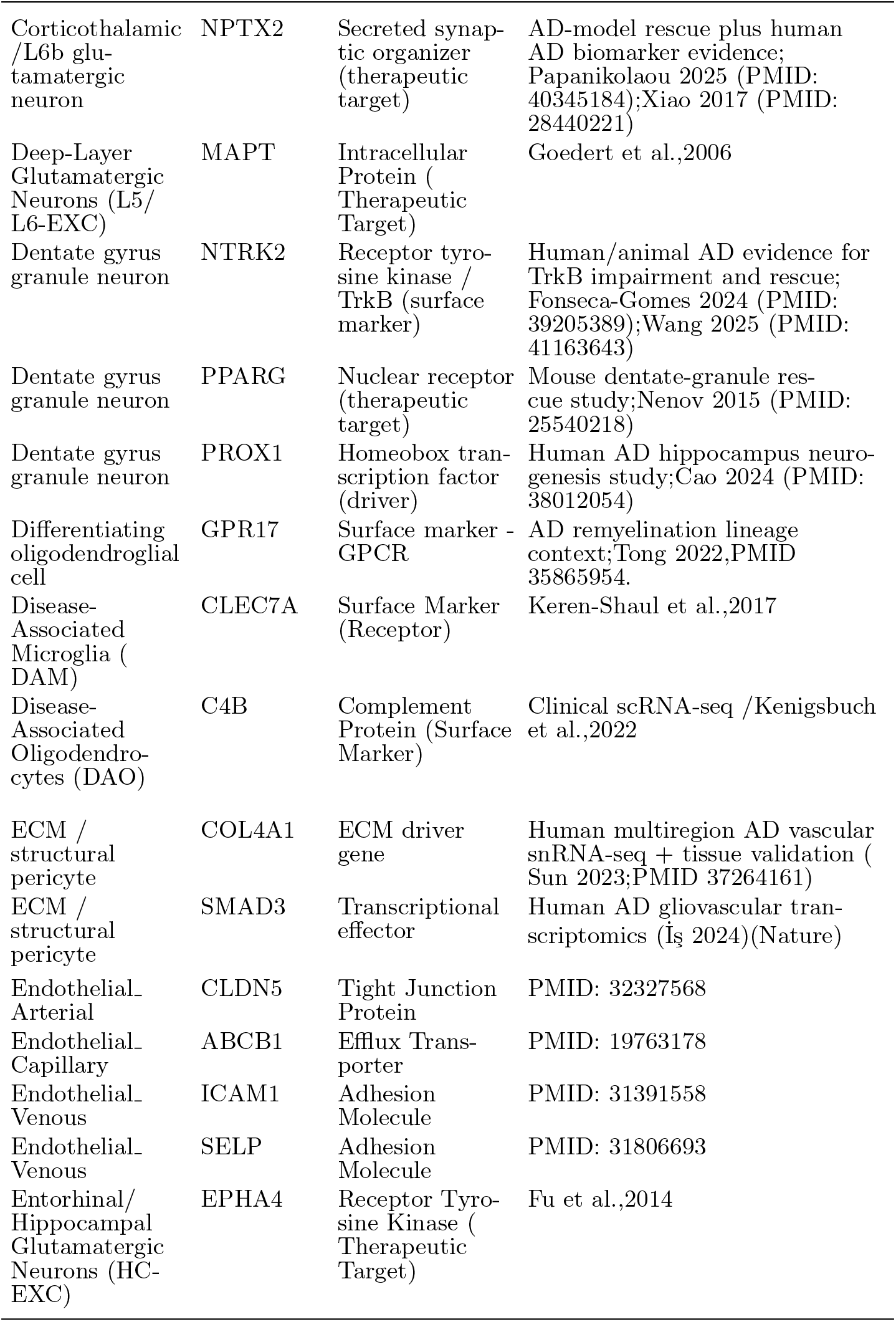
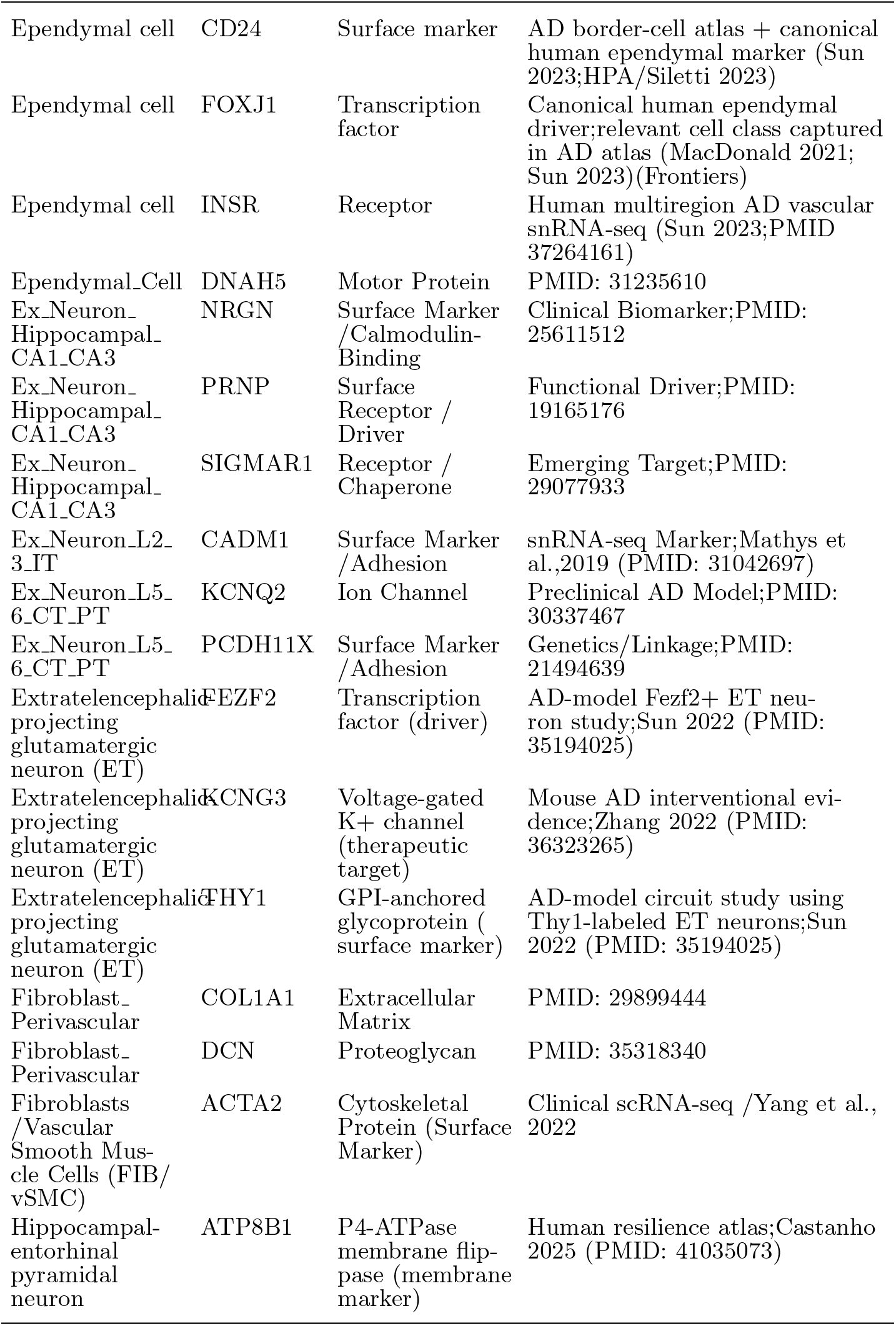
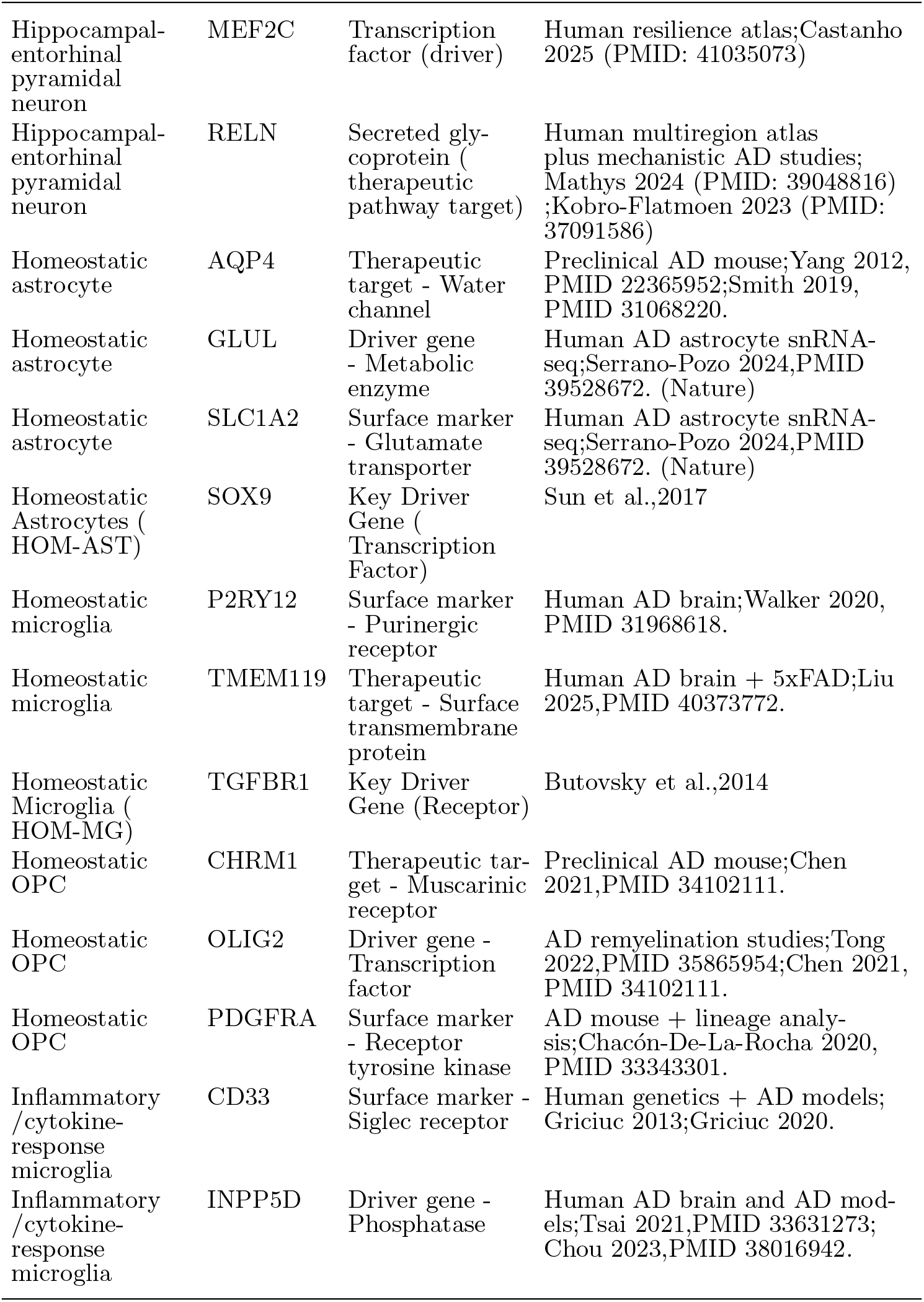
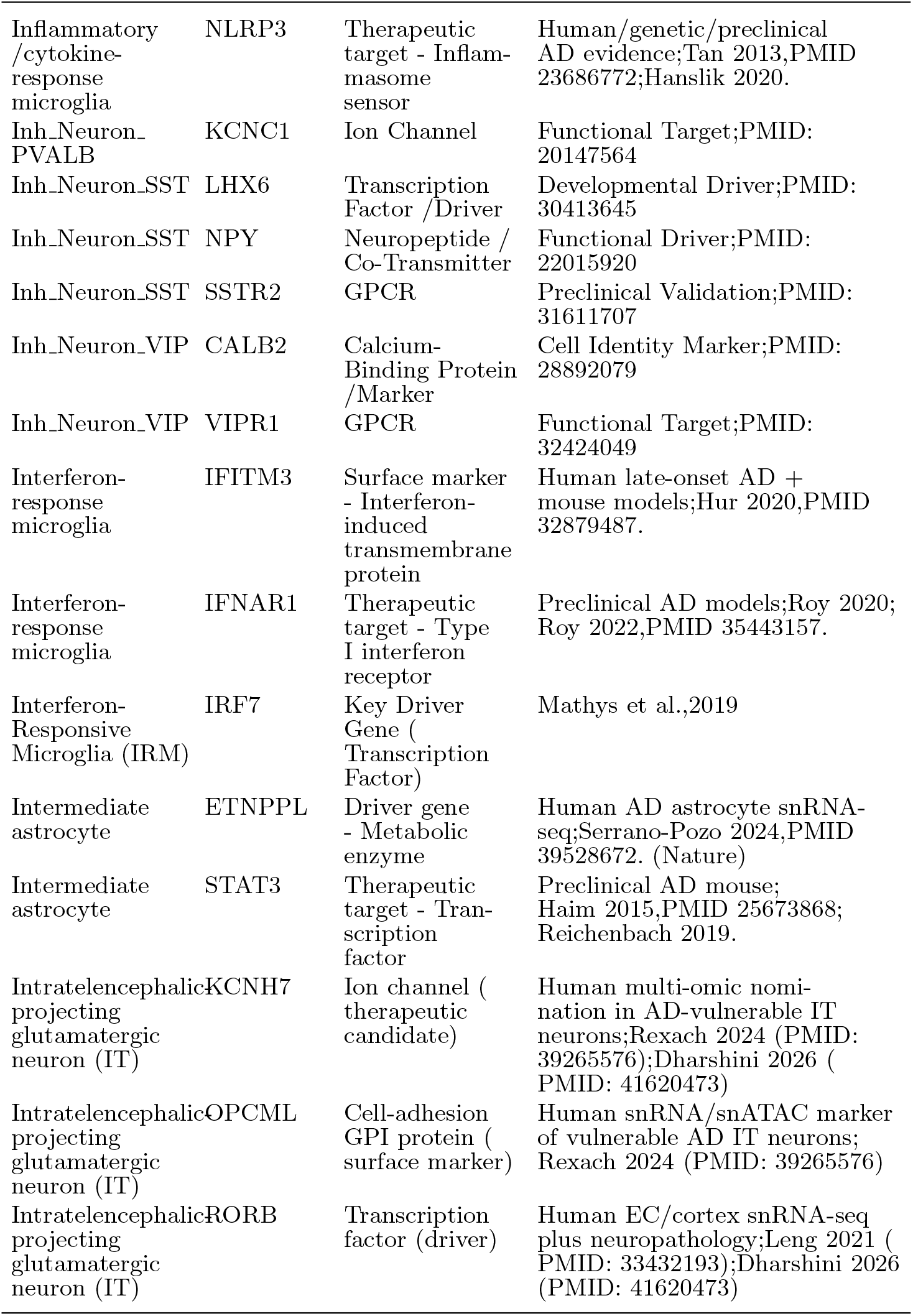
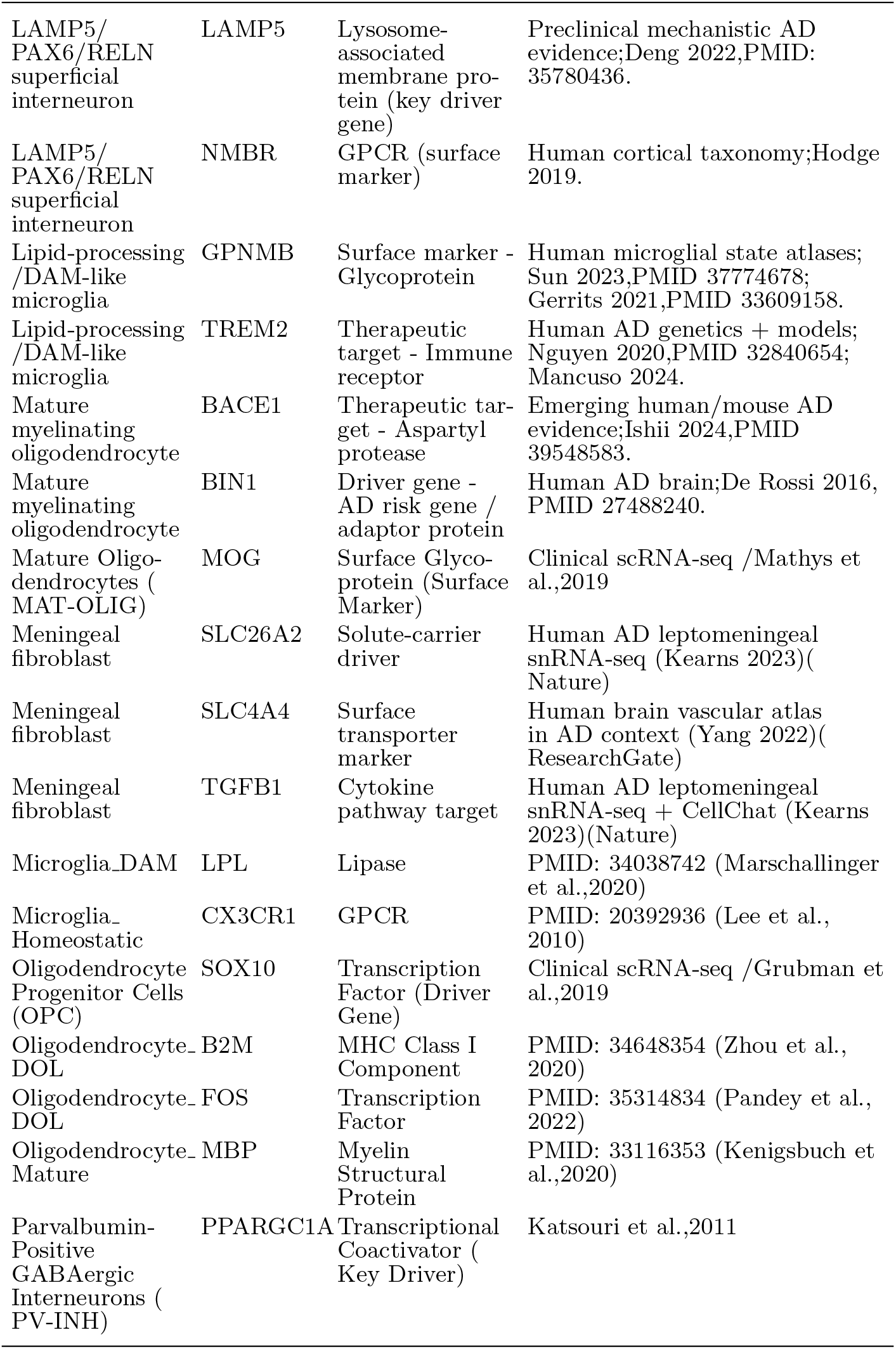
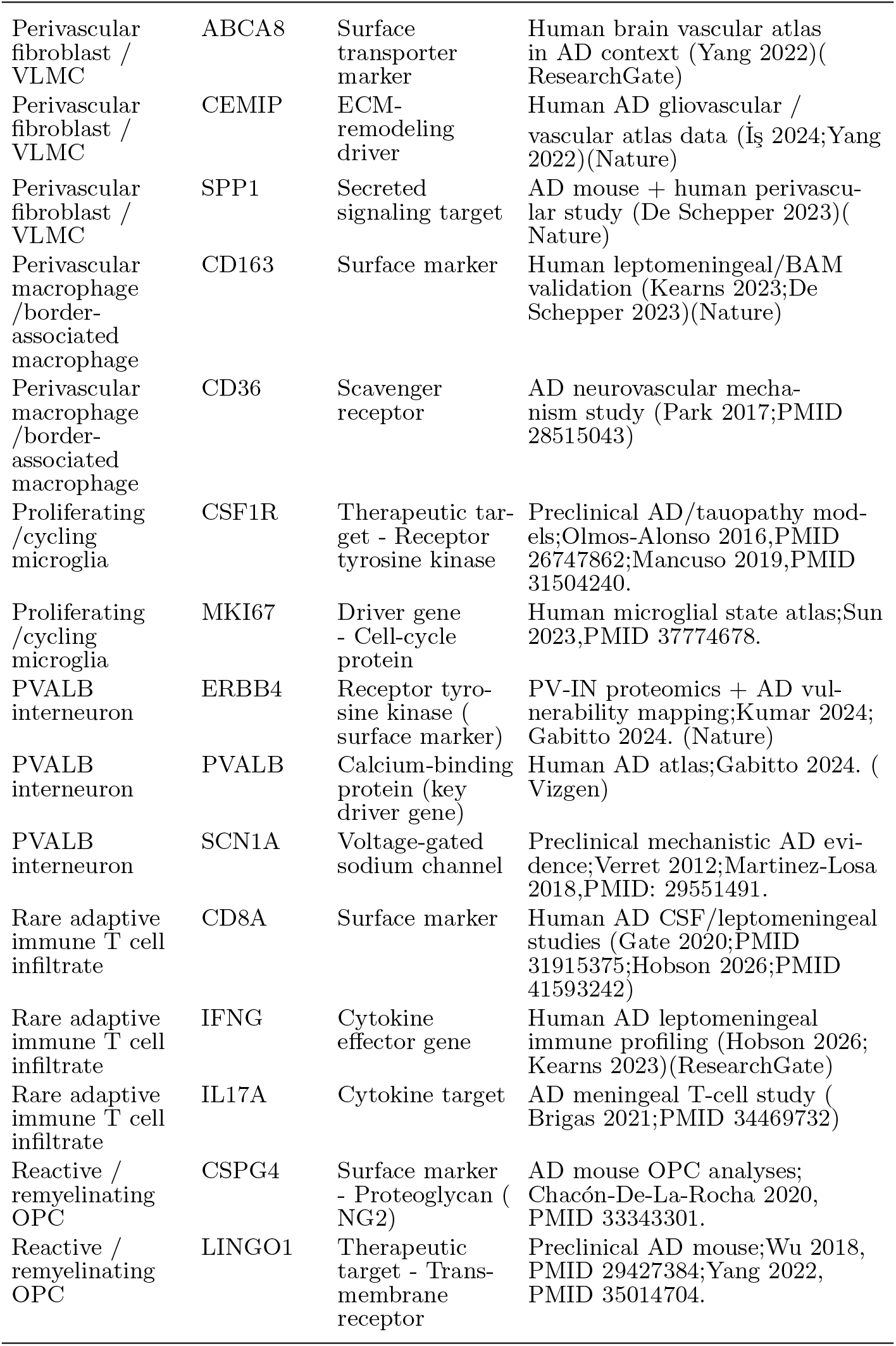
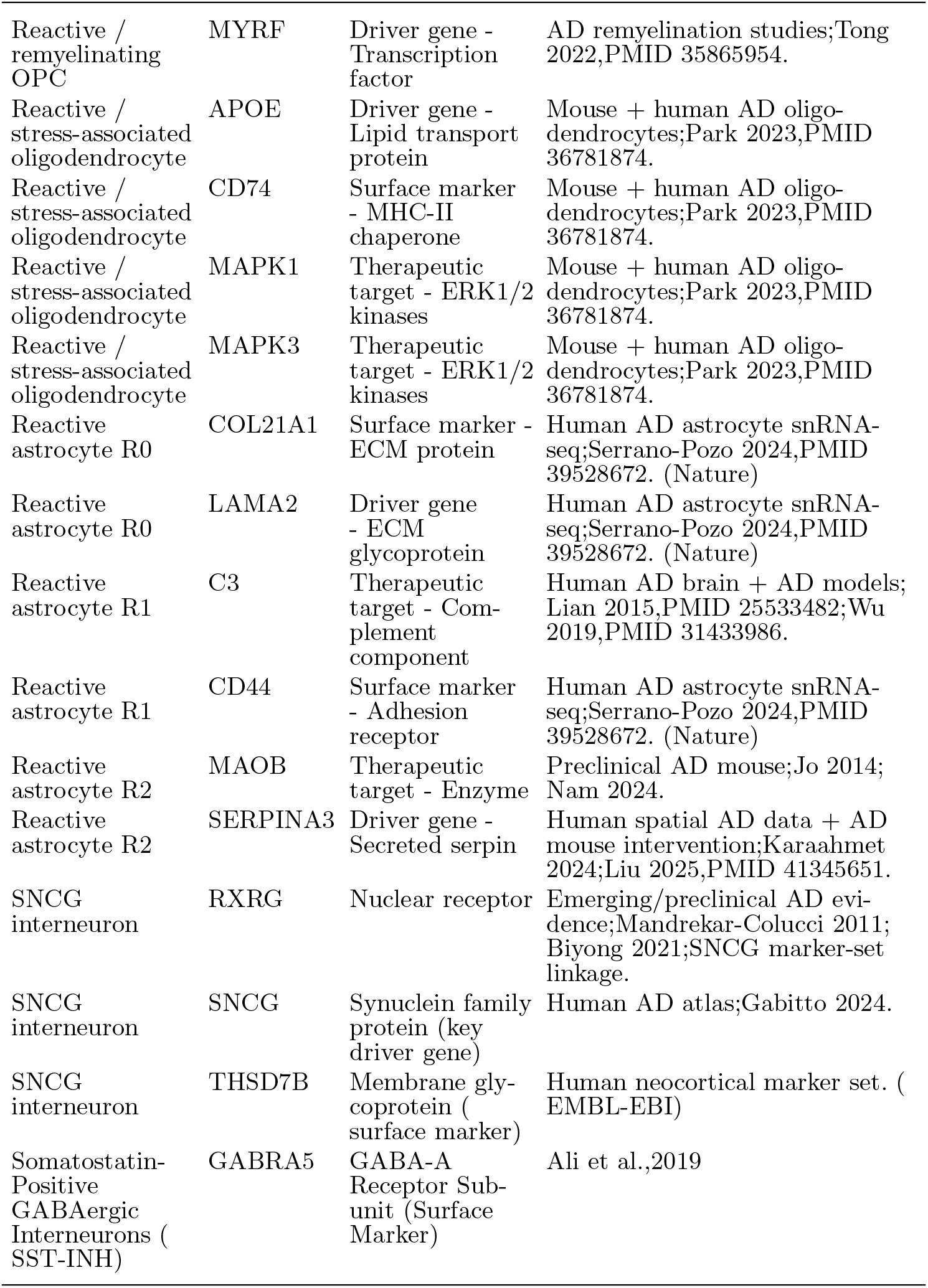
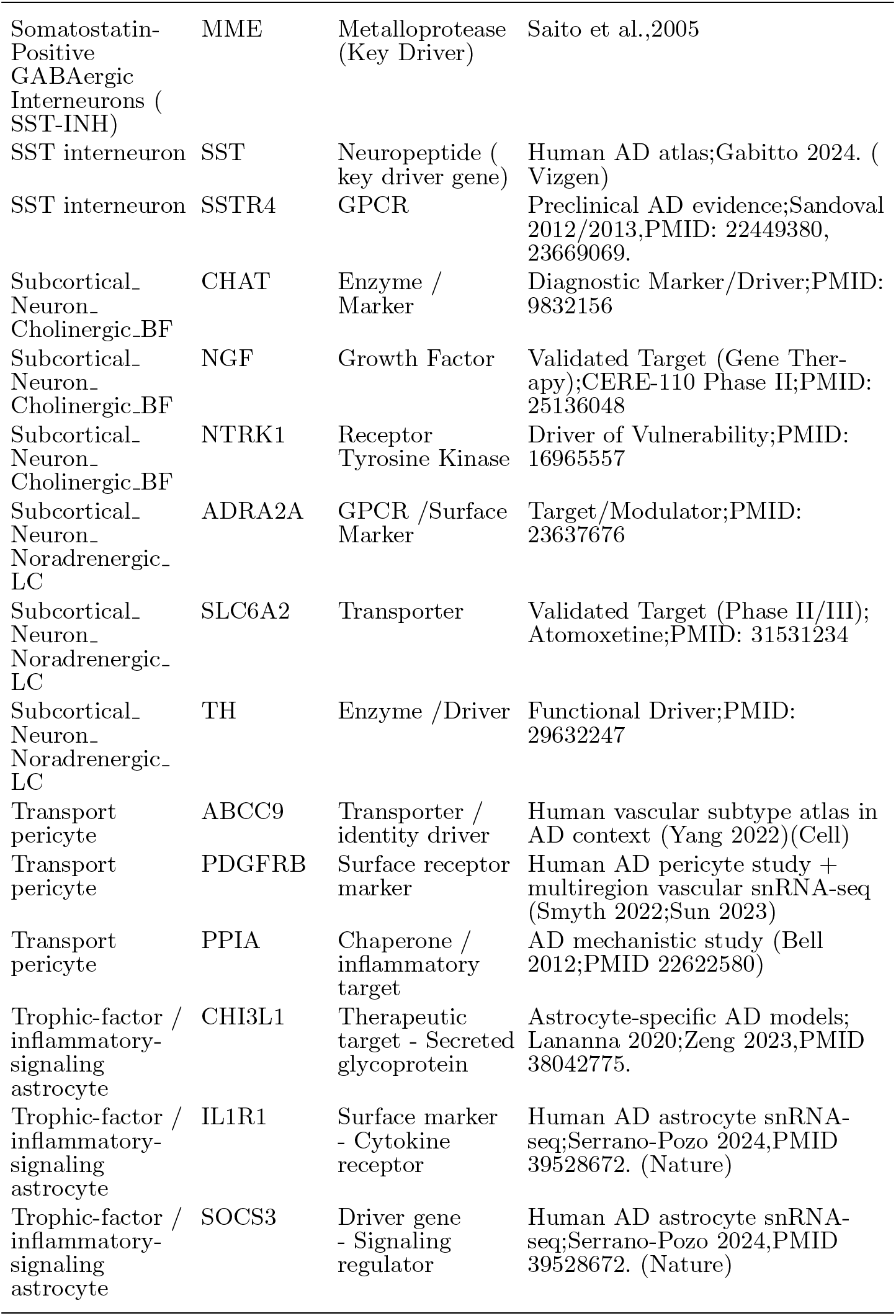
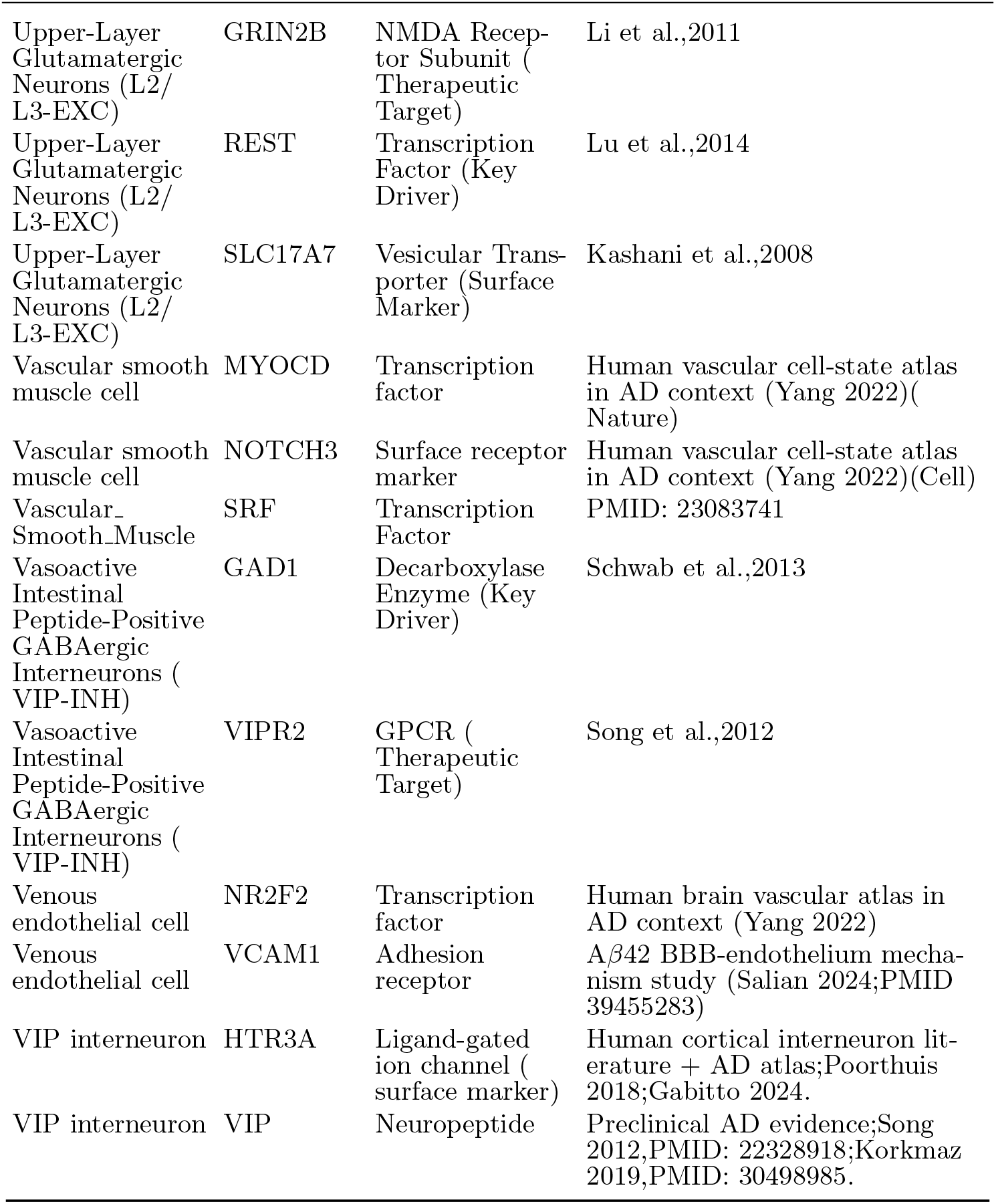
AD cell-specific targets.

**Table S7:**
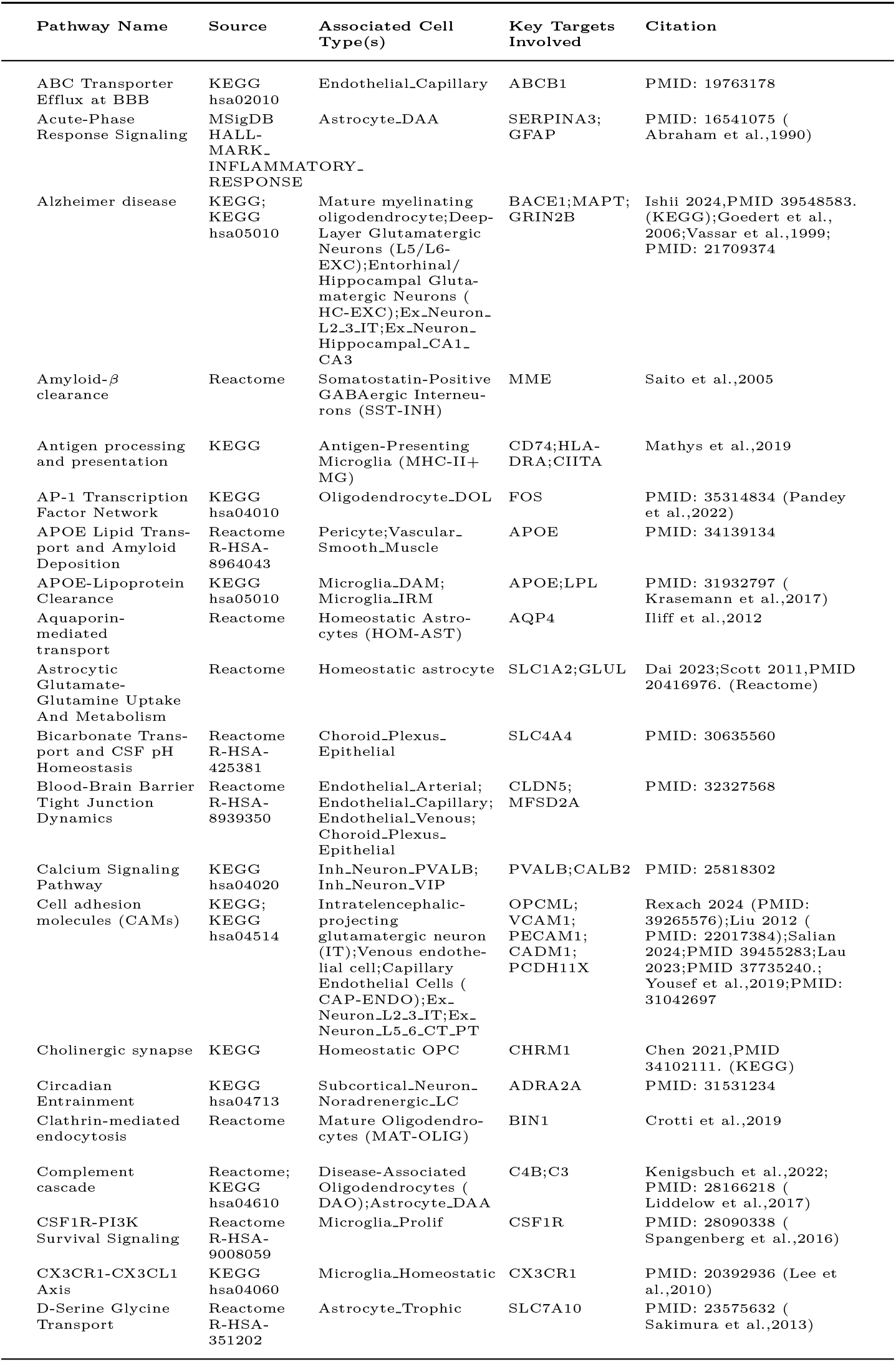
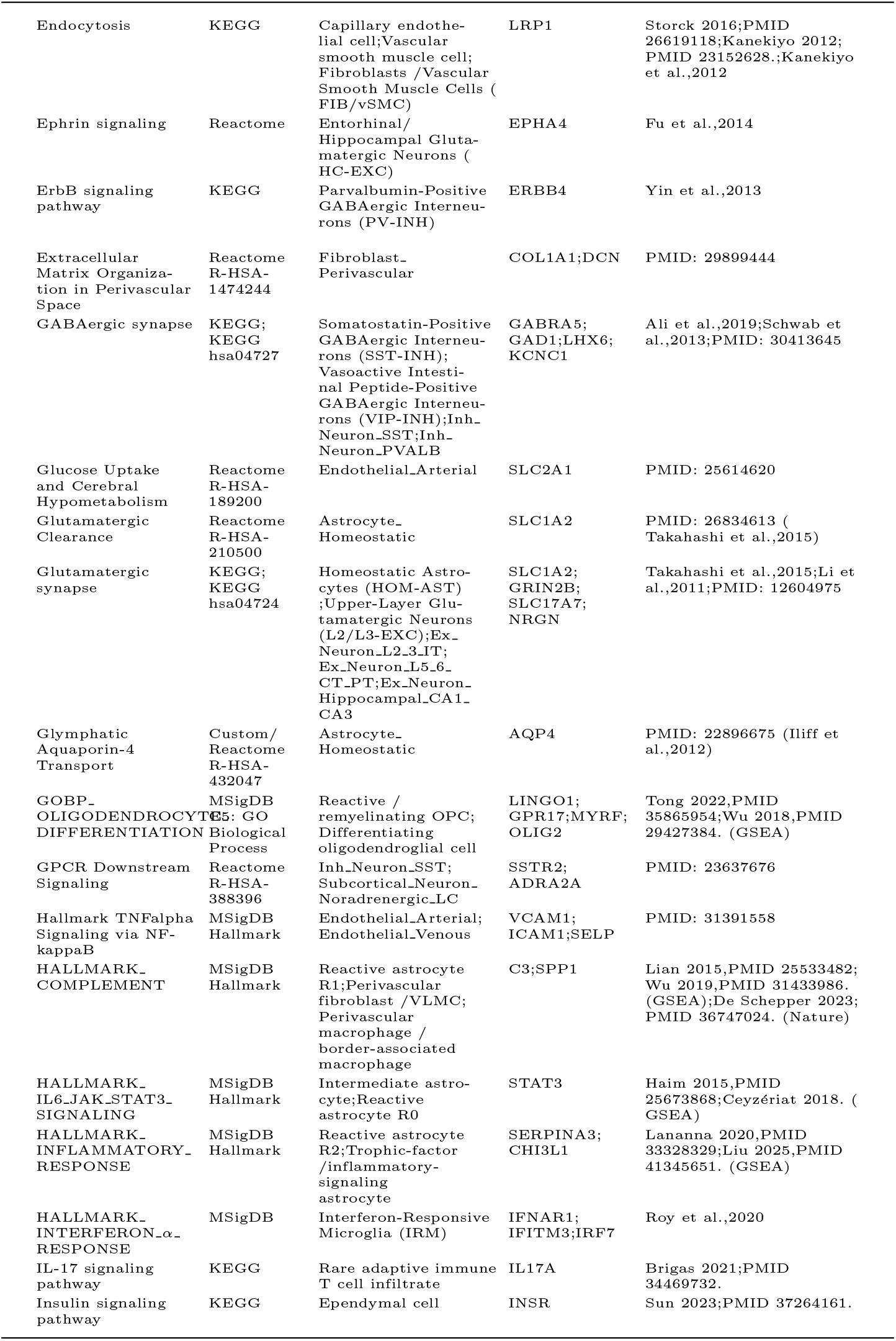
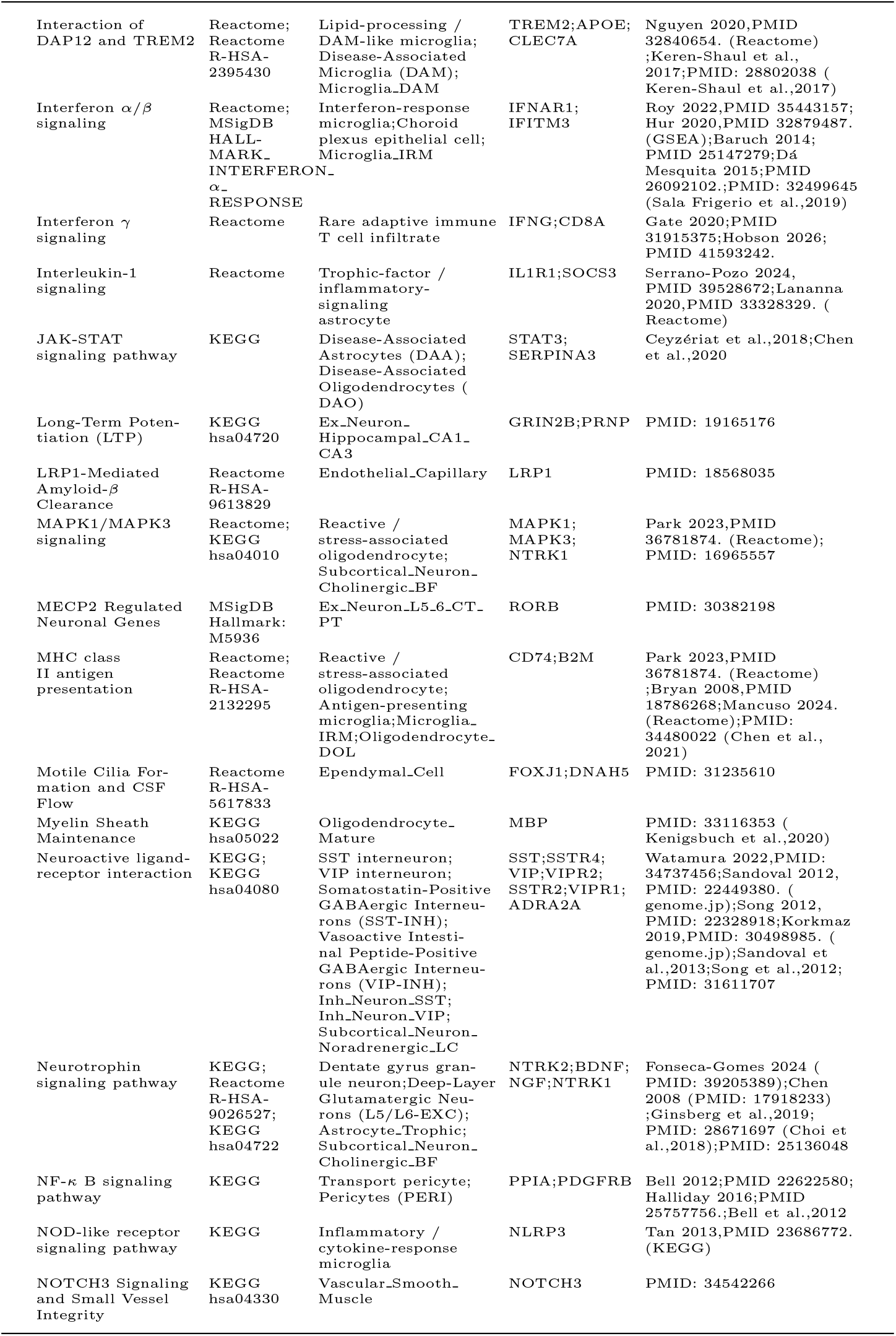
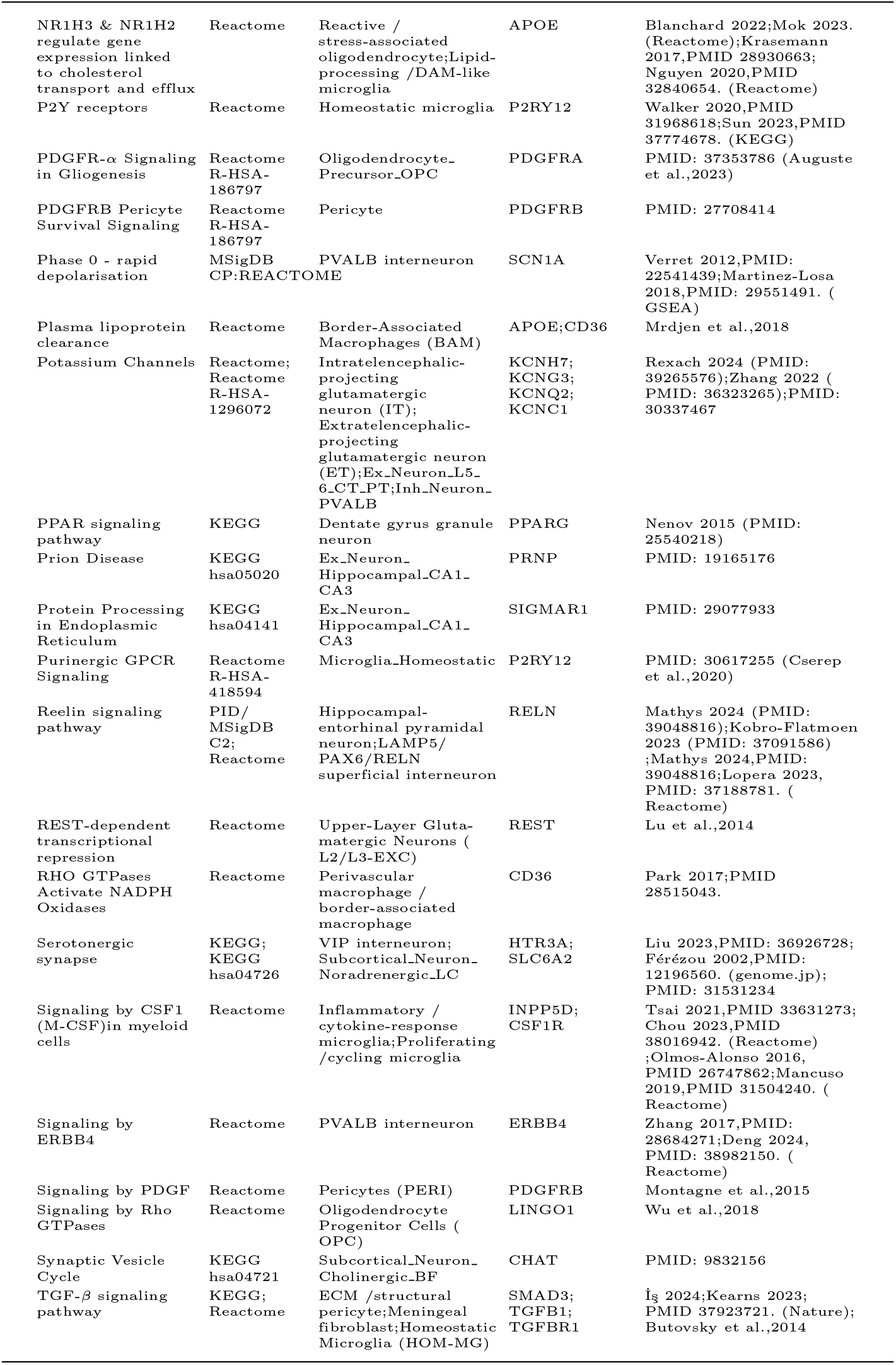
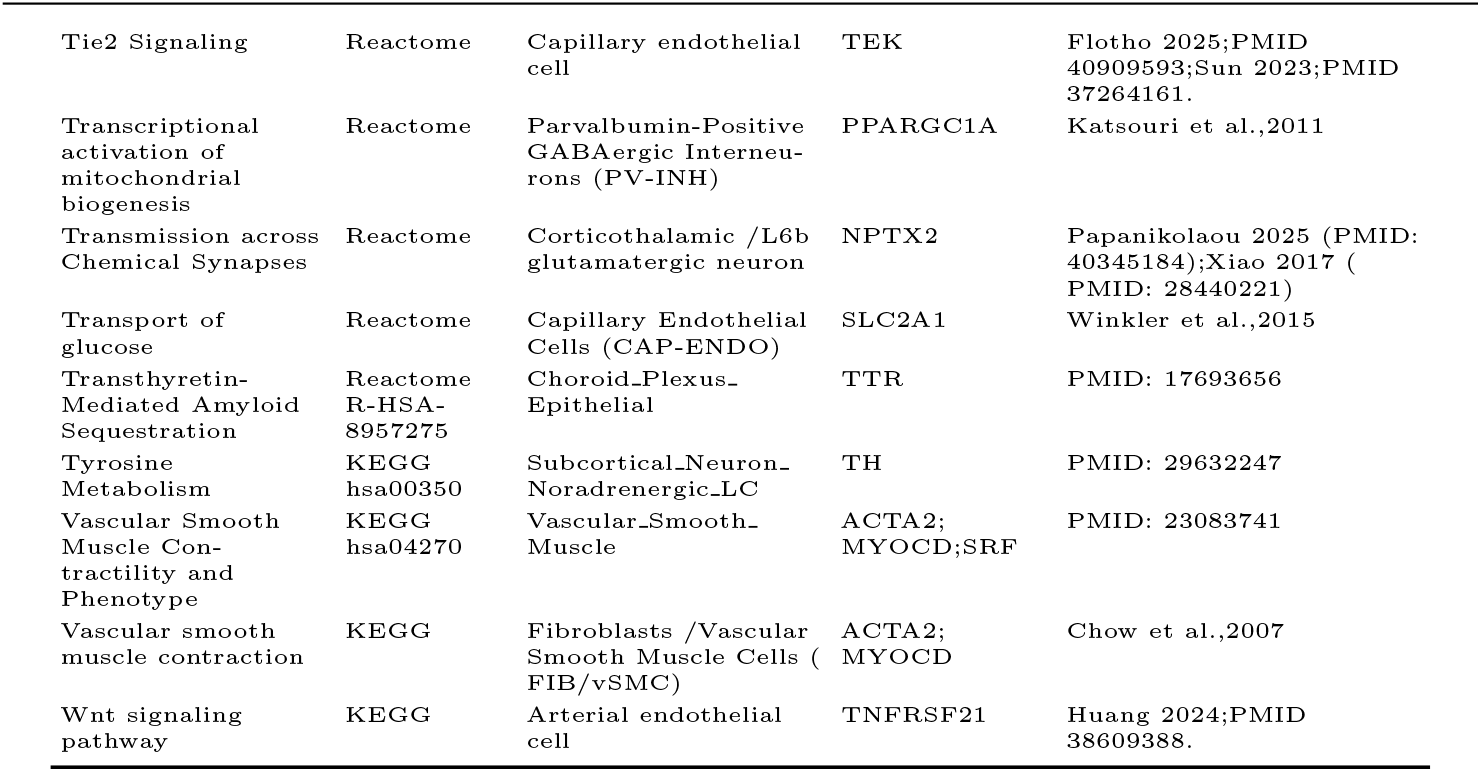
AD cell-specific pathways.

**Table S8:**
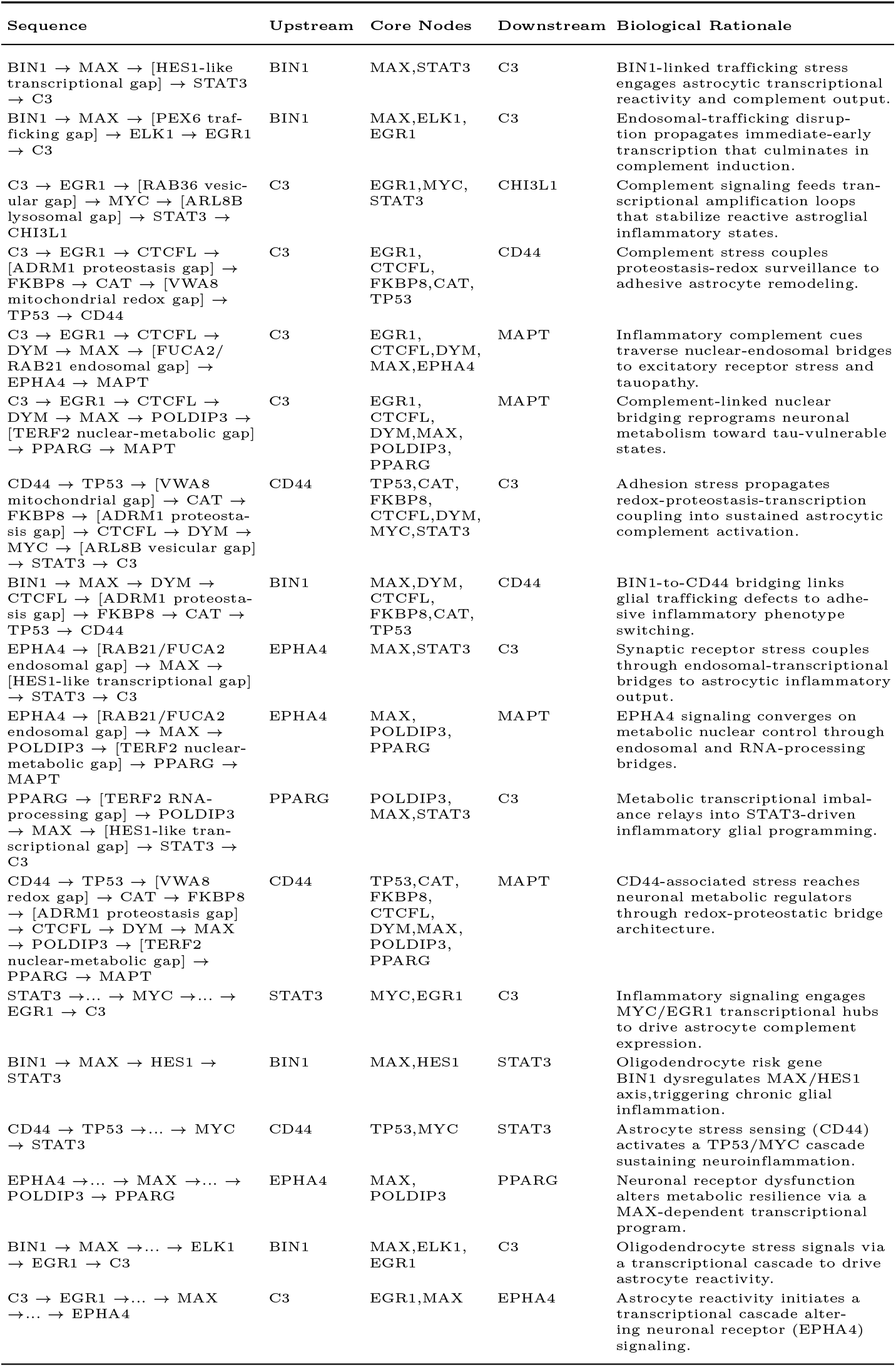
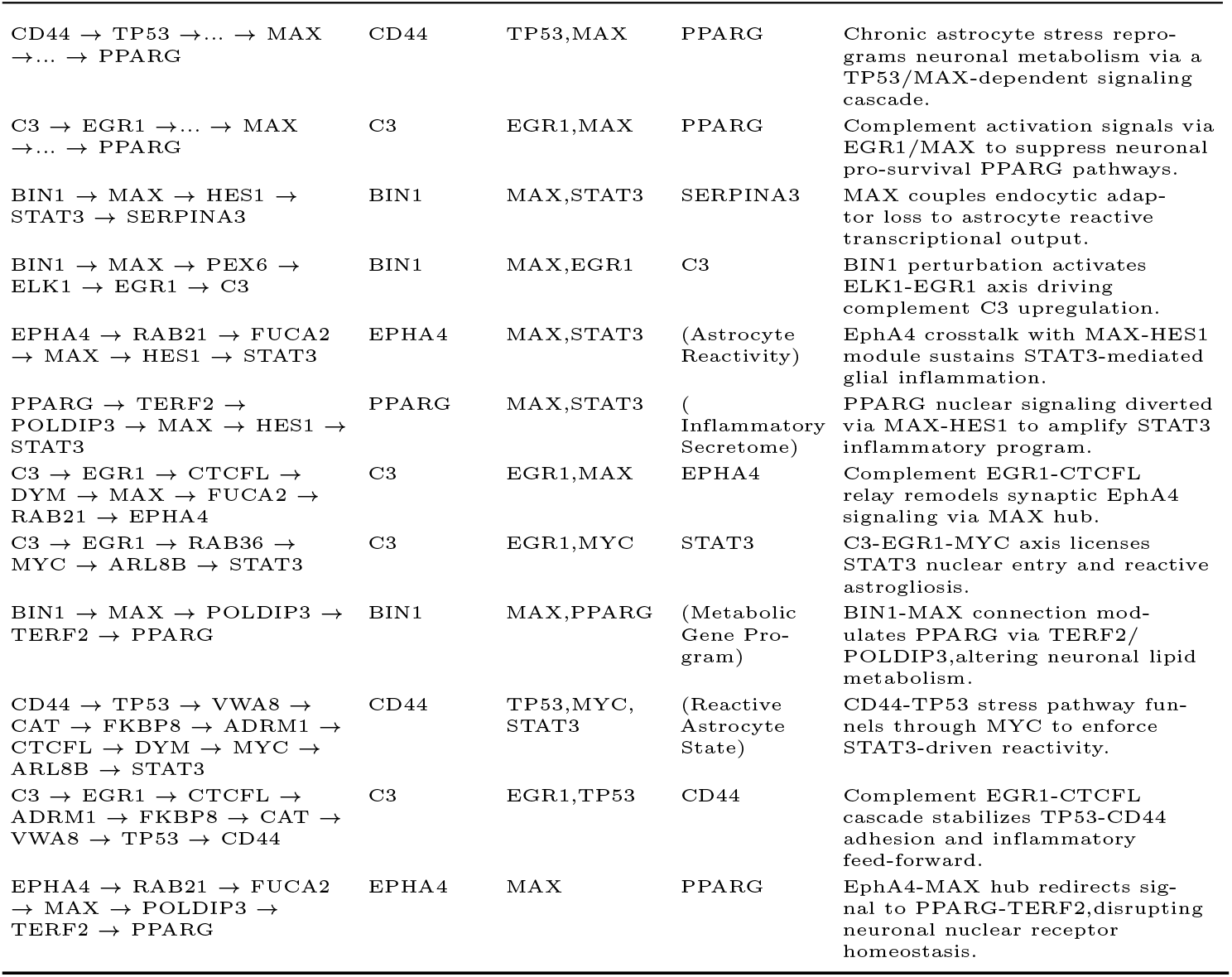
AD LLM-driven mechanism hypotheses.

| Sequence | Upstream | Core Nodes | Downstream | Biological Rationale |
| --- | --- | --- | --- | --- |
| BIN1 → MAX → [HES1-like transcriptional gap] → STAT3 → C3 | BIN1 | MAX,STAT3 | C3 | BIN1-linked trafficking stress engages astrocytic transcriptional reactivity and complement output. |
| BIN1 → MAX → [PEX6 trafficking gap] → ELK1 → EGR1 → C3 | BIN1 | MAX,ELK1, EGR1 | C3 | Endosomal-trafficking disruption propagates immediate-early transcription that culminates in complement induction. |
| C3 → EGR1 → [RAB36 vesicular gap] → MYC → [ARL8B lysosomal gap] → STAT3 → CHI3L1 | C3 | EGR1,MYC, STAT3 | CHI3L1 | Complement signaling feeds transcriptional amplification loops that stabilize reactive astroglial inflammatory states. |
| C3 → EGR1 → CTCFL → [ADRM1 proteostasis gap] → FKBP8 → CAT → [VWA8 mitochondrial redox gap] → TP53 → CD44 | C3 | EGR1, CTCFL, FKBP8,CAT, TP53 | CD44 | Complement stress couples proteostasis-redox surveillance to adhesive astrocyte remodeling. |
| C3 → EGR1 → CTCFL → DYM → MAX → [FUCA2/RAB21 endosomal gap] → EPHA4 → MAPT | C3 | EGR1, CTCFL,DYM, MAX,EPHA4 | MAPT | Inflammatory complement cues traverse nuclear-endosomal bridges to excitatory receptor stress and tauopathy. |
| C3 → EGR1 → CTCFL → DYM → MAX → POLDIP3 → [TERF2 nuclear-metabolic gap] → PPARG → MAPT | C3 | EGR1, CTCFL, DYM,MAX, POLDIP3, PPARG | MAPT | Complement-linked nuclear bridging reprograms neuronal metabolism toward tau-vulnerable states. |
| CD44 → TP53 → [VWA8 mitochondrial gap] → CAT → FKBP8 → [ADRM1 proteostasis gap] → CTCFL → DYM → MYC → [ARL8B vesicular gap] → STAT3 → C3 | CD44 | TP53,CAT, FKBP8, CTCFL,DYM, MYC,STAT3 | C3 | Adhesion stress propagates redox-proteostasis-transcription coupling into sustained astrocytic complement activation. |
| BIN1 → MAX → DYM → CTCFL → [ADRM1 proteostasis gap] → FKBP8 → CAT → TP53 → CD44 | BIN1 | MAX,DYM, CTCFL, FKBP8,CAT, TP53 | CD44 | BIN1-to-CD44 bridging links glial trafficking defects to adhesive inflammatory phenotype switching. |
| EPHA4 → [RAB21/FUCA2 endosomal gap] → MAX → [HES1-like transcriptional gap] → STAT3 → C3 | EPHA4 | MAX,STAT3 | C3 | Synaptic receptor stress couples through endosomal-transcriptional bridges to astrocytic inflammatory output. |
| EPHA4 → [RAB21/FUCA2 endosomal gap] → MAX → POLDIP3 → [TERF2 nuclear-metabolic gap] → PPARG → MAPT | EPHA4 | MAX, POLDIP3, PPARG | MAPT | EPHA4 signaling converges on metabolic nuclear control through endosomal and RNA-processing bridges. |
| PPARG → [TERF2 RNA-processing gap] → POLDIP3 → MAX → [HES1-like transcriptional gap] → STAT3 → C3 | PPARG | POLDIP3, MAX,STAT3 | C3 | Metabolic transcriptional imbalance relays into STAT3-driven inflammatory glial programming. |
| CD44 → TP53 → [VWA8 redox gap] → CAT → FKBP8 → [ADRM1 proteostasis gap] → CTCFL → DYM → MAX → POLDIP3 → [TERF2 nuclear-metabolic gap] → PPARG → MAPT | CD44 | TP53,CAT, FKBP8, CTCFL, DYM,MAX, POLDIP3, PPARG | MAPT | CD44-associated stress reaches neuronal metabolic regulators through redox-proteostatic bridge architecture. |
| STAT3 →... → MYC →... → EGR1 → C3 | STAT3 | MYC,EGR1 | C3 | Inflammatory signaling engages MYC/EGR1 transcriptional hubs to drive astrocyte complement expression. |
| BIN1 → MAX → HES1 → STAT3 | BIN1 | MAX,HES1 | STAT3 | Oligodendrocyte risk gene BIN1 dysregulates MAX/HES1 axis,triggering chronic glial inflammation. |
| CD44 → TP53 →... → MYC → STAT3 | CD44 | TP53,MYC | STAT3 | Astrocyte stress sensing (CD44) activates a TP53/MYC cascade sustaining neuroinflammation. |
| EPHA4 →... → MAX →... → POLDIP3 → PPARG | EPHA4 | MAX, POLDIP3 | PPARG | Neuronal receptor dysfunction alters metabolic resilience via a MAX-dependent transcriptional program. |
| BIN1 → MAX →... → ELK1 → EGR1 → C3 | BIN1 | MAX,ELK1, EGR1 | C3 | Oligodendrocyte stress signals via a transcriptional cascade to drive astrocyte reactivity. |
| C3 → EGR1 →... → MAX →... → EPHA4 | C3 | EGR1,MAX | EPHA4 | Astrocyte reactivity initiates a transcriptional cascade altering neuronal receptor (EPHA4) signaling. |
| CD44 → TP53 → ... → MAX<br>→ ... → PPARG | CD44 | TP53,MAX | PPARG | Chronic astrocyte stress reprograms neuronal metabolism via a TP53/MAX-dependent signaling cascade. |
| C3 → EGR1 → ... → MAX<br>→ ... → PPARG | C3 | EGR1,MAX | PPARG | Complement activation signals via EGR1/MAX to suppress neuronal pro-survival PPARG pathways. |
| BIN1 → MAX → HES1 →<br>STAT3 → SERPINA3 | BIN1 | MAX,STAT3 | SERPINA3 | MAX couples endocytic adaptor loss to astrocyte reactive transcriptional output. |
| BIN1 → MAX → PEX6 →<br>ELK1 → EGR1 → C3 | BIN1 | MAX,EGR1 | C3 | BIN1 perturbation activates ELK1-EGR1 axis driving complement C3 upregulation. |
| EPHA4 → RAB21 → FUCA2<br>→ MAX → HES1 → STAT3 | EPHA4 | MAX,STAT3 | (Astrocyte Reactivity) | EphA4 crosstalk with MAX-HES1 module sustains STAT3-mediated glial inflammation. |
| PPARG → TERF2 →<br>POLDIP3 → MAX → HES1 →<br>STAT3 | PPARG | MAX,STAT3 | (<br>Inflammatory Secretome) | PPARG nuclear signaling diverted via MAX-HES1 to amplify STAT3 inflammatory program. |
| C3 → EGR1 → CTCFL →<br>DYM → MAX → FUCA2 →<br>RAB21 → EPHA4 | C3 | EGR1,MAX | EPHA4 | Complement EGR1-CTCFL relay remodels synaptic EphA4 signaling via MAX hub. |
| C3 → EGR1 → RAB36 →<br>MYC → ARL8B → STAT3 | C3 | EGR1,MYC | STAT3 | C3-EGR1-MYC axis licenses STAT3 nuclear entry and reactive astrogliosis. |
| BIN1 → MAX → POLDIP3 →<br>TERF2 → PPARG | BIN1 | MAX,PPARG | (Metabolic Gene Program) | BIN1-MAX connection modulates PPARG via TERF2/POLDIP3,altering neuronal lipid metabolism. |
| CD44 → TP53 → VWA8 →<br>CAT → FKBP8 → ADRM1 →<br>CTCFL → DYM → MYC →<br>ARL8B → STAT3 | CD44 | TP53,MYC,<br>STAT3 | (Reactive Astrocyte State) | CD44-TP53 stress pathway funnels through MYC to enforce STAT3-driven reactivity. |
| C3 → EGR1 → CTCFL →<br>ADRM1 → FKBP8 → CAT →<br>VWA8 → TP53 → CD44 | C3 | EGR1,TP53 | CD44 | Complement EGR1-CTCFL cascade stabilizes TP53-CD44 adhesion and inflammatory feed-forward. |
| EPHA4 → RAB21 → FUCA2<br>→ MAX → POLDIP3 →<br>TERF2 → PPARG | EPHA4 | MAX | PPARG | EphA4-MAX hub redirects signal to PPARG-TERF2,disrupting neuronal nuclear receptor homeostasis. |

**Table S9:**
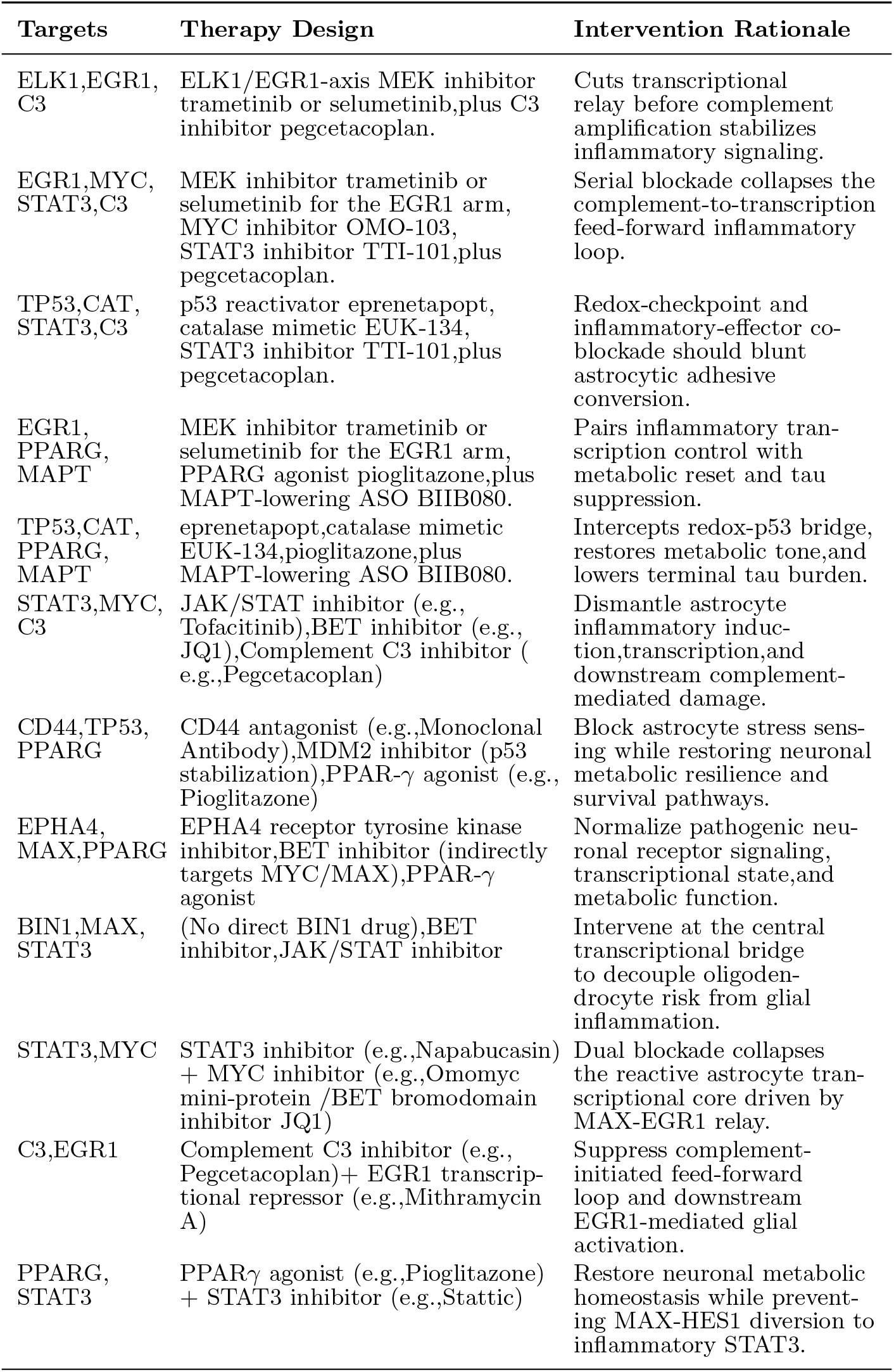
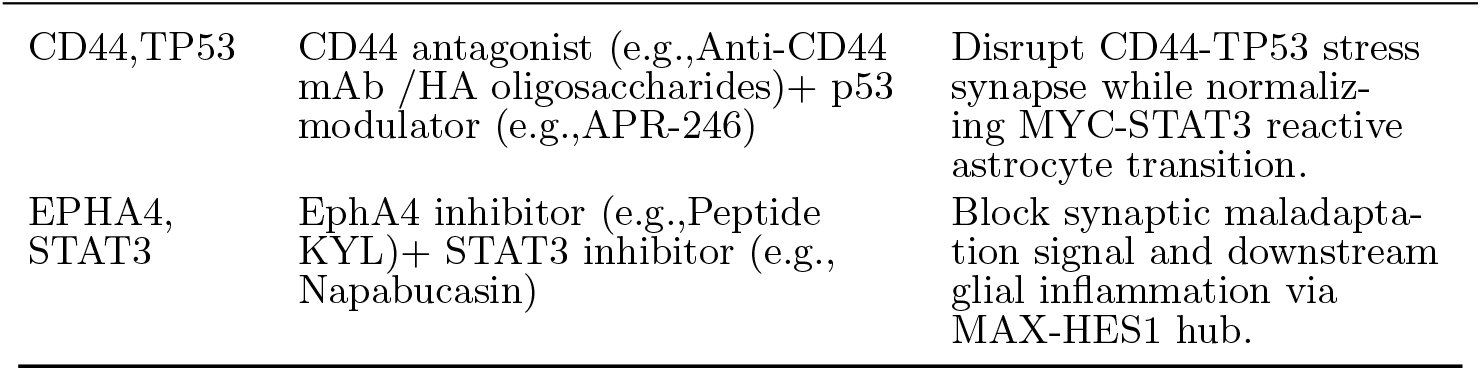
AD upgraded therapy combinations.

| Targets | Therapy Design | Intervention Rationale |
| --- | --- | --- |
| ELK1,EGR1, C3 | ELK1/EGR1-axis MEK inhibitor trametinib or selumetinib,plus C3 inhibitor pegcetacoplan. | Cuts transcriptional relay before complement amplification stabilizes inflammatory signaling. |
| EGR1,MYC, STAT3,C3 | MEK inhibitor trametinib or selumetinib for the EGR1 arm, MYC inhibitor OMO-103, STAT3 inhibitor TTI-101,plus pegcetacoplan. | Serial blockade collapses the complement-to-transcription feed-forward inflammatory loop. |
| TP53,CAT, STAT3,C3 | p53 reactivator eprenetapopt, catalase mimetic EUK-134, STAT3 inhibitor TTI-101,plus pegcetacoplan. | Redox-checkpoint and inflammatory-effector co-blockade should blunt astrocytic adhesive conversion. |
| EGR1, PPARG, MAPT | MEK inhibitor trametinib or selumetinib for the EGR1 arm, PPARG agonist pioglitazone,plus MAPT-lowering ASO BIIB080. | Pairs inflammatory transcription control with metabolic reset and tau suppression. |
| TP53,CAT, PPARG, MAPT | eprenetapopt,catalase mimetic EUK-134,pioglitazone,plus MAPT-lowering ASO BIIB080. | Intercepts redox-p53 bridge, restores metabolic tone,and lowers terminal tau burden. |
| STAT3,MYC, C3 | JAK/STAT inhibitor (e.g., Tofacitinib),BET inhibitor (e.g., JQ1),Complement C3 inhibitor ( e.g.,Pegcetacoplan) | Dismantle astrocyte inflammatory induction,transcription,and downstream complement-mediated damage. |
| CD44,TP53, PPARG | CD44 antagonist (e.g.,Monoclonal Antibody),MDM2 inhibitor (p53 stabilization),PPAR-γ agonist (e.g., Pioglitazone) | Block astrocyte stress sensing while restoring neuronal metabolic resilience and survival pathways. |
| EPHA4, MAX,PPARG | EPHA4 receptor tyrosine kinase inhibitor,BET inhibitor (indirectly targets MYC/MAX),PPAR-γ agonist | Normalize pathogenic neuronal receptor signaling, transcriptional state,and metabolic function. |
| BIN1,MAX, STAT3 | (No direct BIN1 drug),BET inhibitor,JAK/STAT inhibitor | Intervene at the central transcriptional bridge to decouple oligodendrocyte risk from glial inflammation. |
| STAT3,MYC | STAT3 inhibitor (e.g.,Napabucasin) + MYC inhibitor (e.g.,Omomyc mini-protein /BET bromodomain inhibitor JQ1) | Dual blockade collapses the reactive astrocyte transcriptional core driven by MAX-EGR1 relay. |
| C3,EGR1 | Complement C3 inhibitor (e.g., Pegcetacoplan)+ EGR1 transcriptional repressor (e.g.,Mithramycin A) | Suppress complement-initiated feed-forward loop and downstream EGR1-mediated glial activation. |
| PPARG, STAT3 | PPARγ agonist (e.g.,Pioglitazone) + STAT3 inhibitor (e.g.,Stattic) | Restore neuronal metabolic homeostasis while preventing MAX-HES1 diversion to inflammatory STAT3. |
| CD44,TP53 | CD44 antagonist (e.g.,Anti-CD44 mAb /HA oligosaccharides)+ p53 modulator (e.g.,APR-246) | Disrupt CD44-TP53 stress synapse while normalizing MYC-STAT3 reactive astrocyte transition. |
| EPHA4, STAT3 | EphA4 inhibitor (e.g.,Peptide KYL)+ STAT3 inhibitor (e.g., Napabucasin) | Block synaptic maladaptation signal and downstream glial inflammation via MAX-HES1 hub. |

**Table S10:**
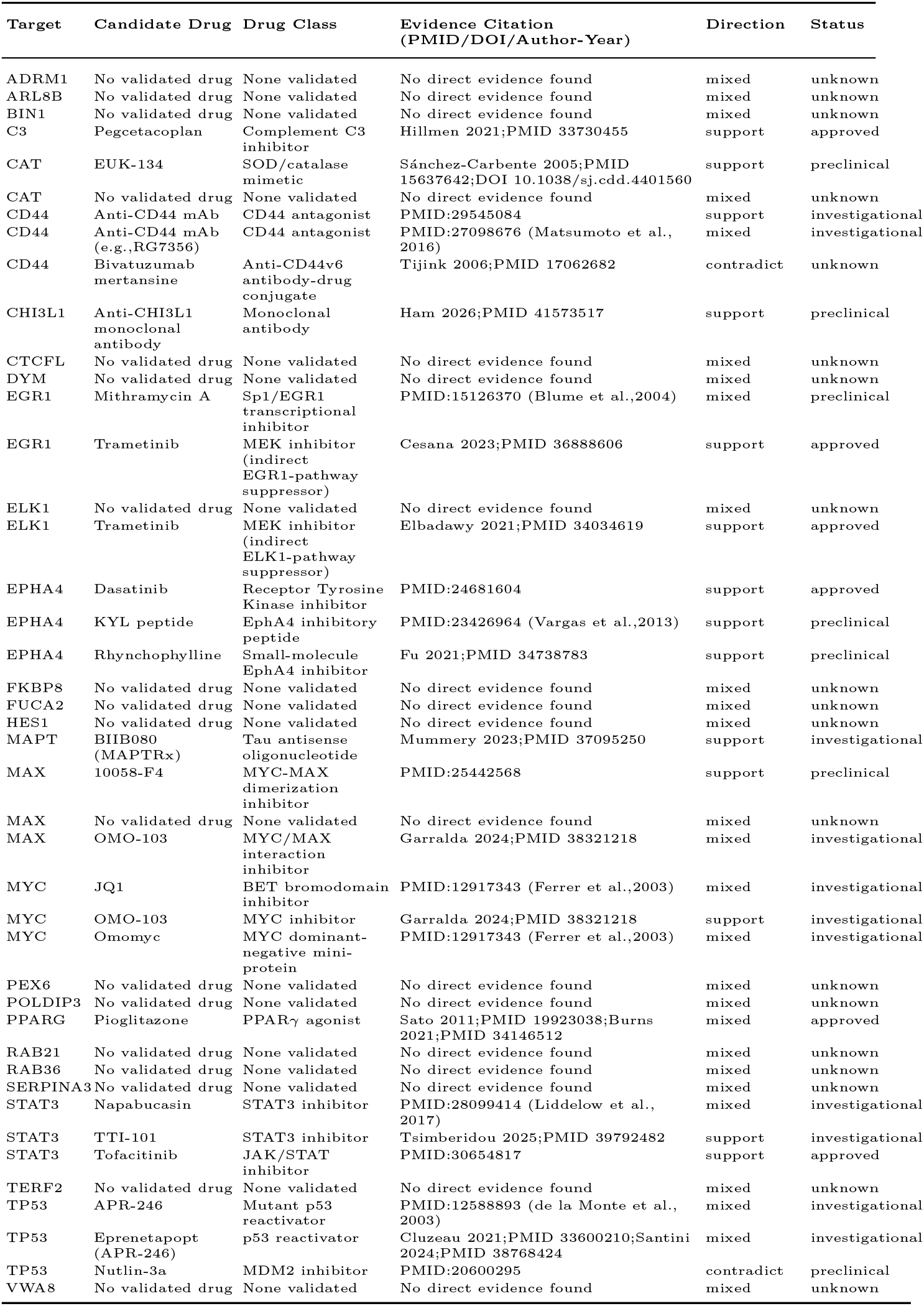
AD target-drug evidence ledger (deduplicated)

